# The *Drosophila* histone methyl-transferase SET1 coordinates multiple signaling pathways in regulating male germline stem cell maintenance and differentiation

**DOI:** 10.1101/2024.02.14.580277

**Authors:** Velinda Vidaurre, Annabelle Song, Taibo Li, Wai Lim Ku, Keji Zhao, Jiang Qian, Xin Chen

## Abstract

Many cell types come from tissue-specific adult stem cells that maintain the balance between proliferation and differentiation. Here, we study how the H3K4me3 methyltransferase, Set1, regulates early-stage male germ cell proliferation and differentiation in *Drosophila*. Early-stage germline-specific knockdown of *set1* results in a temporally progressed defects, arising as germ cell loss and developing to overpopulated early-stage germ cells. These germline defects also impact the niche architecture and cyst stem cell lineage in a non-cell-autonomous manner. Additionally, wild-type Set1, but not the catalytically inactive Set1, could rescue the *set1* knockdown phenotypes, highlighting the functional importance of the methyl-transferase activity of the Set1 enzyme. Further, RNA-seq experiments reveal key signaling pathway components, such as the JAK-STAT pathway gene *stat92E* and the BMP pathway gene *mad*, that are upregulated upon *set1* knockdown. Genetic interaction assays support the functional relationships between *set1* and JAK-STAT or BMP pathways, as mutations of both the *stat92E* and *mad* genes suppress the *set1* knockdown phenotypes. These findings enhance our understanding of the balance between proliferation and differentiation in an adult stem cell lineage. The germ cell loss followed by over-proliferation phenotypes when inhibiting a histone methyl-transferase raise concerns about using their inhibitors in cancer therapy.

## Introduction

In multicellular organisms, homeostasis and regeneration of many tissues largely depend on adult stem cells. These endogenous stem cells often undergo asymmetric cell divisions to allow both the stem cell to maintain its own population through self-renewal and the differentiation process to replace cells lost under physiological and pathological conditions (Kahney et al., 2019; Knoblich, 2008; Morrison and Spradling, 2008; Venkei and Yamashita, 2018). Disruption of these processes can potentially result in the mis-regulation of stem cell activity, leading to cancer or tissue degeneration (Clevers, 2005; Knoblich, 2010; Morrison and Kimble, 2006; Zion et al., 2020). There are two main modes of dysregulation in adult stem cell lineages that can result in unrestrained cell proliferation underlying tumorigenesis: Constraints of normal stem cell expansion may be compromised or inactivated. This situation could occur if the dependence of stem cells on the niche is disrupted, resulting in a niche-independent overpopulation of stem cells or enhancement of stem cell activities. Alternatively, the transit-amplifying cells could fail to exit proliferation and enter the terminal differentiation program (Clarke and Fuller, 2006; Mukherjee et al., 2015; Sell, 2010; Zhang and Hsu, 2017).

*Drosophila* spermatogenesis is an excellent model system to study stem cell proliferation and differentiation (Davies and Fuller, 2008; Fuller, 1993; Gleason et al., 2018). Spermatogenesis is initiated with the asymmetric division of the germline stem cells (GSCs) to produce a self-renewed GSC and a gonialblast. The gonialblast undergoes four rounds of transit-amplifying mitotic divisions as spermatoginal cells. After mitosis, the 16 spermatogonial cells enter meiosis with a prolonged G2-phase as the primary spermatocytes. In addition to the GSC-derived germline lineage, the testis also has at least two somatic cell populations: the hub cells and the cyst cells (de Cuevas and Matunis, 2011). The hub cells are post-mitotic and they support the GSCs and the cyst stem cells (CySCs) (Kiger et al., 2001; Leatherman, 2013; Leatherman and Dinardo, 2010; Losick et al., 2011; Tran et al., 2000; Yamashita et al., 2003). The CySCs give rise to the cyst cells that support the germ cells throughout spermatogenesis (Leatherman and Dinardo, 2008; Lim and Fuller, 2012). The CySC also undergoes an asymmetric cell division to produce a self-renewed CySC and a cyst cell, which never divides again (Cheng et al., 2011). Two cyst cells encapsulate the germ cells as they divide and differentiate (Fig. 1A) (Fuller and Spradling, 2007; Matunis et al., 2012; Spradling et al., 2011).

**Figure 1:**
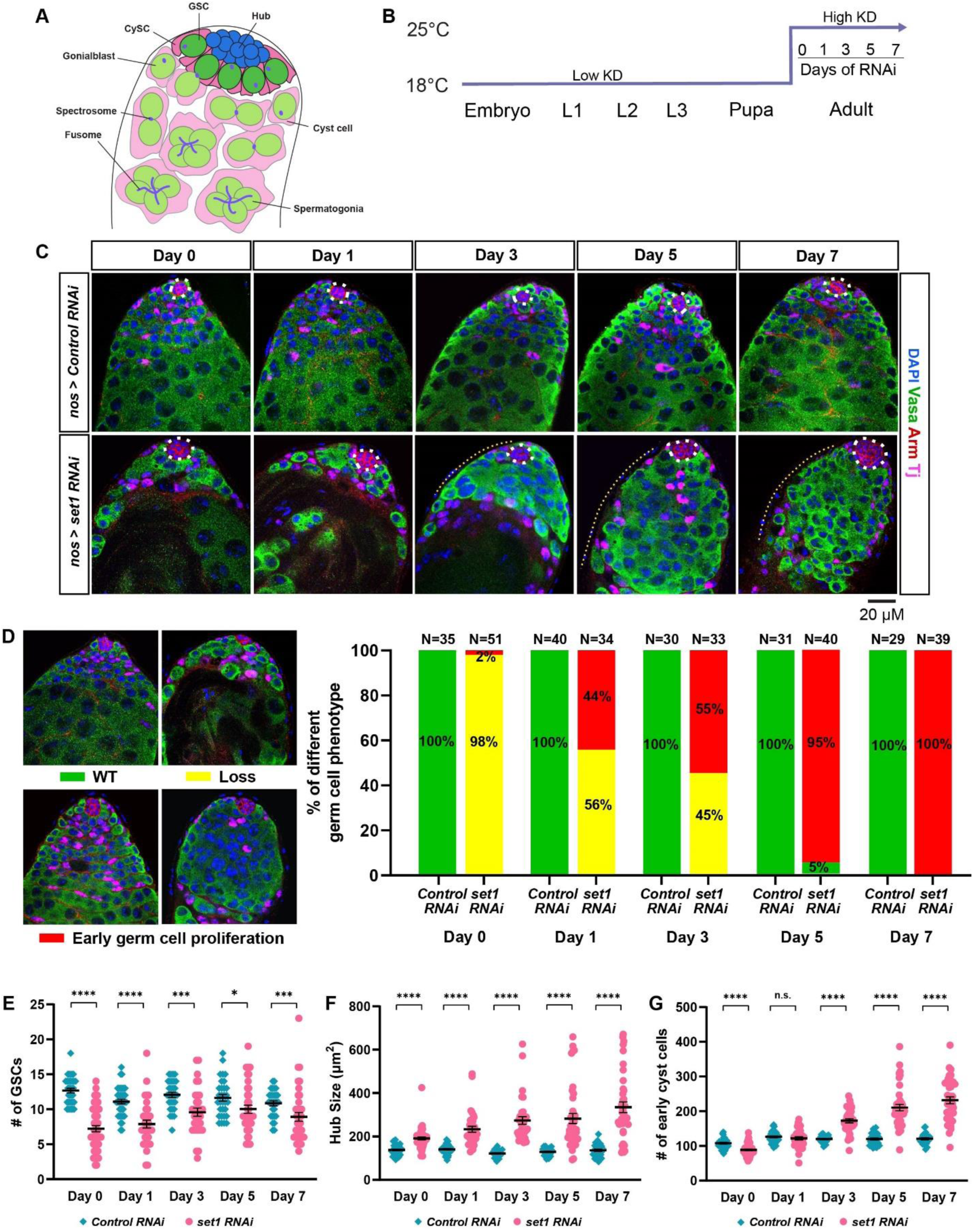
The *Drosophila set1* is required cell autonomously for germline maintenance and differentiation. (**A**) A schematic diagram of the tip of the *Drosophila* testis. GSC: germline stem cell; CySC: cyst stem cell. (**B**) *nos>Control RNAi* (*Ctrl* KD) and *nos>set1 RNAi RNAi* (*set1* KD) flies are grown at 18 ° C until eclosion, and then shifted to 25 ° C for 0, 1, 3, 5, and 7 days, respectively. (**C**) Representative *Ctrl* KD and *set1* KD testes at day 0, 1, 3, 5 and 7 post eclosion stained with the GSC lineage marker Vasa (green), Arm (red) for the hub area (white dotted outline), and Tj (magenta) for the CySC lineage cells. Yellow dotted lines indicate overpopulated early germ cells. (**D**) Quantification of the percent of testes with the representative germline phenotypes: germ cell loss and early-germline overpopulation, present in *Ctrl* KD and *set1* KD testes at 0, 1, 3, 5, and 7 days post eclosion. (**E**) Quantification of GSC number for *Ctrl* KD testes (Day 0: N = 35, Day 1: N = 40, Day 3: N = 30, Day 5: N = 31, Day 7: N = 29) and *set1* KD testes (Day 0: N = 51, Day 1: N = 34, Day 3: N = 33, Day 5: N = 40, Day 7: N = 39). Refer to Table S1. (**F**) Quantification of hub size for *Ctrl* KD and *set1* KD testes. Refer to Table S2. (**G**) Quantification of early cyst cell number for *Ctrl* KD and *set1* KD testes. Refer to Table S3. For (**E-G**): Individual data points and mean values are shown. Error bars represent SEM. \*\*\*\**P*<10^-4^, \*\*\**P*<10^-3^, \**P*< 0.05, n.s.: not significant; unpaired t test to compare two individual datasets to each other.

Two important signaling pathways involved in the maintenance of the GSC niche are the <u>JA</u>nus <u>K</u>inase <u>S</u>ignal <u>T</u>ransducer and <u>A</u>ctivator of <u>T</u>ranscription (JAK-STAT) and <u>B</u>one <u>M</u>orphogenetic <u>P</u>rotein (BMP) signaling pathways (Amoyel et al., 2014; Herrera and Bach, 2019; Inaba et al., 2015; Kiger et al., 2001; Leatherman and Dinardo, 2008, 2010; Shivdasani and Ingham, 2003; Tulina and Matunis, 2001). Furthermore, epigenetic mechanisms, such as chromatin remodeling and histone modifications, have been shown to play important roles in the male germline lineage (Tarayrah and Chen, 2013). In particular, multiple examples have shown the interplay between extrinsic signaling pathways and intrinsic epigenetic mechanisms in the *Drosophila* adult testes (Gleason and Chen, 2023; Vidaurre and Chen, 2021). For example, previous studies have shown that the H3K27me3 histone demethylase, <u>U</u>biquitously transcribed <u>T</u>etratricopeptiderepeat gene on the <u>X</u> chromosome (UTX), acts as a negative regulator of JAK-STAT signaling by maintaining the transcription of *socs36E*, whose proper expression and function maintain the balance between GSCs and CySCs (Amoyel et al., 2016; Issigonis et al., 2009; Singh et al., 2010; Tarayrah et al., 2013). Moreover, genes of the <u>E</u>pidermal <u>G</u>rowth <u>F</u>actor (EGF) signaling pathway may be directly regulated by the H3K27me3 methyltransferase Enhancer of zestes [E(z)] in cyst cells, to ensure proper germ cell identity (Eun et al., 2014). Additionally, the *Drosophila* demethylase for H3K4me3, Little imaginal disc (Lid), regulates JAK-STAT signaling in the male germline to maintain GSC activity and proliferation (Tarayrah et al., 2015). However, unlike the methyltransferase and demethylase for H3K27me3, the methyltransferases for H3K4me3 have not been studied in the *Drosophila* male germline.

SET domain containing 1 (Set1) is the main H3K4me3 methyltransferase in *Drosophila*, whose function is conserved from yeast to mammals (Lee et al., 2007a; Simonet et al., 2007). The post-translational modification of H3K4me3 has previously been shown to correlate with active transcription and is enriched at the promoter and the 5’ coding regions of genes (Bernstein et al., 2002; Heintzman et al., 2007; Pokholok et al., 2005; Santos-Rosa et al., 2002). The role of H3K4me3 in activating transcription is not fully understood, although it is thought that H3K4me3 acts as a docking scaffold for the transcription pre-initiation complex and chromatin remodeling complexes (Vermeulen et al., 2007; Wysocka et al., 2006). In yeast, Set1 catalyzes all methylation forms of H3K4 (i.e., H3K4me1/2/3) and its catalytic activity is modulated *via* a multi-subunit protein complex known as the Complex of proteins associated with Set1 (COMPASS) (Dehe et al., 2005; Miller et al., 2001; Nagy et al., 2002; Roguev et al., 2001). The *Drosophila* Set1 also interacts with the COMPASS complex to catalyze H3K4 di- and tri-methylation (H3K4me2/3) at the promoter proximal regions (Ardehali et al., 2011; Hallson et al., 2012; Mohan et al., 2011). In the *Drosophila* female germline, it has been shown that Set1 regulates GSC maintenance and differentiation, however the mechanism has not been fully determined (Xuan et al., 2013; Yan et al., 2014).

## Results

### The *Drosophila* Set1 regulates germline survival and proper germ cell differentiation in the adult testis

To explore the function of Set1 in the *Drosophila* male germline, a *short hairpin RNA (RNAi)* specifically targeting the coding sequence of the *set1* gene was driven by the early-stage germline driver *nanos-Gal4* (*nos-Gal4*) (Van Doren et al., 1998) for RNAi knockdown (KD) experiments. The *set1* null allele is lethal at the pupal stage and the conventional mosaic analysis cannot be performed due to the genomic location of the *set1* gene very close to the centromere of chromosome 3. Therefore, KD strategy is feasible to interrogate its roles in the adult testis (Ardehali et al., 2011; Hallson et al., 2012). In *nos>set1 RNAi* (*set1* KD) testes, H3K4me3 signal is greatly reduced in the early-stage germ cells, from GSCs to spermatogonial cells (Fig. S1B), compared to the germ cells at the comparable stages in the *nos>mCherry RNAi* testes (*nos>Control RNAi* or *Ctrl* KD, Fig. S1A). In addition, in both *set1* KD and *Ctrl* KD testes, H3K4me3 signals are present in the somatic gonadal cells, indicating germline-specific and efficient inactivation of the methyl-transferase function of Set1 (Fig. S1A-B).

To investigate potential germline defects in the adult testis, a time course experiment was performed where *set1* KD and *Ctrl* KD flies were grown at 18 C° until eclosion and then shifted to 25 C° for 0, 1, 3, 5, and 7 days (Fig. 1B). At Day 0 and Day 1 post-eclosion, 98% and 56%, respectively, of the *set1* KD testes, show germ cell loss phenotype compared to 0% of the *Ctrl* KD testes (Fig. 1C-D). At Day 3 post-eclosion, although 45% of *set1* KD testes have germline loss, 55% have an early germ cell overpopulation phenotype, where the germ cells surrounding the hub form a large, disorganized cluster with very few intercalating cyst cells and are often devoid of or have very few late-stage spermatocytes (Fig. 1D). By Day 5 and 7, 95% and 100% of *set1* KD testes exhibit this early-stage germ cell overpopulation phenotype (Fig. 1D). Throughout this time course, GSC number is consistently reduced in the *set1* KD compared to the *Ctrl* KD testes (Fig. 1E). These results indicate that Set1 is required intrinsically for germ cell maintenance and proper germline differentiation.

In addition to the germline phenotypes observed in the *set1* KD testes, two other phenotypes are detected in the somatic cell lineages of the testis. First, the hub area is significantly increased compared to the *Ctrl* KD testes throughout the time course experiment with higher degree of difference toward the later time points (Fig. 1F). Second, the number of cyst cells is significantly increased in the *set1* KD testes compared to the *Ctrl* KD testes at the later time points in the time course experiments (Fig. 1G). These changes in cyst cell number appear to coincide with the changes in the germline phenotypes over the duration of the time course in the *set1* KD testes, indicating Set1 acts in the germline to regulate somatic gonadal cells in a non-cell-autonomous manner.

Since the Gal4/UAS system may still be functional at 18 C°, knockdown of Set1 at earlier stages of development could contribute to those phenotypes detected in adulthood (Brand and Perrimon, 1993). To address this, we knocked down *set1* in the early germline exclusively in adulthood using the temperature-sensitive Gal80 controlled by the *tubulin* promoter (*tub-Gal80^ts^*). At the permissive temperature (18 C°), functional Gal80 protein inhibits Gal4 from associating with the *UAS* sequences and turning on the *RNAi* expression, but at the restrictive temperature (29 C°) Gal80 is inactivated and Gal4 can associate with *UAS* and activate *RNAi* (McGuire et al., 2003). The *tub-Gal80^ts^, nos>set1 RNAi (set1* ts-KD) and *tub-Gal80^ts^, nos>Control RNAi (Ctrl* ts-KD) flies are grown at 18 C° until eclosion, when they are shifted to 29 C° for 0, 7, 14, 21, and 28 days, respectively, for another time course experiment (Fig. S2A). At Day 0, the *set1* ts-KD and *Ctrl* ts-KD testes have comparable germline morphology, indicating that active Gal80 at 18 C° effectively prevents *set1* knockdown (Day 0, Fig. S2B-C). However, 7 days after shifting to 29 C°, 73% of the *set1* KD testes show germline loss, while 27% exhibit the early germline overpopulation phenotype (Day 7, Fig. S2B-C). These results indicate that the early germline phenotypes can be induced upon knocking down *set1* in the adulthood. In addition to the germline phenotypes detected in the *set1* KD time course (Fig. 1C-E), in *set1* ts-KD testes at later timepoints, a new phenotype classified as “none” is detected, where the testes are almost completely devoid of germ cells (Fig. S2B-C: 48.5% at Day 14, 78% at Day 21 and 65% at Day 28). Additionally, the number of GSCs is significantly reduced at every timepoints in *set1* ts-KD testes compared to the *Ctrl* ts-KD testes (Fig. S2D: Day 7, 14, 21, and 28). Furthermore, the hub area is significantly increased at the later timepoints in the *set1* ts-KD testes (Fig. S2E: Day 14, 21, and 28), whereas the cyst cell number is significantly reduced in *set1* ts-KD testes at the earlier time points (Fig. S2F: Day 7 and Day 14). The variation in somatic gonadal cell phenotypes could be due to the secondary effects induced by the germline *set1* KD and/or the different temperature shift regimes in *set1* KD *versus set1* ts-KD time course experiments, since it has been shown that temperature contributes to germ cell differentiation in *Drosophila* (Gandara and Drummond-Barbosa, 2022, 2023). Together, both strategies to knock down *set1* in the germline shown in Figure 1 and Figure S2 demonstrate that Set1 is required cell autonomously for GSC maintenance and proper germ cell differentiation.

### Set1 is not required in somatic gonadal cells nor late-stage spermatogonial cells for the detected GSC loss phenotype

To determine whether the GSC loss phenotype observed (Fig. 1E and S2D) is specific to knocking down *set1* in the early germline, we first used cell-type-specific strategies to knock down *set1* in the CySCs lineage using the *tj-Gal4* driver (Li et al., 2003). In *tj>set1 RNAi* testes, H3K4me3 signal is reduced in the CySCs and cyst cells compared to the signals in other cell types in the adult testis (Fig. S3A). At 7 days post eclosion, no obvious germline phenotype could be detected in the *tj>set1 RNAi* testes compared to the *tj>Control RNAi* testes (Fig. 2A). The number of GSCs is actually increased in the *tj>set1 RNAi* testes (Fig. 2D), contrary to what has been detected in the *nos>set1 RNAi* testes (Fig. 1E). The hub area has no significant difference between the *tj>set1 RNAi* testes and the *tj>Control RNAi* testes, but the number of cyst cells is significantly increased in the *tj>set1 RNAi* testes (Fig. S3A’ and S3A’’). In addition, to determine the stage specificity of the GSC loss phenotype present in *nos>set1 RNAi* testes, we used the stage-specific *bam-Gal4* driver, which turns on target gene expression specifically in 4-16 stage spermatogonial cells (Chen and McKearin, 2003). In *bam>set1 RNAi* testes, H3K4me3 is diminished in the germline past the 4-cell spermatogonia cell stage but still present in the GSCs and very early-stage germ cells (Fig. S3B). At 7 days post-eclosion, there are no obvious morphological changes in the germline of *bam>set1 RNAi* testes compared to *bam>Control RNAi* testes (Fig. 2B). In addition, the number of GSCs and the hub area in *bam>set1 RNAi* testes and *bam>Control RNAi* testes are not significantly different from each other, even though the *bam>set1 RNAi* testes had significantly more cyst cells (Fig. 2E and Fig. S3B-B’’). In summary, these results indicate that the knockdown of *set1* in the early-stage germ cells is responsible for the detected GSC loss phenotypes shown in Figure 1 and Figure S2.

**Figure 2:**
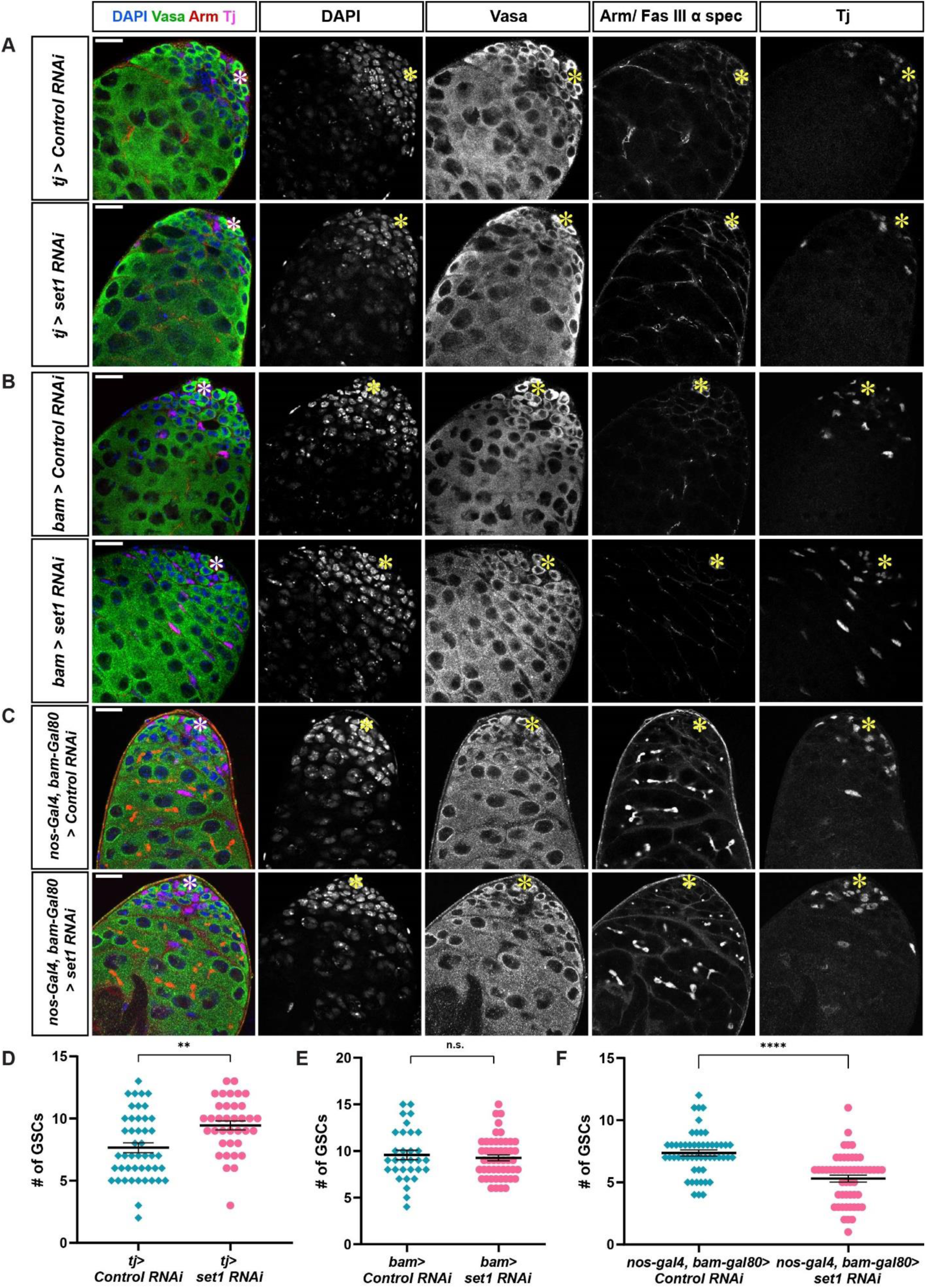
Set1 is dispensable in somatic gonadal and spermatogonial cells to regulate GSC maintenance. (**A**) Representative images of *tj>Control RNAi* and *tj>set1 RNAi* testes at 7 days post eclosion. (**B**) Representative images of *bam>Control RNAi* and *bam>set1 RNAi* testes at 7 days post eclosion. (**C**) Representative images of *nos-Gal4ΔVP16, bam-Gal80>Control RNAi* and *nos-Gal4ΔVP16, bam-Gal80>set1 RNAi* testes at 7 days post eclosion. All images in (**A-C**) stained with germ cell marker Vasa (green), Arm (red) for the hub area (yellow asterisk), and Tj (magenta) for the CySC lineage cells. Scale: 20 μm. (**D**) Quantification of GSC number for *tj>Control RNAi* testes (N = 46) and *tj>set1 RNAi* testes (N = 36). (**E**) Quantification of GSC number for *bam>Control RNAi* testes (N = 31) and *bam>set1 RNAi* testes (N = 50). (**F**) Quantification of GSC number for *nos-Gal4ΔVP16, bam-Gal80>Control RNAi* testes (N = 52) and *nos-Gal4ΔVP16, bam-Gal80>set1 RNAi* testes (N = 52). For (**D-F**): Individual data points and mean values are shown. Refer to Table S1. Error bars represent SEM. \*\*\*\**P*<10^-4^, ** *P*<10^-2^, n.s.: not significant; unpaired t test to compare two individual datasets to each other.

To confirm these findings, we used *nos-Gal4ΔVP16, bam-Gal80* to drive *set1 RNAi* only in the GSCs, gonialblasts, and 2-cell spermatongial cells, since Gal80 protein driven by *bam* promoter prevents *RNAi* expression in the 4-16 stage spermatogonial cells (Eliason et al., 2018). In the *nos-Gal4ΔVP16, bam-Gal80 >set1 RNAi* testes, 95% had a reduction in H3K4me3 specifically in GSCs, gonialblasts and early-stage spermatogonial cells (Fig. S3C). At 7 days post eclosion, the hub size and number of cyst cells in the *nos-Gal4ΔVP16, bam-Gal80>set1 RNAi* testes are comparable to those in *nos-Gal4ΔVP16, bam-Gal80>Control RNAi* testes (Fig. S3C’ and S3C’’). However, the number of GSCs in the *nos-Gal4ΔVP16, bam-Gal80>set1 RNAi* testes is significantly reduced compared to the *nos-Gal4ΔVP16, bam-Gal80>Control RNAi* testes (Fig. 2F). Together, these data demonstrate that the knockdown of *set1* in the early-stage germ cells can partially account for the GSC loss phenotype seen in the *nos>set1 RNAi* testes. The differences in other phenotypes, especially the non-cell-autonomous phenotypes, are likely due to the cell-type- and stage-specificities, as well as the strength of the KD effects, using different drivers. We next focused on using *nos>set1 RNAi* testes for the following experiments to understand the mechanisms underlying the *set1* KD germline phenotypes.

### The methyltransferase activity of Set1 is required for its proper activity in germ cells

To ensure the phenotypic effects seen in the *nos-Gal4>set1 RNAi* testes (Fig. 3A, adapted from Fig. 1D) are due to the knockdown of the *set1* gene, but not the off-targets using RNAi, a rescue experiment was performed. In *nos-Gal4>set1 RNAi* testes, a *GFP*-tagged *set1* cDNA transgene is expressed using the same *nos-Gal4* driver. Furthermore, since the *set1 RNAi* target sequence is within the coding region of *set1*, silent mutations were made within the target region to prevent KD of the transgene, namely a GFP-tagged *set*1 cDNA transgene with the RNAi silent mutations (*WT Rescue*, Fig. 3B and S4A). As controls, three additional transgenes were generated and expressed at the same genetic background: the WT Rescue transgene with the RNAi silent mutations and an E-to-K mutation in the catalytic SET domain (*Mut Rescue*, Fig. 3C and S4B), a GFP-tagged *set1* cDNA transgene without the RNAi silent mutations (*WT*, Fig. 3D and S4C), and a GFP transgene with a nuclear localization signal (*GFP-NLS*, Fig. 3E and S4D). At Day 1 and Day 5 post eclosion, 93% and 77% of the *nos-Gal4>WT Rescue, set1 RNAi* testes show full rescue of the *set1* knockdown phenotypes (Fig. 3B’-B” and S4A’). By contrast, in the *nos-Gal4>Mut Rescue, set1 RNAi* testes, only 2% show full rescue, while 82% and 38% exhibit the germline loss phenotype, as well as 16% and 60% display the early germ cell proliferation phenotype at Day 1 and Day 5 post eclosion, respectively (Fig. 3C’-C” and S4B’), indicating that the catalytic SET domain is indispensable for the normal function of Set1 in the early-stage male germline. Both the *nos-Gal4>WT, set1 RNAi* testes and *nos-Gal4>GFP-NLS, set1 RNAi* testes show similar results compared to the *nos-Gal4>Mut Rescue, set1 RNAi* testes (Fig. 3D’-D” and S4C’; Fig. 3E’-E” and S4D’), suggesting that adding back *set1* without changing the RNAi recognition sequences or a non-specific GFP-NLS transgene does not rescue the *set1* knockdown phenotypes in the male germline.

**Figure 3:**
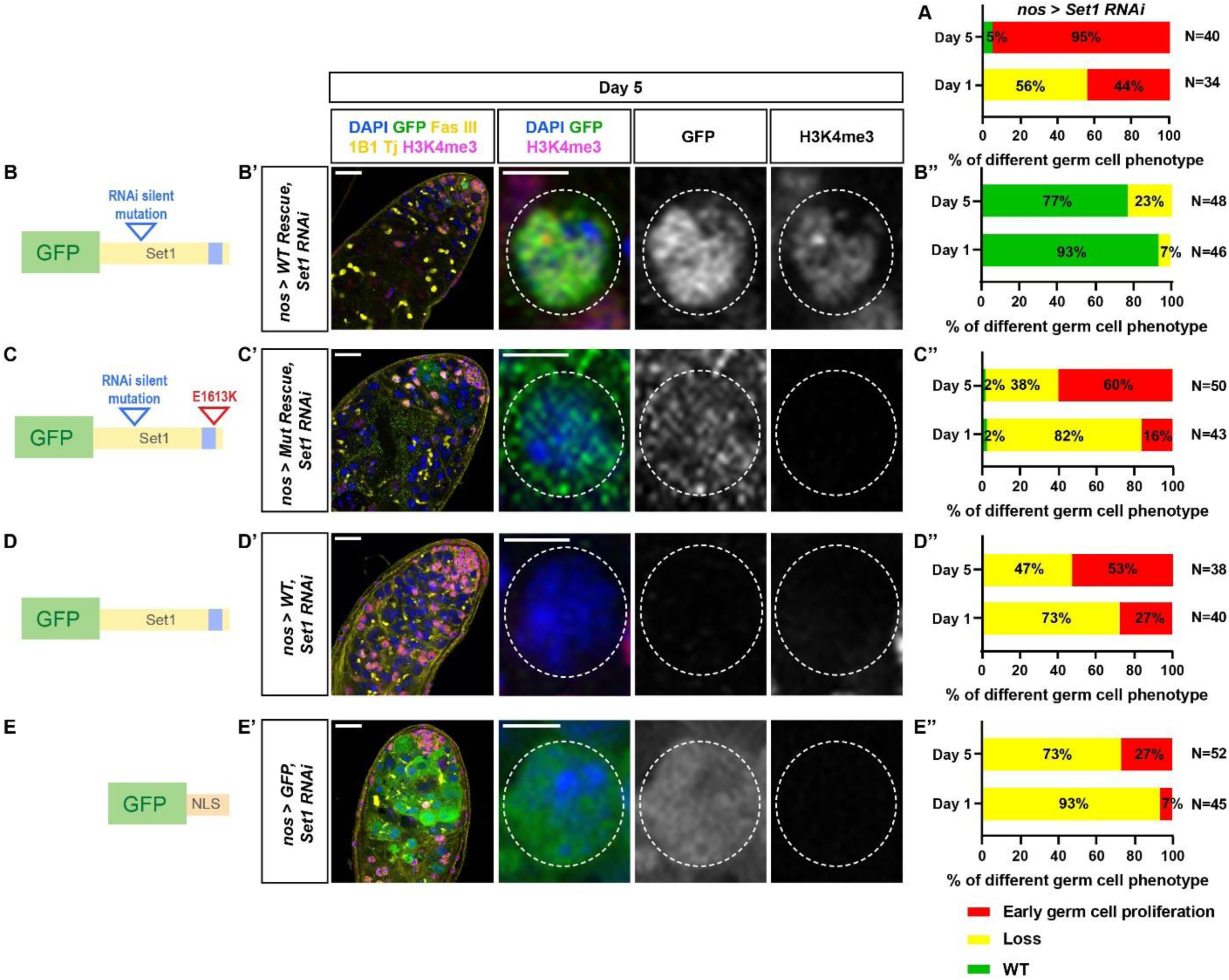
The methyltransferase activity of Set1 is required for its function in the male germline. (**A**) Quantification of the percent of testes with the representative germline phenotypes: germ cell loss and early-germline overpopulation, present in *nos-Gal4>set1 RNAi* testes at two time points at 1 and 5 days post eclosion (adapted from Fig. 1D). (**B-E”**) Cartoon depictions: the *set1* cDNA transgene with the RNAi recognition sequences mutated, named WT rescue (**B**); the *set1* cDNA transgene with the RNAi recognition sequences mutated and an E→K amino acid change in the SET domain, named Mut rescue (**C**); the *set1* cDNA transgene without the RNAi recognition sequences mutated, named WT (**D**); the *GFP* cDNA with the *nuclear localization* sequence, named GFP (**E**). Representative images of *nos-Gal4>WT Rescue, set1 RNAi* testis (**B’**), *nos-Gal4>Mut Rescue, set1 RNAi* testis (**C’**), *nos-Gal4>WT, set1 RNAi* testis (**D’**), *nos-Gal4>GFP, set1 RNAi* testis (**E’**), all testes are from males 5 days post eclosion, immunostained with H3K4me3 (magenta), GFP (green), Fas III (yellow) for the hub region, and Tj (yellow) for the CySC lineage cells. White dotted outline: germ cell expressing the corresponding transgenes. Scale: 20 μm for the testis image; 5 μm for individual germ cell images. Quantification of the percent of testes with the representative germline phenotypes: germ cell loss and early-germline overpopulation, present in *nos-Gal4>WT Rescue, set1 RNAi* testes (**B’’**), *nos-Gal4>Mut Rescue, set1 RNAi* testes (**C’’**), *nos-Gal4>WT, set1 RNAi* testes (**D’’**), *nos-Gal4>GFP, set1 RNAi* testes (**E’’**), at two time points at 1 and 5 days post eclosion.

Furthermore, the GFP signals from both the *nos-Gal4>WT Rescue* and the *nos-Gal4> Mut Rescue* GFP-tagged fusion proteins are exclusively detected in the early germline nuclei, indicating that the silent mutations make these transgenes resistant to the *RNAi* knockdown, allowing for their proper expression and localization (Fig. 3B’-C’ and Fig. S4A’-B’). In contrast, no GFP signal could be detected in the *nos-Gal4>WT, set1 RNAi* testes without the RNAi silent mutations, indicating effective knockdown. In summary, these results support that the germline phenotypes observed in *nos-Gal4>set1 RNAi* testes is due to the methyltransferase activity of the Set1 protein.

### *Set1* regulates gene expression of multiple signaling pathway components in the *Drosophila* testis

To understand the molecular mechanisms underlying the function of Set1 in the germline, RNA-seq was performed to compare the transcriptomes between *nos>Control RNAi* (*Ctrl* KD) and *nos>set1 RNAi* (*set1* KD) testes at 0, 1, 3, and 5 days post eclosion, respectively (Fig. 4 and S5). To obtain a global picture of the transcriptome changes between *Ctrl* KD and *set1* KD testes at different time points, principal component analyses were performed for all 24 samples (Fig. S5A). Among the 24 samples, the *Ctrl* KD samples cluster together and the *set1* KD samples cluster together (Fig. S5A). In addition, the biological replicates for each *set1* KD sample timepoints cluster more closely with one another than the replicates of the *Ctrl* KD sample timepoints (Fig. S5A). This can be explained since all 12 *Ctrl* KD samples should have similar gene expression patterns to one another given the age difference between each consecutive timepoints is only 1-2 days. In contrast, the 12 *set1* KD samples do not, as visible changes could be detected at each timepoints in the *set1* KD testes. To better analyze the transcriptome changes between the *Ctrl* KD and *set1* KD samples at every timepoint, a heatmap of all the differentially expressed genes was created (Fig S5B). The 16,610 genes included in the heatmap, according to the *Drosophila* Release 6 reference genome (dos Santos et al., 2015), are classified into four groups. Group 1 contains genes that are highly expressed at the earlier timepoints of the *set1* KD samples (Day 0 and 1), while Group 2 has the genes that are highly expressed at the later timepoints of the s*et1* KD samples (Day 3 and 5). Additionally, Group 3 consists of the genes that are downregulated in the *set1* KD samples. Finally, Group 4 is comprised of the genes with variable expression profiles across all the timepoints and thus did not fit the clear expression patterns seen for the genes in groups 1-3 (Fig. S5B).

**Figure 4:**
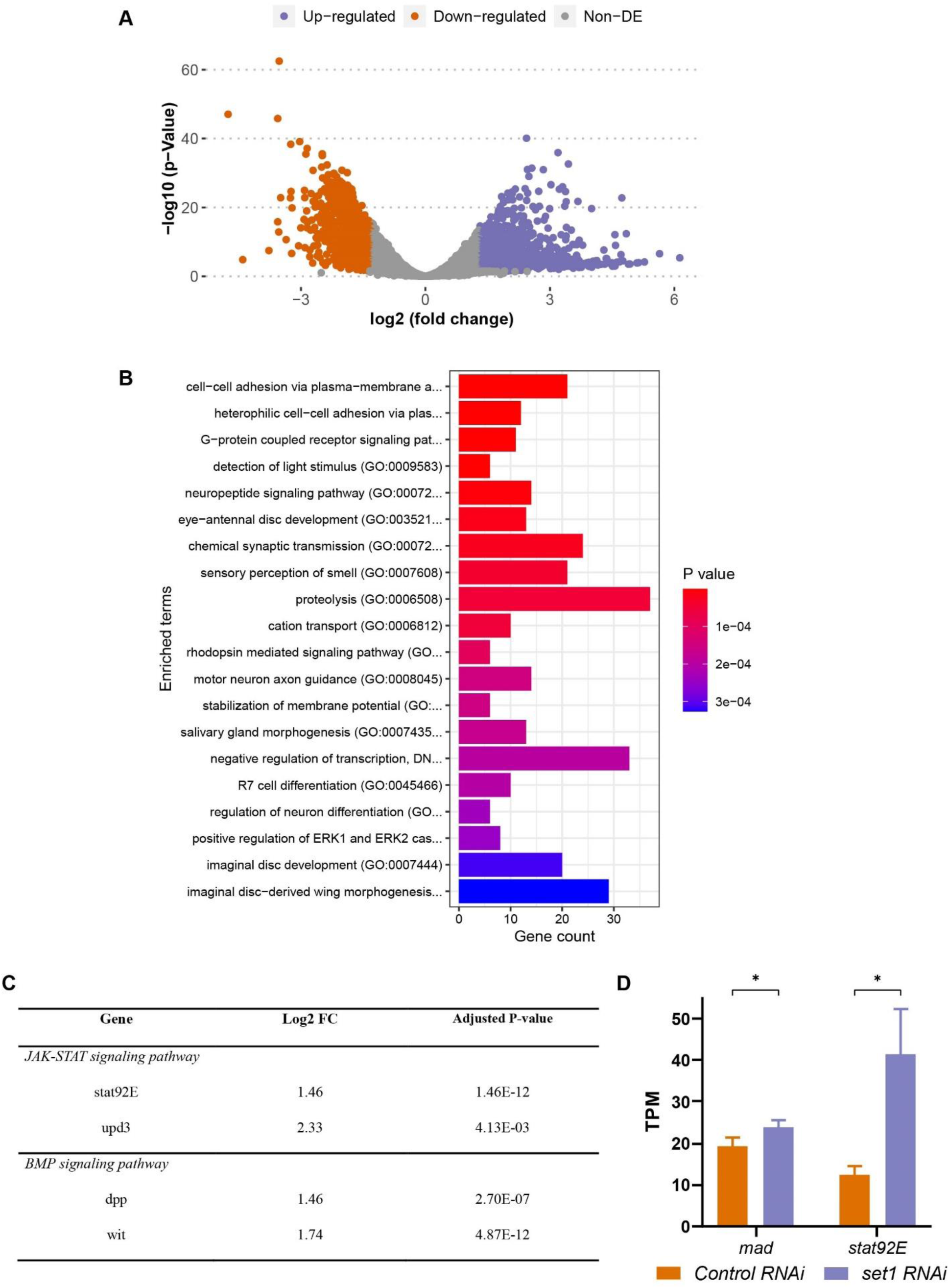
JAK-STAT and BMP signaling pathway components have increased expression in *set1* knockdown testis. (**A**) A volcano plot of differentially expressed genes (DEGs) in *set1* KD testes *versus Ctrl* KD testes at 3 days post eclosion (≥log_2_1.3= 2.46-fold change, *P* < 0.05). (**B**) GO analysis of the 1,216 upregulated genes identified from (**A**). GO analysis of the 764 downregulated genes does not show any significant category. (**C**) A table of several JAK-STAT and BMP signaling pathway components which are upregulated in *set1* KD testes compared with *Ctrl* KD testes 3 days post eclosion. (**D**) Expression levels of *mad,* and *stat92E* in *set1* KD testes *versus Ctrl* KD testes. Error bars represent SEM, N = 3 biological replicates, \**P* < 0.05 using Unpaired t test.

After these initial analyses on all 24 samples, we focused on Day 3 of the *set1* KD samples for two reasons: First, it is the timepoint when the germline mass phenotype becomes predominant; second, it has the most differentially expressed genes even with a stringent cutoff (≥log_2_1.3= 2.46-fold and *P*<0.05, Fig. 4A and S5C). At this timepoint, 1,216 genes are significantly upregulated, and 764 genes are significantly downregulated in the *set1* KD samples compared to the *Ctrl* KD samples (Fig. 4A and S5C). GO analysis of the 1,216 upregulated genes show enriched functions, with a significant category to be “negative regulation of transcription” (Fig. 4B).

Interestingly, components of both JAK-STAT and BMP signaling pathways are found to be upregulated in the *set1* KD compared to the *Ctrl* KD samples at Day 3 (Fig S5D). For example, the JAK-STAT pathway genes, *stat92E* and *upd 3*, and the BMP pathway genes, *dpp* and *wit*, are found to be significantly upregulated (Fig. 4C). Given the importance of both signaling pathways in the testis (Inaba et al., 2015; Kawase et al., 2004; Kiger et al., 2001; Leatherman and Dinardo, 2008, 2010; Schulz et al., 2004; Singh et al., 2010; Tulina and Matunis, 2001), these results indicate that the early-stage germ cell overpopulation phenotype could be due to ectopic expression of both JAK-STAT and BMP signaling pathway genes.

### The interactions between *set1* and JAK-STAT or BMP pathway genes are responsible for the early-stage germline phenotypes

To further test the functional relevance of both JAK-STAT and BMP signaling pathways with the *set1* KD phenotypes, we examined two key downstream genes of JAK-STAT and BMP pathways, *mad* and *stat92E*, respectively. Both genes are identified to be upregulated in the *set1* KD testes compared to the *Ctrl* KD testes using the RNA-seq dataset (Fig 4D). However, given that Set1 generates H3K4me3 to activate transcription, the direct target genes of Set1 are likely to be downregulated in *set1* KD testes. Nevertheless, we reasoned that the ectopic expression of *mad* or *stat92E* could contribute to the early-germline over-population phenotypes in the *set1* KD testes. Next, we tested their genetic interactions with *set1*.

In heterozygous loss-of-function *mad^12^* testes (Raftery et al., 1995), pMad is only very faintly detectable in the GSCs (Fig. 5A: *mad^12^/+*) and 100% show normal morphology (Fig. 5C). In contrast, in the *set1* KD testes, pMad can be detected in GSCs and even ectopically in the spermatogoial cells (Fig. 5A: *nos>set1 RNAi*). Here, 85% of the *set1* KD testes exhibit the early germ cell overpopulation phenotype (Fig. 5C). Even though both pMad patterns are detectable in *set1* KD testes with one copy of the *mad^12^* allele (Fig. 5A: *mad^12^/+; nos>set1 RNAi*), the addition of one *mad^12^* allele to the *set1* KD background reduce the occurrence of the early germ cell overpopulation phenotype from 85% to 65% (*P*< 0.05, Fig. 5C).

**Figure 5:**
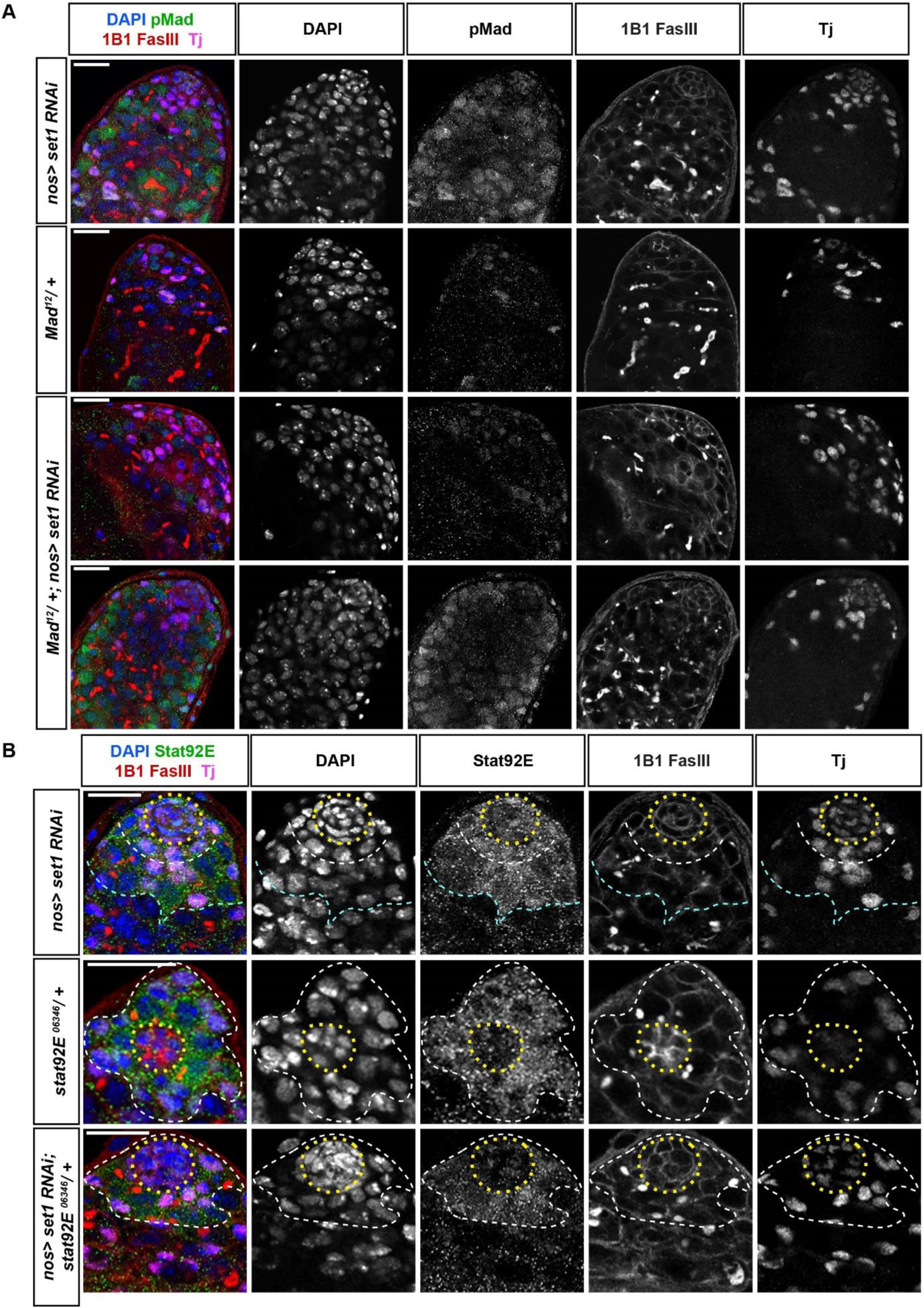
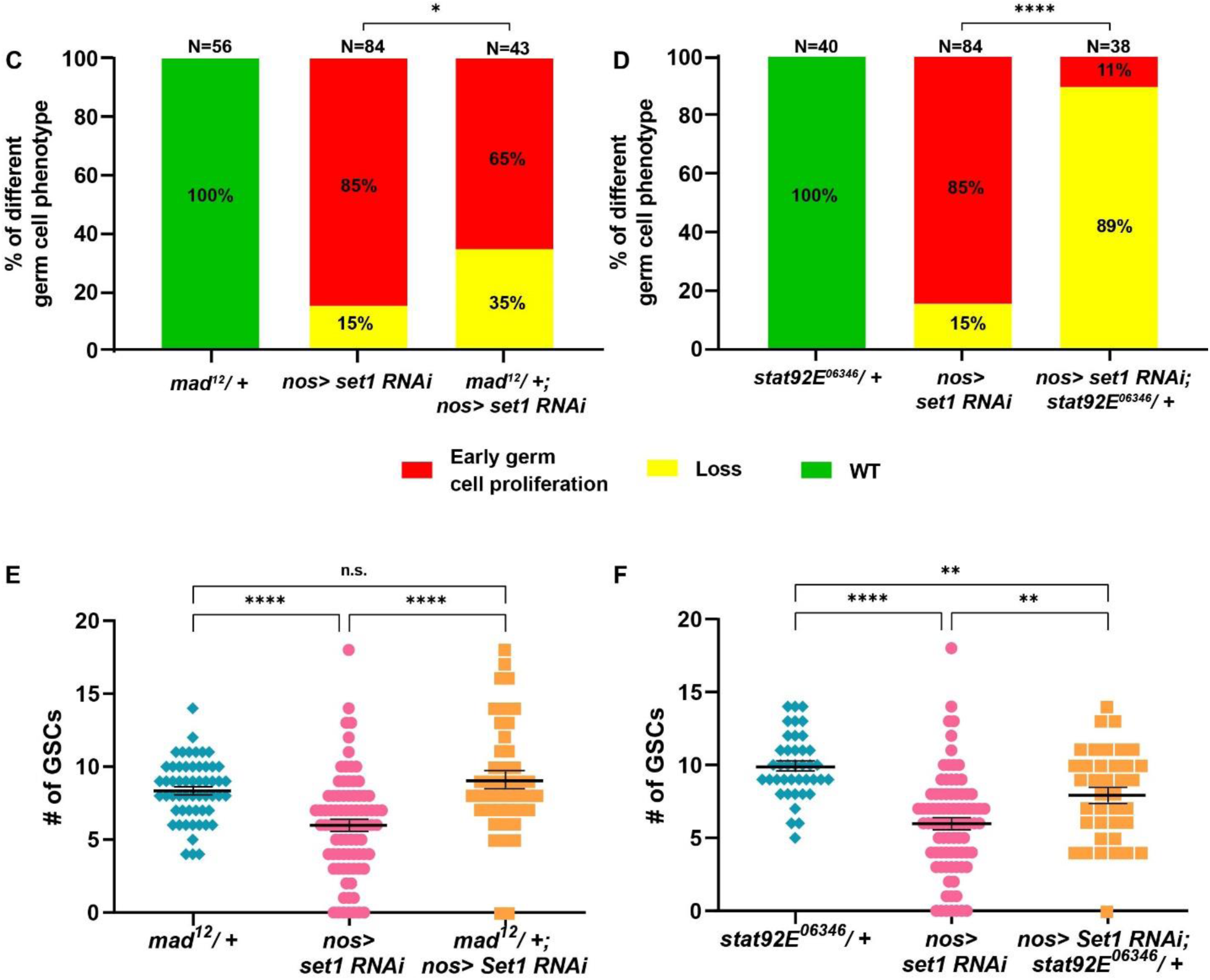
Set1 regulates key JAK-STAT and BMP signaling pathway components. (**A**) Representative images of *nos-Gal4>set1 RNAi*, *mad^12^/+*, and *mad^12^/+; nos-Gal4>set1 RNAi* testes at 5 days post eclosion, immunostained with pMad (green), Fas III (red) for the hub region, and Tj (magenta) for the CySC lineage cells. (**B**) Representative images of *nos-Gal4>set1 RNAi*, *stat92E^06346^/+*, and *stat92E^06346^/+; nos-Gal4>set1 RNAi* testes at 5 days post eclosion, immunostained with Stat92E (green), Fas III (red) for the hub region, and Tj (magenta) for the CySC lineage cells. Scale: 20 μm. In (**B**): yellow dotted outline: hub; white dotted outline: the GSCs and CySCs region; cyan dotted outline: spermatogonial region. (**C**) Quantification of the percent of testes with the representative germline phenotypes: germ cell loss and early-germline overpopulation, present in *nos-Gal4>set1 RNAi*, *mad^12^/+*, and *mad^12^/+; nos-Gal4>set1 RNAi* testes at 5 days post eclosion. *P*< 0.05, Chi-square test. (**D**) Quantification of the percent of testes with the representative germline phenotypes: germ cell loss and early-germline overpopulation, present in *nos-Gal4>set1 RNAi*, *stat92E^06346^/+*, and *stat92E^06346^/+; nos-Gal4>set1 RNAi* testes at 5 days post eclosion. \*\*\*\**P* < 10^-4^, Chi-square test. (**E**) Quantification of GSC number for *mad^12^/+* (N = 56), *nos-Gal4>set1 RNAi* (N = 84), and *mad^12^/+; nos-Gal4>set1 RNAi* (N = 43) testes at 5 days post eclosion. (**F**) Quantification of GSC number for *stat92E^06346^/+* (N = 40), *nos-Gal4>set1 RNAi* (N = 84), and *stat92E^06346^/+; nos-Gal4>set1 RNAi* (N = 38) testes at 5 days post eclosion Individual data points and mean values are shown. Refer to Table S1. Error bars represent SEM. \*\*\*\**P*<10^-4^, ** *P*<10^-2^, n.s.: not significant; unpaired t test to compare two individual datasets to each other.

Additionally, in heterozygous testes with the loss-of-function *stat92E^06346^*allele (Hou et al., 1996), Stat92E expression is detected only in the GSCs (white outline in Fig. 5B: *stat92E^06346^/+*), but in *set1* KD testes Stat92E is not only found in the GSCs but also in hub cells (yellow outline in Fig. 5B: *nos>set1 RNAi*) and in spermatogonial cells (cyan outline in Fig. 5B: *nos>set1 RNAi*). Interestingly, upon compromising Stat92E with *stat92E^06346^* in the *set1* KD background, the expression of Stat92E is no longer found in hub cells (yellow outline in Fig. 5B: *nos>set1 RNAi; stat92E^06346^/+*) or in the spermatogonial cells. Furthermore, when the level of Stat is reduced in *nos>set1 RNAi; stat92E^06346^/+* testes, the incidence of the early-stage germ cell overpopulation phenotype decreases from 85% to 11% (*P* < 10^-4^, Fig. 5D). These results suggest that enhanced Stat92E expression upon inactivating Set1 significantly contributes to the early germ cell overpopulation phenotype *set1* KD testes. Moreover, compromising BMP pathway (by *mad^12^/+*) fully suppress the GSC loss phenotypes in *set1* KD testes (Fig. 5E), while compromising JAK-STAT pathway (by *stat92E^06346^/+*) could partially suppress the GSC loss phenotype in *set1* KD testes (Fig. 5F). However, neither conditions could fully restore the loss of some spermatogonial cells in the *set1* KD testes (Fig. 5C-D), despite to a less degree, suggesting that other downstream factors are contributing to the loss of early-stage germ cells at the *set1* KD background. Together, these genetic interaction results support the functional relationships between *set1* and BMP or JAK-STAT pathways, with the primary role by the BMP signaling to the GSC loss phenotype and the predominant contribution by the JAK-STAT pathway to the early-stage germline overpopulation phenotype, consistent with the transcriptome results shown in Figure 4.

## Discussion

This study explores the *in vivo* roles of *Drosophila* Set1, a histone methyl-transferase responsible for ‘writing’ the H3K4me3 histone modification, in the *Drosophila* adult testis. Through a series of cell-type and stage-specific RNAi knockdown *via* a time course regime, our findings unveil a fascinating progression of germline defects (Fig 6A-D). These *set1* loss-of-function phenotypes have both cellular and molecular specificities: First, the phenotypes are predominantly linked to the loss of Set1 function in early-stage germ cells, as compromising Set1 in late-stage germ cells or somatic gonadal cells fail to replicate these defects. Moreover, we conducted experiments utilizing transgenes encoding both the wild-type and catalytic inactive forms of Set1. Only the wild-type Set1 effectively rescues the knockdown phenotypes, underscoring the specificity of these effects to the methyl-transferase activity of Set1. To further understand the underlying molecular mechanisms, we performed RNA-seq experiments to identify genes with changed expression upon knocking down s*et1 g*ene. Through this assay, we identified several key signaling pathway genes, such as *stat92E* and *mad*, the downstream components of the JAK-STAT and BMP signaling pathways, respectively, have increased expression when *set1* is knocked down. Genetic interaction analyses further confirm the functional relationship between *set1* and these signaling pathways, as mutations of the *stat92E* and *mad* genes suppress the *set1* knockdown phenotypes. Collectively, our investigations shed light on the molecular mechanisms at play, enhancing our comprehension of the pivotal decision-making process governing proliferation *versus* differentiation within adult stem cell lineages. This insight bears significant implications for both stem cell biology and the broader field of cancer biology.

**Figure 6:**
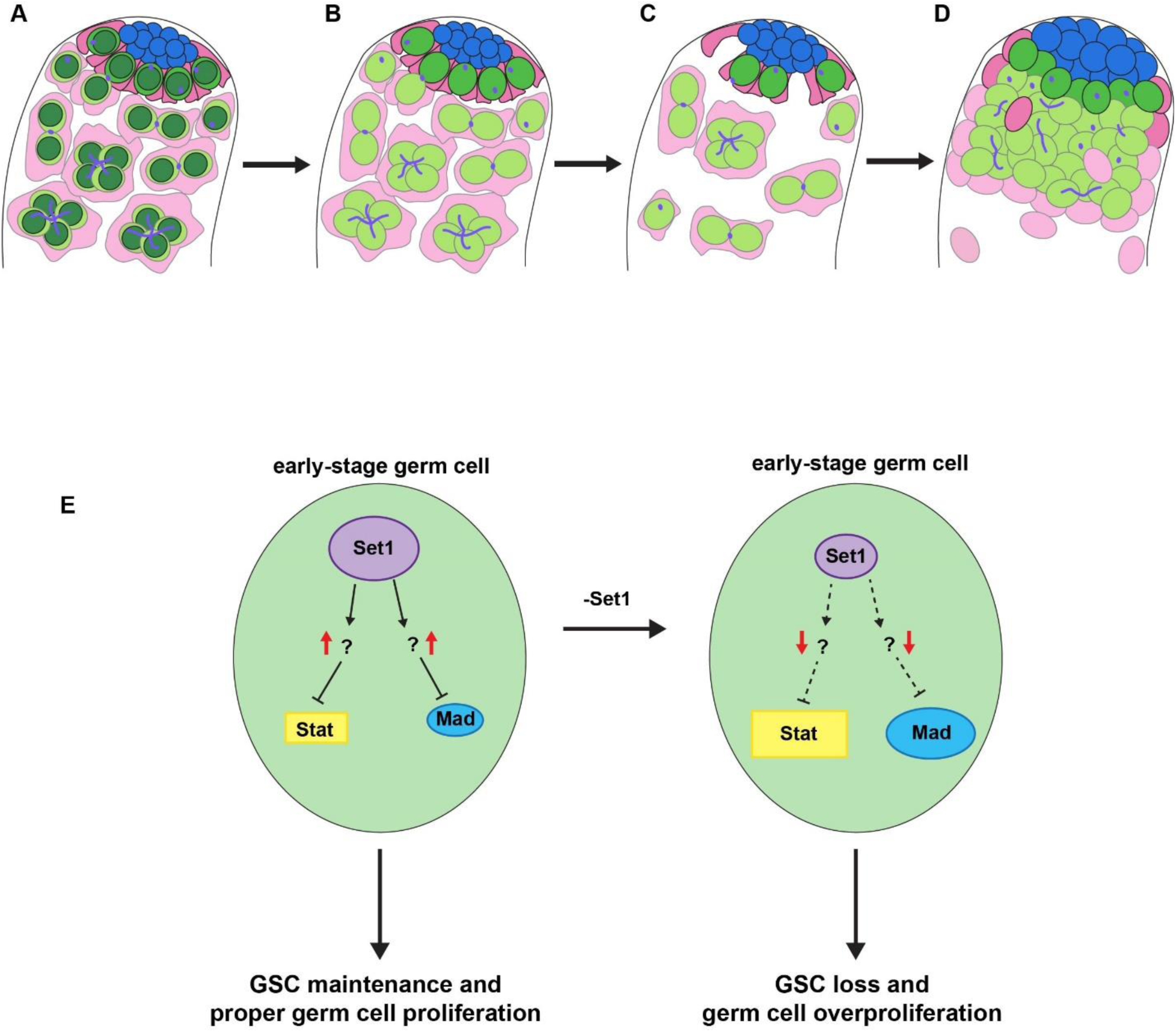
Illustration of the progression of *set1* knockdown phenotypes in adult testis. (**A**) In a wild-type testis, H3K4me3 (dark green) is present in all cell types of the testis, but for simplicity the presence of H3K4me3 (dark green) is only shown in the germ cells, including GSCs (bright green), gonialblasts and spermatogonia (light green). CySCs (magenta), cyst cells (pink), hub cells (blue). Knockdown of *set1* in the early germline results in the reduction of H3K4me3 in GSCs, gonialblasts and spermatogonia (**B**). The phenotypes at the earlier time points include a reduction of germ cells and cyst cells (**C**). The phenotypes at the later time points include overpopulated early-stage germ cells ectopically expressing Stat92E and Mad, as well as increased hub region (**D**). (**E**) Putative pathways of Set1 function in early-stage germline including GSCs. Given both *stat92E* and *mad* have increased expression in the *set1* KD testes, they are likely indirect targets of Set1. One possible explanation is that Set1 directly regulates inhibitors of both *stat92E* and *mad*. Compromising *set1* leads to the downregulation of these inhibitors and ectopic expression of *stat92E* and *mad*, resulting in the GSC loss and early-stage germ cell overproliferation phenotypes.

### Set1 may regulate JAK-STAT and BMP signaling pathway inhibitors in the *Drosophila* male germline

Here, our RNA-seq results reveal that upon knockdown of *set1* in the *Drosophila* male germline, a key downstream factor of the JAK-STAT signaling pathway, *stat92E*, is upregulated (Fig. 4D). This is unexpected for the direct targets of Set1, given that Set1 is the methyltransferase for the transcriptional activating mark, H3K4me3. In addition, previous ChIP-seq results using progenitor germ cell-enriched *bam* testes (McKearin and Spradling, 1990) indicate that H3K4me3 is not enriched at the promoter region of the *stat92E* gene [Fig S6A, (Gan et al., 2010)]. These results suggest that *stat92E* is most likely an indirect target of Set1. However, given the suppression of the early germ cell overpopulation phenotype upon the addition of a single copy of a *stat92E* mutation at the *set1* KD background, they should both function within the same regulatory pathway in some capacity. We propose that Set1 acts in the *Drosophila* male germline by regulating the transcriptional activity of a JAK-STAT inhibitor. When Set1 is compromised, the expression of this inhibitor is downregulated, allowing for the ectopic expression of Stat92E (Fig. 6E). One candidate JAK-STAT inhibitor is protein tyrosine phosphatase 61F (Ptp61F). Ptp61F is a known inhibitor of JAK-STAT that has been previously identified as a target of *little imaginal discs* (*lid*), the H3K4 demethylase, in the germline (Baeg et al., 2005; Tarayrah et al., 2015). In addition, *ptp61F* is downregulated in our RNA-seq data (Fig. S6G) and is enriched for H3K4me3 at its promoter regions (Fig. S6C).

In addition to Stat92E, the downstream factor of the BMP signaling pathway, *mad*, is also upregulated in the *set1* KD background. Unlike *stat92E,* H3K4me3 is enriched at the promoter region of the *mad* gene in progenitor germ cells (Fig. S6B) (Gan et al., 2010). However, given that *mad* is slightly upregulated (Fig. 4D) and suppresses the early germ cell overpopulation phenotype when a single copy of *mad* mutation is combined with the *set1* KD condition (Fig. 5A and 5C), we hypothesize BMP inhibitors could be direct targets of Set1, whose downregulation upon inactivation of Set1 could lead to the ultimate upregulation of *mad*. When examining the candidate BMP inhibitors Cul-2, MAN1, and Ube3a: Cul-2 is a BMP inhibitor in the *Drosophila* female germline while both MAN1 and Ub3a are BMP inhibitors in other *Drosophila* tissues, as shown previously (Ayyub et al., 2015; Duan et al., 2020; Laugks et al., 2017; Li et al., 2016; Wagner et al., 2010). All three BMP inhibitors have downregulated expression in the *set1* KD testes (Fig. S6G) and H3K4me3 enrichment at their corresponding promoter regions (Fig. S6D-F) (Gan et al., 2010). Additional experiments are needed to determine whether these inhibitors of JAK-STAT and BMP signaling are *bona fide* direct target genes and have a functional relationship with Set1 in the *Drosophila* male germline.

### Interactions of Set1 with other histone modifying enzymes in regulating key signaling pathways

In *Drosophila*, the *little imaginal discs* (*lid*) gene encodes the histone demethylase responsible for the ‘erasing’ of H3K4me3 histone modification (Eissenberg et al., 2007; Lee et al., 2007b), whose enzymatic activity directly antagonizes the function of Set1. It has been shown that in the male germline, Lid is required for the proper levels of Stat92E. In the *lid* knockdown or mutant cells, the *stat92E* expression is decreased (Tarayrah et al., 2015). Therefore, the two histone modifying enzymes with the opposite activities, Set1 and Lid, display the opposite regulation of *stat92E* expression. This study introduces a new epigenetic regulator within the intricate framework of the JAK-STAT signaling pathway in the *Drosophila* testis niche. Together, these studies underscore the intricate balance controlled by both a histone methyl-transferase and a histone demethylase in orchestrating a major signaling pathway.

### Commonality and difference of the roles of *Drosophila* Set1 in other stem cell systems

As the major H3K4 methyl-transferase in *Drosophila*, Set1 has also been shown to have functional roles in the female germline. For example, it has been demonstrated that Set1 acts with an E3 ubiquitin ligase Bre1 to generate the majority of H3K4me3 modification and regulate female GSC maintenance and differentiation. Knockdown of *set1* in the early-stage germline results in significant loss of GSCs in the germaria due to decreased expression of BMP signaling components, such as Mad or Dad (Bray et al., 2005; Xuan et al., 2013). This phenotype is similar to the male GSC loss phenotype detected at the early course of the germline defects in this study. However, in the male germline, inactivation of Set1 seems to affect the JAK-STAT signaling pathway more than the BMP pathway; and this effect progresses into an upregulation of Stat92E, leading to overpopulated early-stage germ cells at the late course of the germline defects in this study. Based on an extensive RNAi screen that identified hundreds of genes with functional roles in the female germline, *set1* stands out as a crucial factor required for female GSC maintenance and proper differentiation. When *set1* is strongly knocked down in the female germline, ovaries exhibited a spectrum of phenotypes, including empty ovarioles likely caused by female GSC loss and so-called “pseudoegg” chambers enriched with undifferentiated cells (Yan et al., 2014). The latter phenotype resembles the early-germline overpopulation phenotype exhibited in the *set1* male germline knockdown phenotype shown in this work (Fig. 1C-D). Moreover, the non-cell-autonomous roles of Set1 in the *Drosophila* ovary are more substantial from somatic gonadal cells to impact female germline. For example, compromising Set1 in different somatic gonadal cells in the ovary results in either GSC loss or overpopulation of GSC-like cells in the germaria (Xuan et al., 2013). In contrast, reducing Set1 in the somatic gonadal cells in the testis does not seem to cause any obvious phenotype, as shown in this study (Fig. 2A). However, in the *Drosophila* testis the non-cell-autonomous effects are identified from germ cells to influence the niche architecture and cyst cells (Fig. 1F-G).

In the somatic stem cell lineages, two types of neuroblast (NBs), type I NBs (NB-I) and type II NBs (NB-II), differ in their number (e.g., 184 NB-Is and 16 NB-IIs) and lineage progression to ultimately produce the diverse neuron and glia cells composing the brain (Bello et al., 2008; Boone and Doe, 2008). Interestingly, Set1 seems to be dispensable for the NB-II differentiation, which is distinct from the profound functions of Set1 in both ovary and testis systems for GSC differentiation (Yan et al., 2014), suggesting different requirements of histone modifications and their modifying enzymes in different stem cell systems.

### The progressed germline phenotypes due to loss-of-function of Set1 necessitates cautions using histone methyl-transferase inhibitors in cancer therapy

Many short-lived cell types in the body are produced from adult stem cell precursors, either continuously or in response to physiological signals or trauma. During these processes, the occurrence of genetic lesions or epigenetic mis-regulation may induce a shift in their cellular homeostasis to become ever-dividing cells at the expense of proper cellular differentiation, leading to the initiation of diseases such as cancers.

The COMPASS complex has been shown to be involved in pathogenesis such as cancer (Shilatifard, 2012). Here, our results show profound phenotypes due to male germline knockdown of *set1*, which start with a loss of germ cells, eventually culminating in an intriguing early-stage germline tumor phenotype within a relatively short period of time (Fig. 1B-D). This delayed appearance raises an intriguing possibility: a resilient subset of ‘survivor cells,’ which evade early cell death, initiates a phase of uncontrolled and rapid over-proliferation of undifferentiated cells, reminiscent of the “cancer stem cell” phenomenon (Beck and Blanpain, 2013). Since different histone methyl-transferases have been targeted for therapeutic treatment against cancers, our discovery of an initial cell loss followed by over-proliferation in response to the knockdown of a key histone methyl-transferase raises concerns about the potential use of these enzyme inhibitors in cancer therapy (Morera et al., 2016).

## Data availability

GEO accession number for the RNA-seq data in progress.

## Author contributions

V.V. and X.C. conceptualized the study. V.V. and A.S. performed all experiments. V.V. and T.L., and W.L.K performed data analyses. V.V. and X.C. wrote the manuscript.

## Acknowledgments

We thank Drs. S. Myong, D. Drummond-Barbosa, C. Wu, and Y. Kim for critical comments. We thank Chen lab members for insightful suggestions. We thank Yingying Li for generating the transgenes. We thank Johns Hopkins Integrated Imaging Center for confocal imaging. Supported by NIH 5T32GM007231 (V.V.), F31GM134641 (V.V.), Division of Intramural Research, NHLBI/NIH (K.Z.), NIH R35 GM127075 and R01 HD102474, and the Howard Hughes Medical Institute (X.C.).

## Materials & Correspondence

Correspondence and material requests should be addressed to X.C.

## Supplemental Information

### Materials and methods

#### Fly strains and husbandry

Flies were raised under standard yeast/molasses medium at 25 C° unless stated otherwise. The following flies were used: *nos-Gal4 (with VP16)/Cyo* (Van Doren et al., 1998), *nos-Gal4 (without VP16* or *ΔVP16)* on the 2^nd^ chromosome (from Yukiko Yamashita, University of Michigan, USA)*, UAS-set1 RNAi* (Bloomington *Drosophila* Stock Center, BL33704), *UAS-mCherry RNAi* (Bloomington *Drosophila* Stock Center, BL35785), *tj-Gal4/Cyo* (Tanentzapf et al., 2007), *bam-Gal4/TM6B* (Chen and McKearin, 2003)*, bam-Gal80* on the third chromosome (from Juliette Mathieu and Jean-René Huynh, Collège de France, France), *UAS-GFP.nls* (Bloomington *Drosophila* Stock Center, BL4775)*, UASp-FRT-EGFP-set1 (WT)-PolyA-FRT* (This study), *UASp-FRT-EGFP-set1 (WT rescue)-PolyA-FRT* (This study)*, UASp-FRT-EGFP-set1 (Mutant rescue)-PolyA-FRT* (This study)*, mad^12^, FRT40A/Cyo* (Bloomington *Drosophila* Stock Center, BL58785), *stat92E^06346^/ TM3, Sb* (Bloomington *Drosophila* Stock Center, BL11681), *vasa-GFP knock-in* (from Dr. Tatjana Trcek, Johns Hopkins University, Baltimore, MD).

#### Spatiotemporally controlled experiments

To study the function of Set1 in the *Drosophila* testis, two RNAi lines *UAS-set1 RNAi* and *UAS-mCherry RNAi* were crossed with different drivers *nos-Gal4/Cyo*, *bam-Gal4/TM6B*, and *tj-Gal4/Cyo* at 18 C°. Newly eclosed progenies were transferred to new vials at 25 C° and aged for 0,1, 3, 5, or 7 days before dissection.

For Set1 function study in the germline stem cells and gonialblast cells, two RNAi lines *UAS-set1 RNAi* and *UAS-mCherry RNAi* were crossed with *nos-Gal4ΔVP16/Cyo; bam-Gal80/MKRS* at 25 C°. Newly eclosed progeny were transferred to new vials and maintained at 25 ° for 7 days before dissection.

To determine the role of Set1 in the drosophila early germline at adulthood, two RNAi lines *vasa-GFP knock-in/CyO; UAS-set1 RNAi* and *vasa-GFP knock-in/CyO; UAS-mCherry RNAi* were crossed with *nos-Gal4/Cyo; tub-Gal80^ts^* at 18 C°. Newly eclosed progenies were transferred to new vials at 29 C° and aged for 0, 7, 14, 21, or 28 days before dissection. To study if expression of *EGFP-set1 cDNA* in the early germline is sufficient to rescue *nos>set1 RNAi* germ cell phenotypes, flies with the following genotypes: *nos-Gal4/ UASp-FRT-EGFP-set1 (WT rescue)-PolyA-FRT; UAS-set1 RNAi/+, nos-Gal4/ UASp-FRT-EGFP-set1 (Mutant rescue)-PolyA-FRT; UAS-set1 RNAi/+, nos-Gal4/ UASp-FRT-EGFP-set1 (WT)-PolyA-FRT; UAS-set1 RNAi/+, nos-Gal4/ UAS-GFP.nls; UAS-set1 RNAi/+* were grown at 25 C° and dissected at 1 and 5 days post eclosion.

To identify if *set1* genetically interacts with *mad* and *stat92E*, the alleles *stat92E^06346^*, and *mad^12^* were used. Flies with the following genotypes: *nos-Gal4/+; UAS set1 RNAi/ stat92E^06346^*, *UAS set1 RNAi/stat92E^06346^, nos-Gal4/mad^12^; UAS-set1 RNAi/+*, *nos-Gal4/mad^12^*, and *nos-Gal4/+; UAS set1 RNAi/+* were grown at 25 C° and dissected at 5 days post eclosion.

#### Generation of transgenic fly lines

For the lines *UASp-FRT-EGFP-set1 (WT rescue)-PolyA-FRT, UASp-FRT-EGFP-set1 (Mutant rescue)-PolyA-FRT*, the RNAi recognition sequence was altered. Since the RNAi recognition sequence is in the coding region of the Set1 gene: AAGGTGCAGAGTATAAGAGTA, the third base for each codon of this sequence was changed to make a silent mutation that would allow for the protein to be translated properly but for the RNAi to not recognize the mRNA. For the line *UASp-FRT-EGFP-set1 (Mutant rescue)-PolyA-FRT,* a mutation in the SET domain at G4713A to make the E1613K amino acid replacement was made.

#### Immunofluorescence

Testes were dissected in Schneider’s insect media and then fixed in 4% formaldehyde in 1X PBST (1X PBS with 0.1% Trition X-100) for 8 min at room temperature (RT). The testes were rinsed three times and washed three times for 5 min each time using 1X PBST at RT. Testes were incubated with primary antibodies in 1X PBST + 3% BSA at 4 C° for at least one night. Samples were then rinsed three times and washed three times for 5 min each time in 1X PBST and then incubated in 1:1,000 dilution of Alexa-Flour conjugated secondary antibody in 5% NGS + PBST for 2 hours at RT or rotating for a minimum of 24 hours at 4 C°. Samples were rinsed three times and washed three times, 5 minutes each in 1X PBST and then mounted for microscopy in vectashield antifade mounting medium (Cat#H-1400, Vector laboratories) with or without DAPI. Samples were imaged using a Leica SP8 or Stellaris 5 confocal microscope with a 63x oil immersion objective. Images were analyzed using imageJ software. Primary antibodies used are: Vasa (rabbit, 1:5,000, from R. Lehmann), Fas III (mouse, 1:50, DSHB, 7G10), Armadillo (mouse,1:50, DSHB, N2 7A1), Alpha -spec (mouse, 1:50, DHSB, 3A9), TJ (guinea pig, 1:1000, from M. Van Doren), H3K4me3 (rabbit, 1:400, cell signaling, 9751S), GFP (chicken, 1:1000, Abcam ab13970), Stat92E (rabbit, 1:200, gift from Denise Montell, University of Santa Barbara, CA, USA), and pMad (rabbit, 1:100, Abcam, ab52903).

#### RNA-seq and data analysis

Flies with the following genotypes: *nos-Gal4; UAS-mCherry RNAi* or *nos-Gal4; UAS-set1 RNAi* were collected as newly eclosed males and aged for 0, 1, 3, and 5D at 25 C° after shift from 18 C°. Approximately 15 pairs of testes for each genotype were dissected in schneiders media + 10% FBS as one replicate. Three replicates were generated for each time point and genotype. The testes were then desheathed in 500 μL of lysis buffer (tryspin LE + 2 mg/ml collagenase). The samples were incubated in lysis buffer for 10 min in a 37 C° water bath with gentle vortex mixing every 2 min. The samples were then filtered through a 40 μm tissue culture filter followed by a 10 min centrifugation at 1,200 rpm. The cells were washed with 200 ul of PBS and pelleted again for 5 min at 1,200 rpm. Total RNA was purified following the manufacturers instruction of the Quick-RNA Microprep Kit (R1050, Zymo Research corporation). The libraries were generated using the reagents provided in NEBNext Ultra II Directional RNA library Prep Kit for Illumina (E7760S, New England Biolabs Inc) and NEB Next Poly(A) mRNA Magnetic Isolation Module (E7490, New England Biolabs inc). The illumina compatible libraries were sequenced with Illumina Novaseq6000 sequencer at the National Institutes of Health sequencing facility.

The sequencing reads were examined using the FastQC quality software (Galaxy Version 0.73+galaxy0) (Andrews). Reads that passed the quality filter were then mapped to the *Drosophila* genome (D. melanogaster Aug 2014 (BDGP Release 6 + ISO11 MT/dm6)) using Bowtie 2 version 2.5.1 (Langmead et al., 2012). For gene mapping, the gene model (Drosophila_melanogaster.BDGP6.87.gtf) was utilized, and we followed an RNA-seq data analysis tutorial using Galaxy, as detailed in the work (Batut et al., 2018; Bérénice Batut; Hiltemann et al., 2023). Aligned reads were summarized to genes using featureCounts (Liao et al., 2014) with gene features obtained from NCBI RefSeq dm6 assembly. Raw counts were normalized to TPM counts. Principal component analysis (PCA) was performed on 16,610 genes with detectable expression in at least one sample, based on which sample distance matrix was computed. To assess differential gene expression between Set1 knockdown and controls, DESeq2 (Love et al., 2014) was applied on samples from each day, using the Benjamini and Hochberg method for multiple testing correction with default DESeq2 parameters. A threshold of log_2_ fold change of 1.3 and adjusted *P*< 0.05 was applied to ascertain differentially expressed genes, for which EnrichR (Chen et al., 2013) was applied to assess enrichment of molecular pathways in each set. All statistical analysis was conducted using R software version 4.2.0.

#### Phenotype Quantification

All phenotypic quantification was done in Fiji (ImageJ). GSC number was determined by counting every cell that was either Vasa positive or Tj negative and directly next to the hub. Early cyst cell number was quantified by counting every Tj positive cell in a whole testis z stack. Hub area quantification was determined by drawing a line around the z slice with the largest hub size and using the area measurement in Fiji.

#### Statistics and reproducibility

Data was subjected to the Shapiro-Wilk test to determine whether the data was normally distributed or skewed. For normally distributed data, an unpaired two sample t test was used to compare two individual datasets to each other. For skewed data, the Wilcoxon signed rank test was used to compare two individual datasets to each other. Data are presented with error bars representing the mean ± SE (standard error). Significant differences based on these statistical analysis were noted by asterisks (\**P*< 0.05, \*\**P*< 10^-2^, \*\*\**P*< 10^-3^, \*\*\*\**P*< 10^-4^).

### Supplemental figures and figure legends

**Figure S1:**
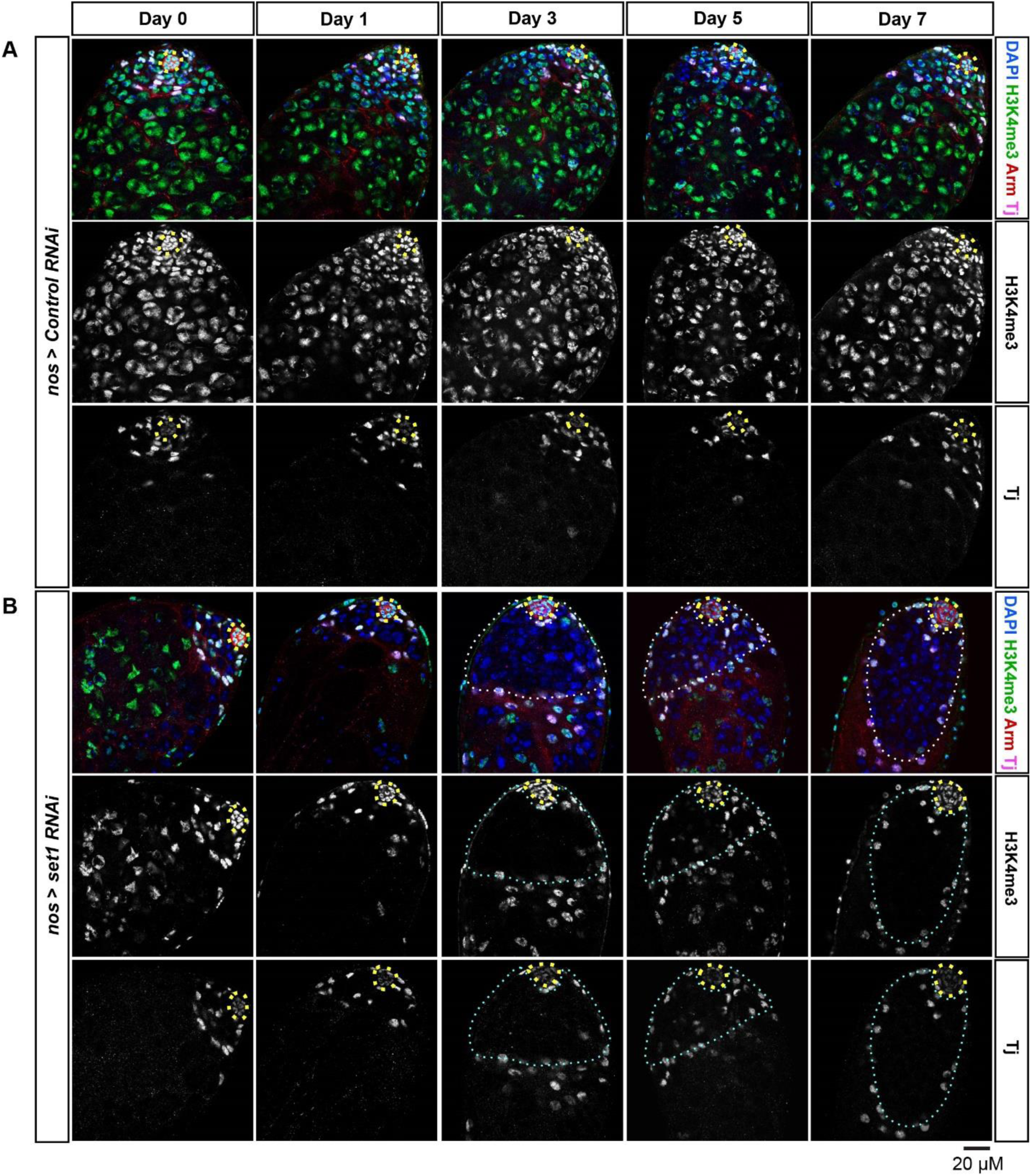
H3K4me3 in the *Drosophila* testis is dependent on Set1. Representative images of *nos>Control RNAi* testes (**A**) and *nos>Set1 RNAi* testes (**B**) at day 0, 1, 3, 5 and 7 post eclosion immunostained with H3K4me3 (green), Arm (red) for the hub region (yellow dotted outline), and Tj (magenta) for the CySC lineage cells, DAPI (blue). Cyan dotted outlines: overpopulated early-stage germ cells. Scale: 20μm.

**Figure S2:**
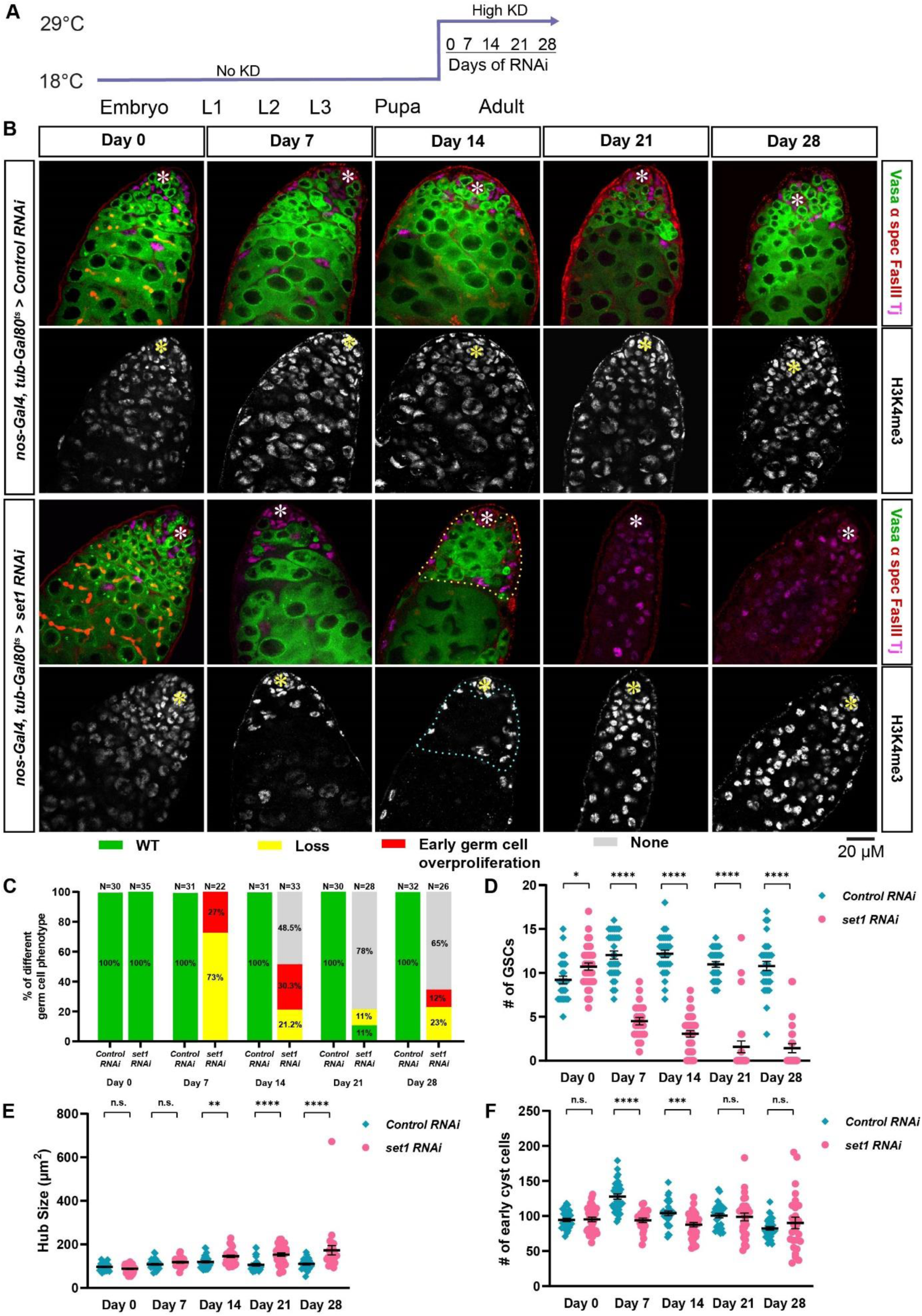
Knockdown of *set1* exclusively in the adult *Drosophila* testis leads to germ cell loss and germline differentiation defects. (**A**) *tub-Gal80^ts^, nos>Control RNAi* and *tub-Gal80^ts^, nos>set1 RNAi* flies are grown at the permissive temperature (18 C°) until eclosion, and shifted to the restrictive temperature (29 ° C) for 0, 7, 14, 21, and 28 days, respectively. (**B**) Representative images of *tub-Gal80^ts^, nos>Control RNAi* and *tub-Gal80^ts^, nos>set1 RNAi* testes at day 0, 7, 14, 21 and 28 post eclosion, *vasa-GFP* from a knock-in strain label the germline, immunostained with H3K4me3 (grey), Fas III (red) for the hub region, and Tj (magenta) for the CySC lineage cells. Asterisk: hub. White dotted outlines: overpopulated early-stage germ cells. (**C-D**) Quantification of the percentage of testes with the germline phenotypes (**C**) and GSC number in *tub-Gal80^ts^, nos>set1 RNAi* and *tub-Gal80^ts^, nos>Control RNAi* testes at 0, 7, 14, 21 and 28 days post eclosion, respectively: *tub-Gal80^ts^, nos>Control RNAi* testes (Day 0: N=30; Day 7: N=31; Day 14: N=31; Day 21: N=30; Day 28: N=32) and *tub-Gal80^ts^, nos>set1 RNAi* testes (Day 0: N=35; Day 7: N=22; Day 14: N=33; Day 21: N=28; Day 28: N=26). Refer to Table S1. (D) Quantification of hub region size for *tub-Gal80^ts^, nos>Control RNAi* and *tub-Gal80^ts^, nos>set1 RNAi* testes. Refer to Table S2. (**F**) Quantification of early cyst cell number for *tub-Gal80^ts^, nos>Control RNAi* and *tub-Gal80^ts^, nos>set1 RNAi* testes. Refer to Table S3. Individual data points and mean values are shown. Error bars represent SEM. \*\*\*\**P*< 10^-4^, \*\*\**P*< 10^-3^, \*\**P*< 10^-2^, \**P*< 0.05, n.s.: not significant; unpaired t test to compare two individual datasets with each other.

**Figure S3:**
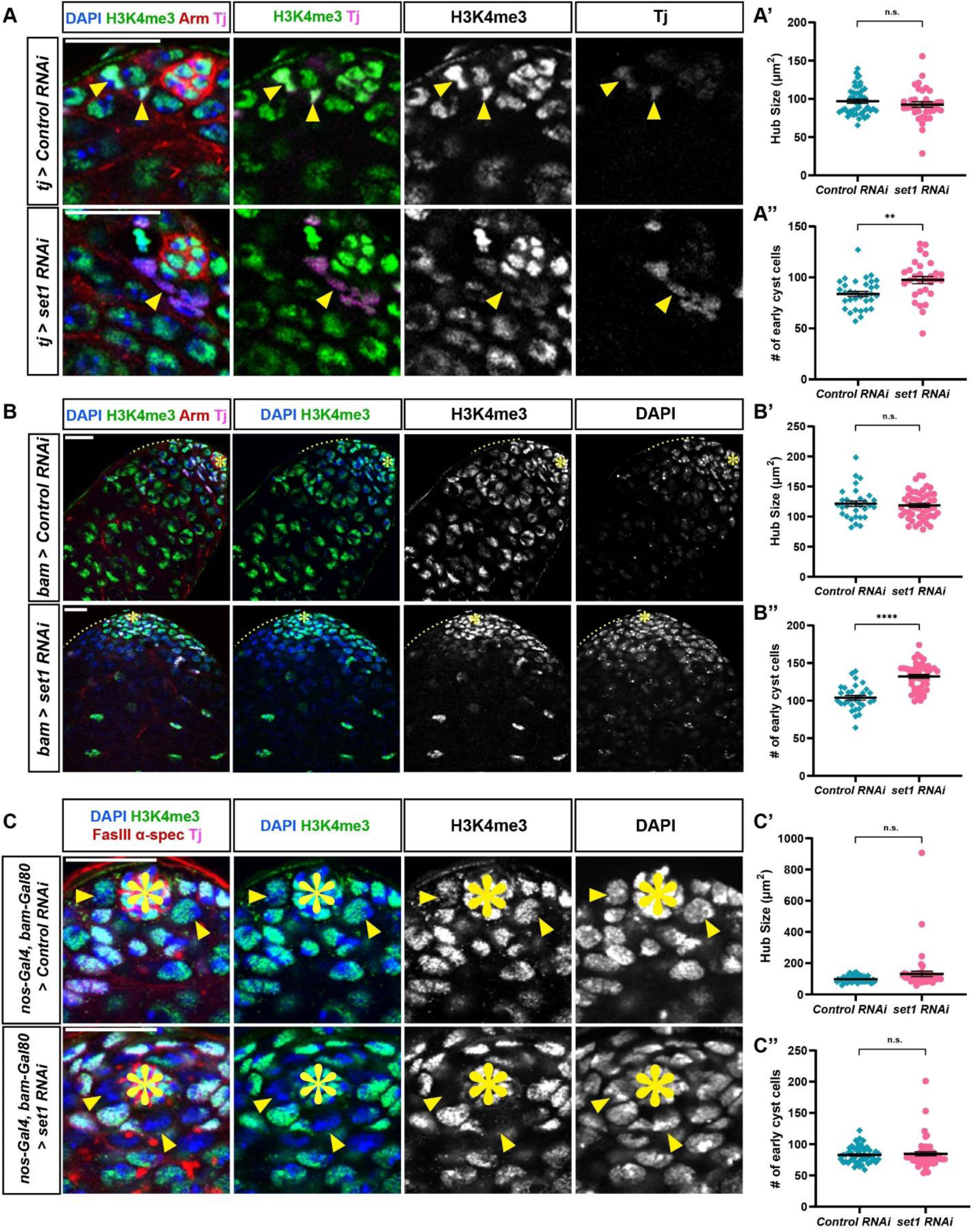
H3K4me3 signals are reduced in the somatic gonadal cells, late spermatogonial cells, and GSCs in the *Drosophila* testis by knockdown of *set1* using cell type- and stage-specific drivers. (**A**) Representative images of *tj>Control RNAi* and *tj>set1 RNAi* testes at day 7 post eclosion, immunostained with H3K4me3 (green), Arm (red) for the hub region, and Tj (magenta) for the CySC lineage cells. (**A’**) Quantification of the hub region size for *tj>Control RNAi* testes (N= 30) and *tj>set1 RNAi* testes (N= 30). Refer to Table S2. (**A’’**) Quantification of cyst cell number for *tj>Control RNAi* and *tj>set1 RNAi* testes. Individual data points and mean values are shown. Refer to Table S3. (**B**) Representative images of *bam>Control RNAi* and *bam>set1 RNAi* testes at day 7 post eclosion, immunostained with H3K4me3 (green), Arm (red) for the hub region, and Tj (magenta) for the CySC lineage cells. (**B’**) Quantification of the hub size for *bam>Control RNAi* testes (N= 31) and *bam>set1 RNAi* testes (N= 50). Refer to Table S2. (**B’’**) Quantification of cyst cell number for *bam>Control RNAi* and *bam>set1 RNAi* testes. Refer to Table S3. (**C**) Representative images of *nos-Gal4ΔVP16, bam-Gal80 >Control RNAi* and *nos-Gal4ΔVP16, bam-Gal80 >set1 RNAi* testes at day 7 post eclosion, immunostained with H3K4me3 (green), Arm (red) for the hub region, and Tj (magenta) for the CySC lineage cells. (**C’**) Quantification of hub region size for *nos-Gal4ΔVP16, bam-Gal80 >Control RNAi* testes (N= 52) and *nos-Gal4ΔVP16, bam-Gal80 >set1 RNAi* testes (N= 52). Refer to Table S2. (**C’’**) Quantification of cyst cell number for *nos-Gal4ΔVP16, bam-Gal80>Control RNAi* and *nos-Gal4ΔVP16, bam-Gal80>set1 RNAi* testes. Refer to Table S3. (**A’-A”**, **B’-B”**, **C’-C”**) Individual data points and mean values are shown. Error bars represent SEM. \*\*\*\**P*< 10^-4^, \*\**P*< 10^-2^, n.s.: not significant, unpaired t test to compare two individual datasets to each other. Asterisk in (**B** and **C**): Hub region. Yellow arrow heads: cyst cells in (**A**); GSCs in (**C**). Yellow dotted line: 4-16 spermatogonial cell region in (**B**). Scale: 20 μm.

**Figure S4:**
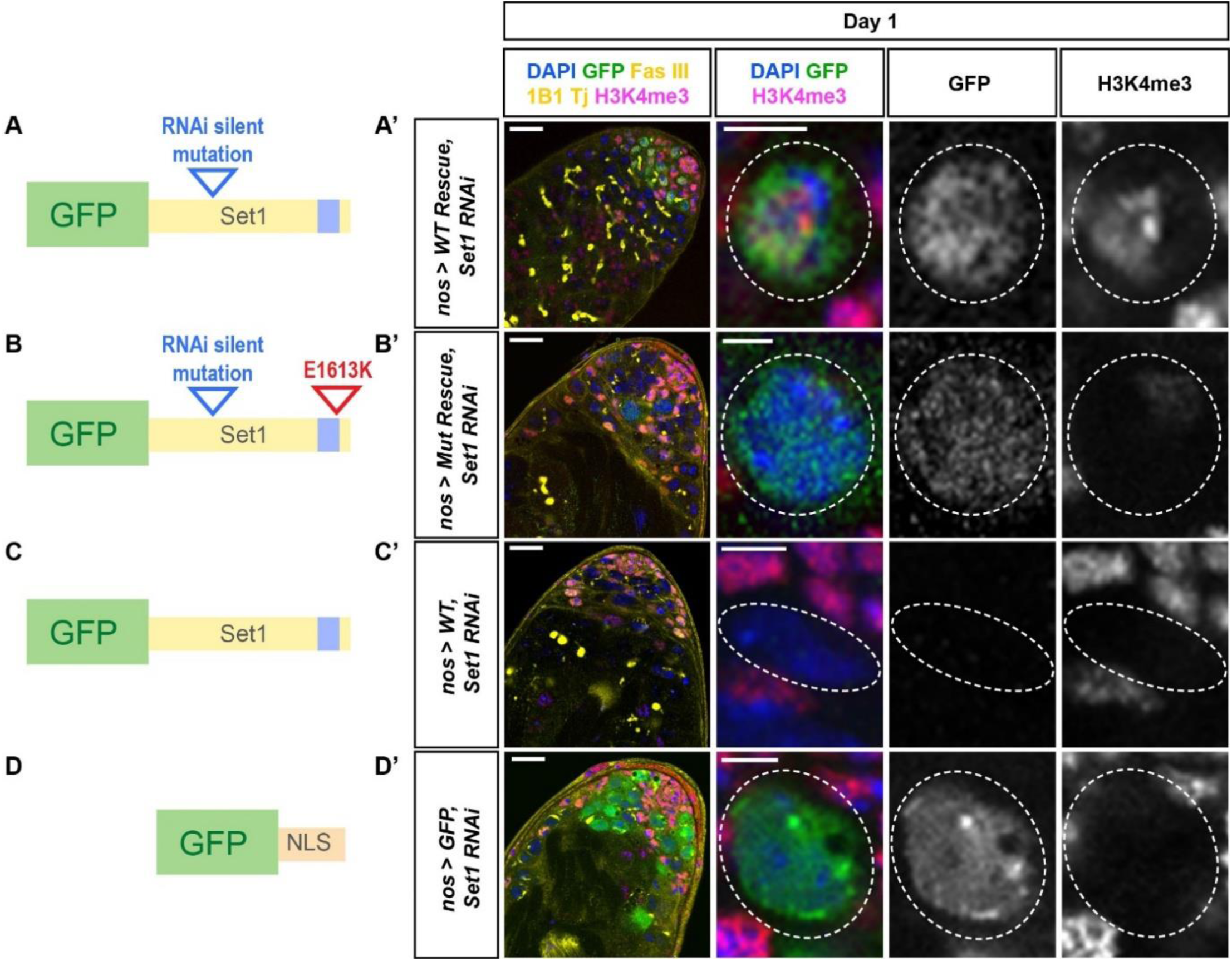
The methyltransferase activity of Set1 is required for its function in the male germline. (**A**-**D**) Cartoon depictions: the *set1* cDNA transgene with the RNAi recognition sequences mutated, named WT rescue (**A**); the *set1* cDNA transgene with the RNAi recognition sequences mutated and an E→K amino acid change in the SET domain, named Mut rescue (**B**); the *set1* cDNA transgene without the RNAi recognition sequences mutated, named WT (**C**); the *GFP* cDNA with the *nuclear localization* sequence, named GFP (**D**). Representative images of *nos-Gal4>WT Rescue, set1 RNAi* testis (**A’**), *nos-Gal4>Mut Rescue, set1 RNAi* testis (**B’**), *nos-Gal4>WT, set1 RNAi* testis (**C’**), *nos-Gal4>GFP, set1 RNAi* testis (**D’**), all testes are from males 1 day post eclosion, immunostained with H3K4me3 (magenta), GFP (green), Fas III (yellow) for the hub region, and Tj (yellow) for the CySC lineage cells. White dotted outline: germ cell expressing the corresponding transgenes. Scale: 20 μm for the testis image; 5 μm for individual germ cell images.

**Figure S5:**
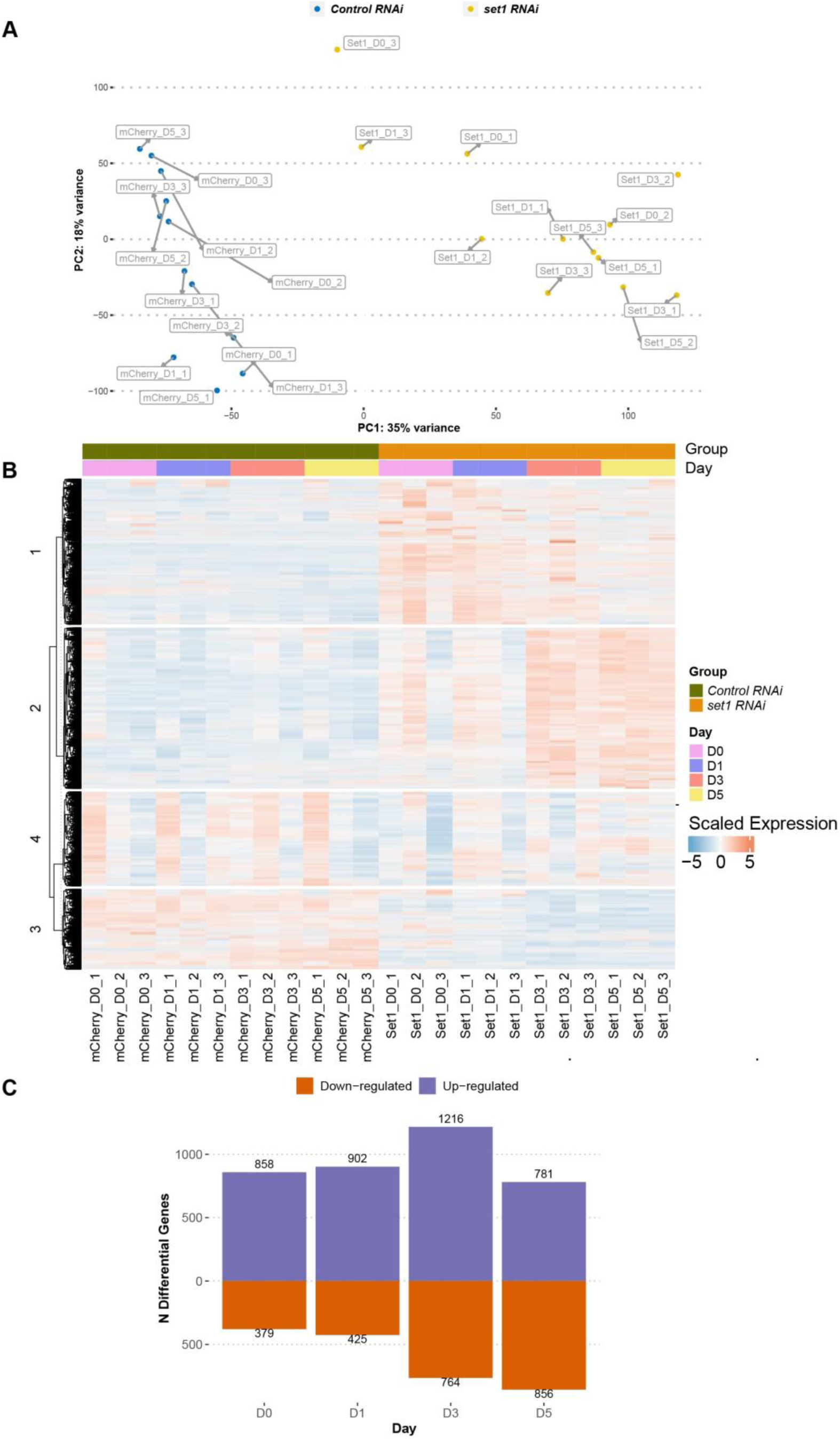
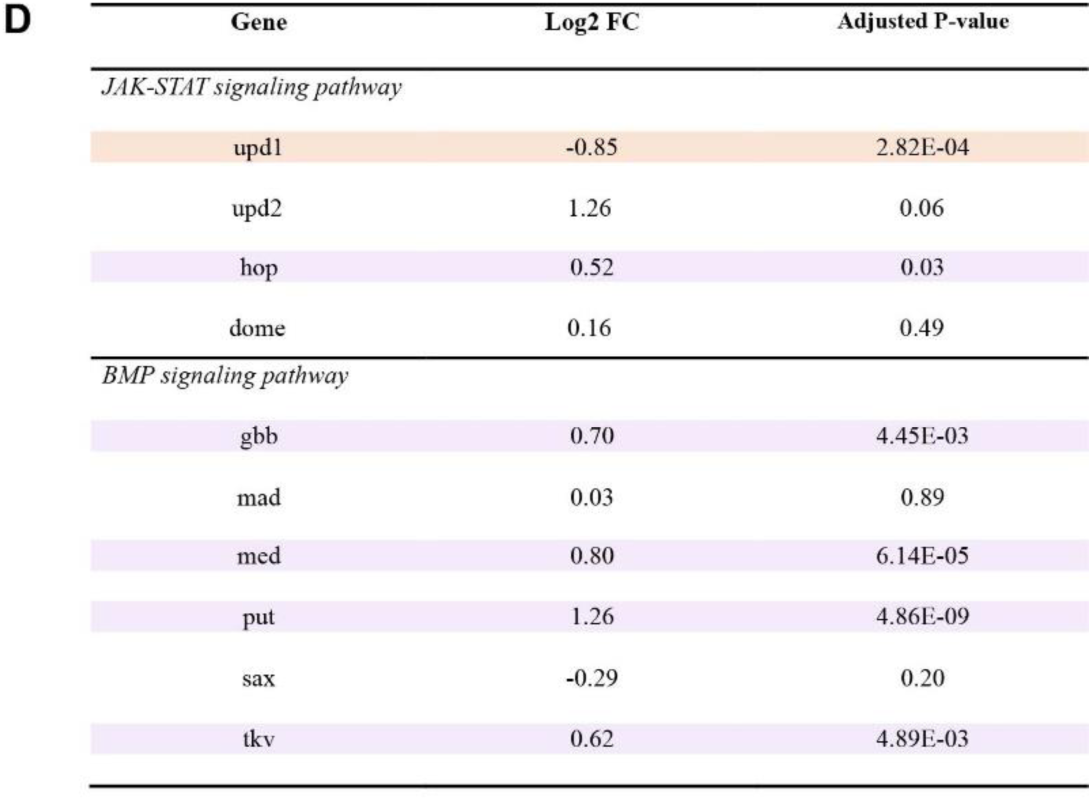
Knockdown of *set1* in the early germline results in global gene expression changes. (**A**) Principal component analysis plot shows multidimensional distribution of all *nos>set1 RNAi* (*set1* KD, 12 samples) and *nos>Control RNAi* (*Ctrl* KD, 12 samples) data sets: Three biological replicates for both *set1* KD and *Ctrl* KD at day 0, 1, 3, 5 post eclosion, respectively. There are two main clusters: a *set1* KD cluster and a *Ctrl* KD cluster. Within the *set1* KD cluster, Day 0 and Day 1 timepoint samples cluster while Day 3 and Day 5 timepoint samples cluster. (**B**) Heatmap shows differentially expressed genes (DEGs) between *set1* KD and *Ctrl* KD testes at day 0, 1, 3, 5 post eclosion, respectively. All genes are separated into four different groups. Group 1 contains the genes that are most expressed in *set1* KD testes at Day 0 and 1. Group 2 consists of the genes that are most expressed in *set1* KD testes at Day 3 and 5. Group 3 is comprised of genes more highly expressed in *Ctrl* KD testes and Group 4 is composed of the genes that did not fit any particular expression pattern with regards to genotype or timepoint. (**C**) Bar plot shows the number of DEGs with ≥1.3-fold (log_2_ scale) and *P*< 0.05 in *set1* KD testes compared to *Ctrl* KD testes at day 0, 1, 3, 5 post eclosion, respectively. (**D**) A table shows the detailed expression changes of several core components of the JAK-STAT and BMP signaling pathways at Day 3. Genes that are significantly upregulated are highlighted in purple and genes that were significantly downregulated are highlighted in orange.

**Figure S6:**
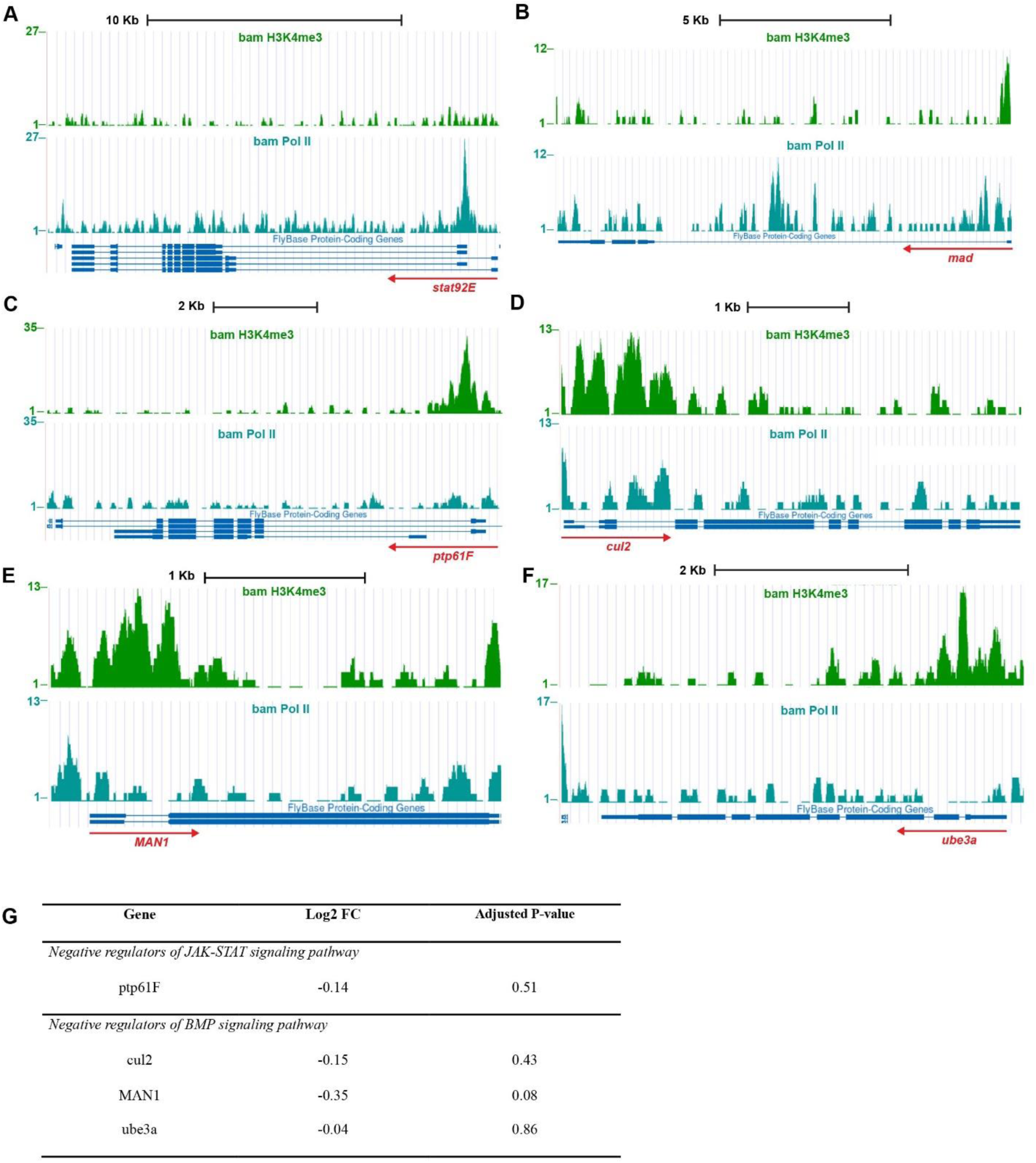
UCSC genome browser screenshots show H3K4me3 and RNA Pol II enrichment at individual gene regions. (**A**-**F**) All data are based on ChIP-seq using progenitor germ cell-enriched *bam* mutant testes (Gan et al., 2010), H3K4me3 (green) and RNA polymerase II (blue). (**A**) At the *stat92E* gene locus, H3K4me3 is not distinctly enriched at the promoter region even though Pol II profiles indicates active transcription. (**B**) At the *mad* gene locus, H3K4me3 is enriched at the promoter region and Pol II profiles indicates active transcription. (**C**) At the *ptp61F* gene locus, H3K4me3 is very distinctly enriched at the promoter region, even though Pol II profiles indicates inactive transcription. (**D**) At the *cul-2* gene locus, H3K4me3 is enriched at the promoter region and Pol II profiles indicates active transcription. (**E**) At the *MAN1* gene locus, H3K4me3 is enriched at the promoter region and Pol II profiles indicates active transcription. (**F**) At the *ube3a* gene locus, H3K4me3 is very distinctly enriched at the promoter region, even though Pol II profiles indicates inactive transcription. (**G**) A table shows the detailed expression changes of several inhibitors of the JAK-STAT and BMP signaling pathways at Day 3.

### Supplemental Tables

**Table S1:**
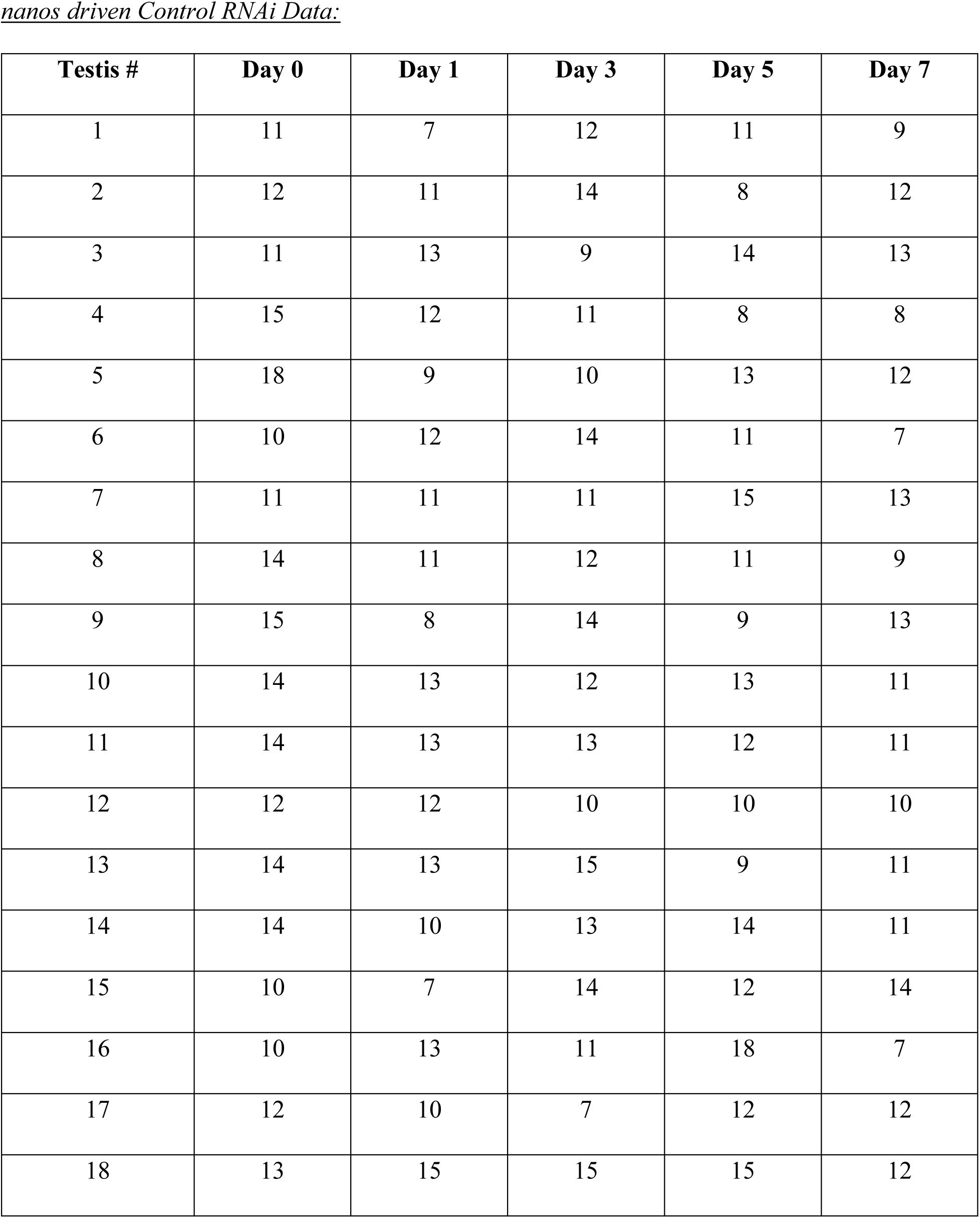
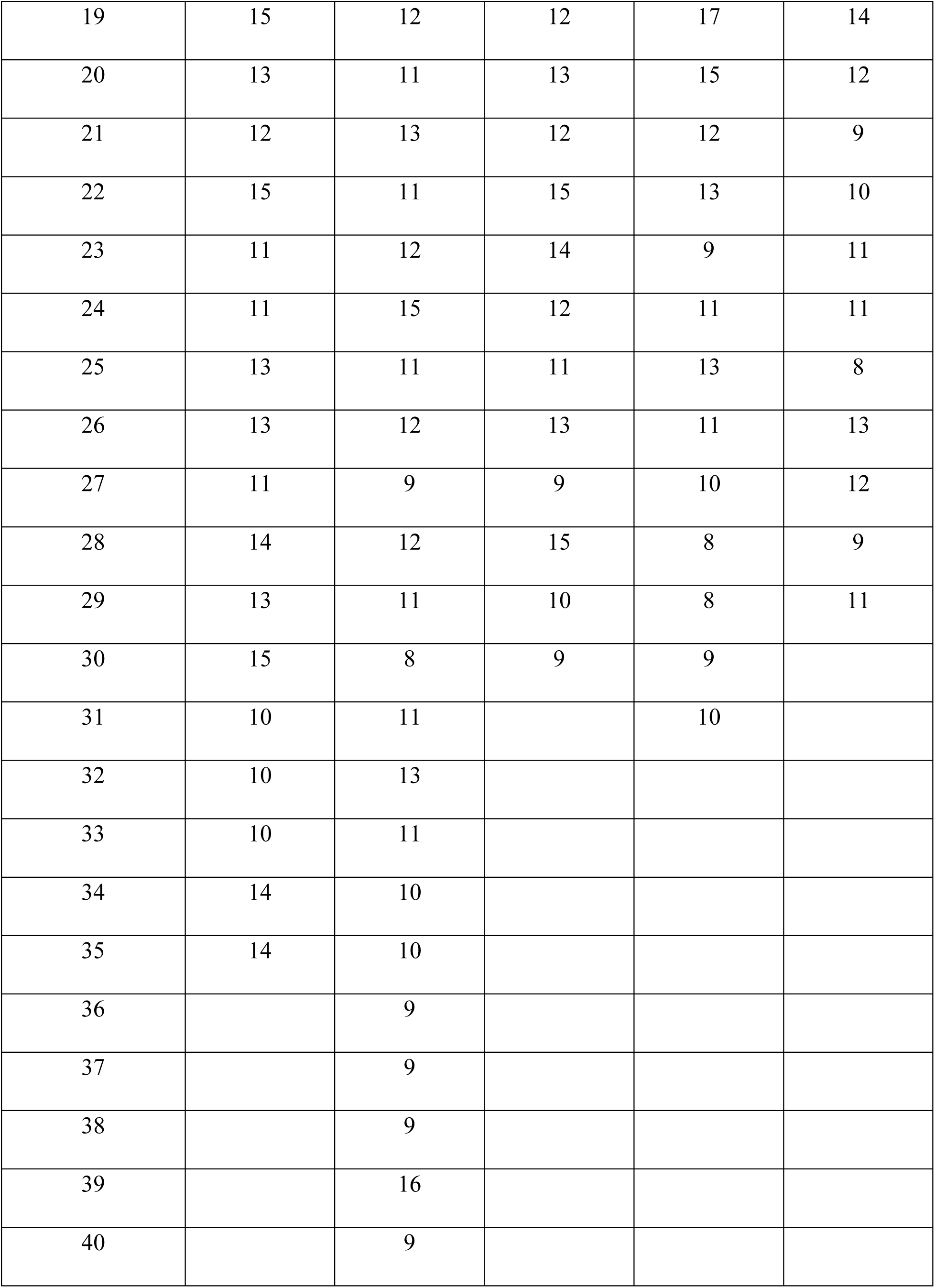
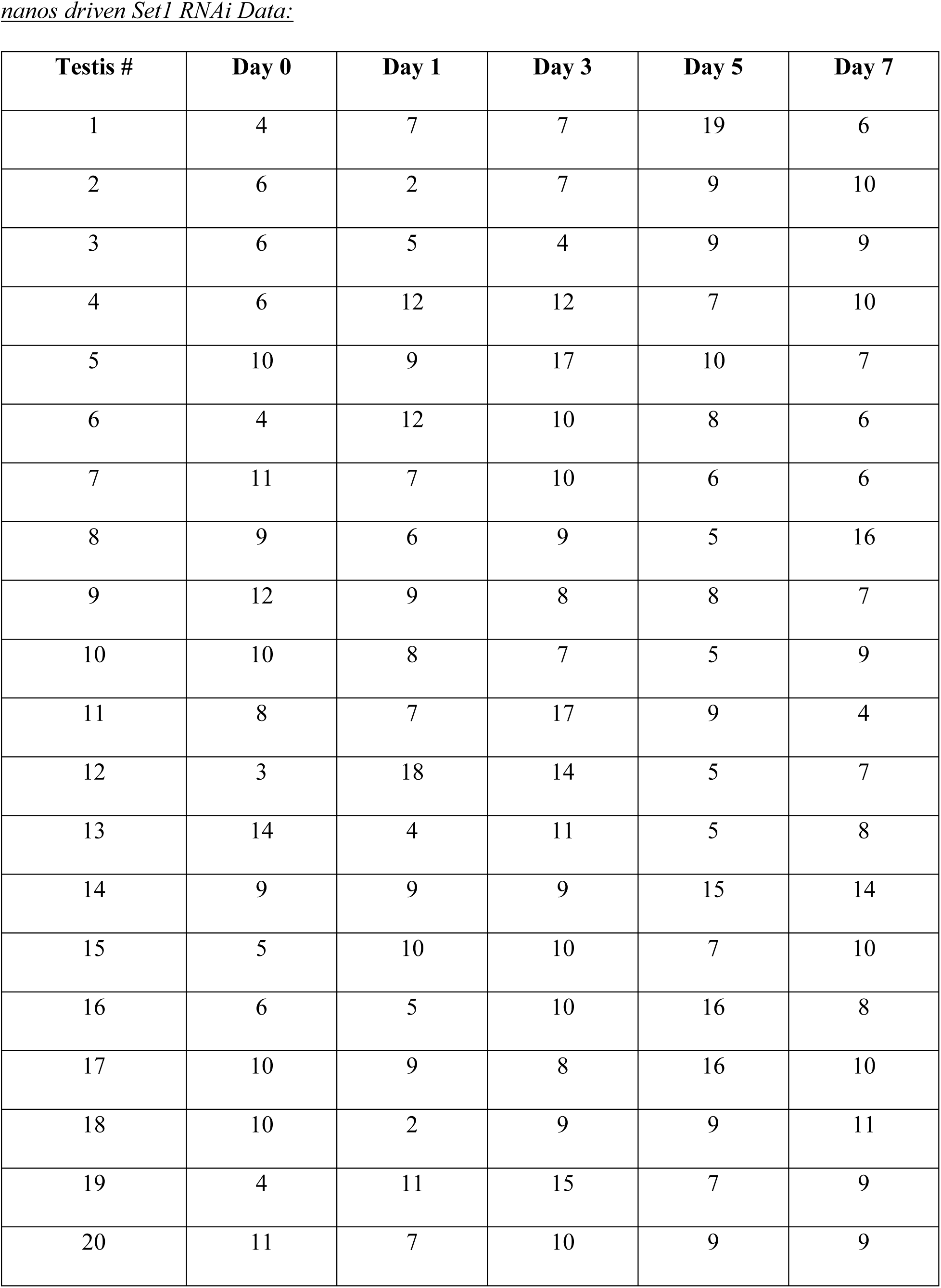
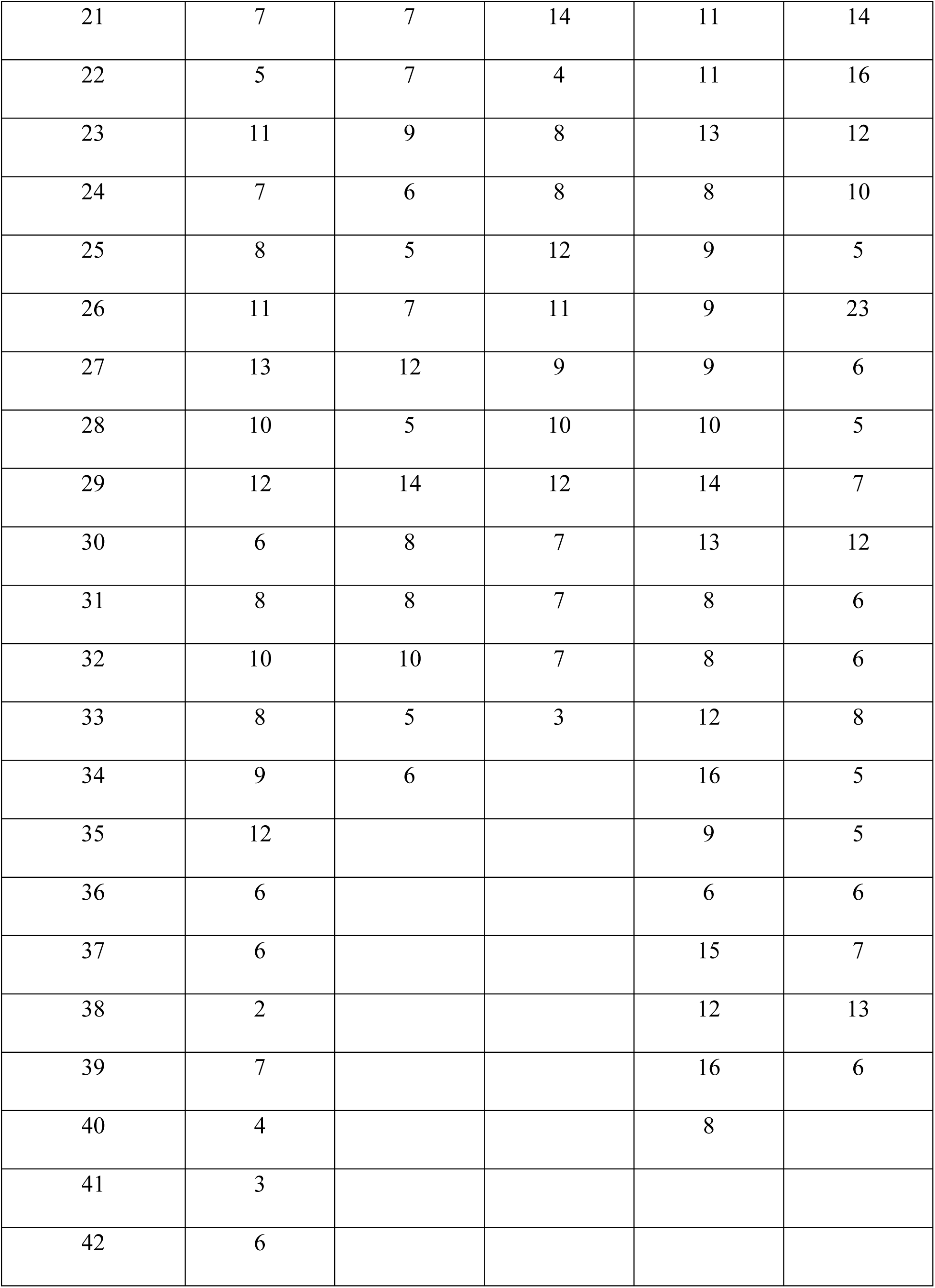
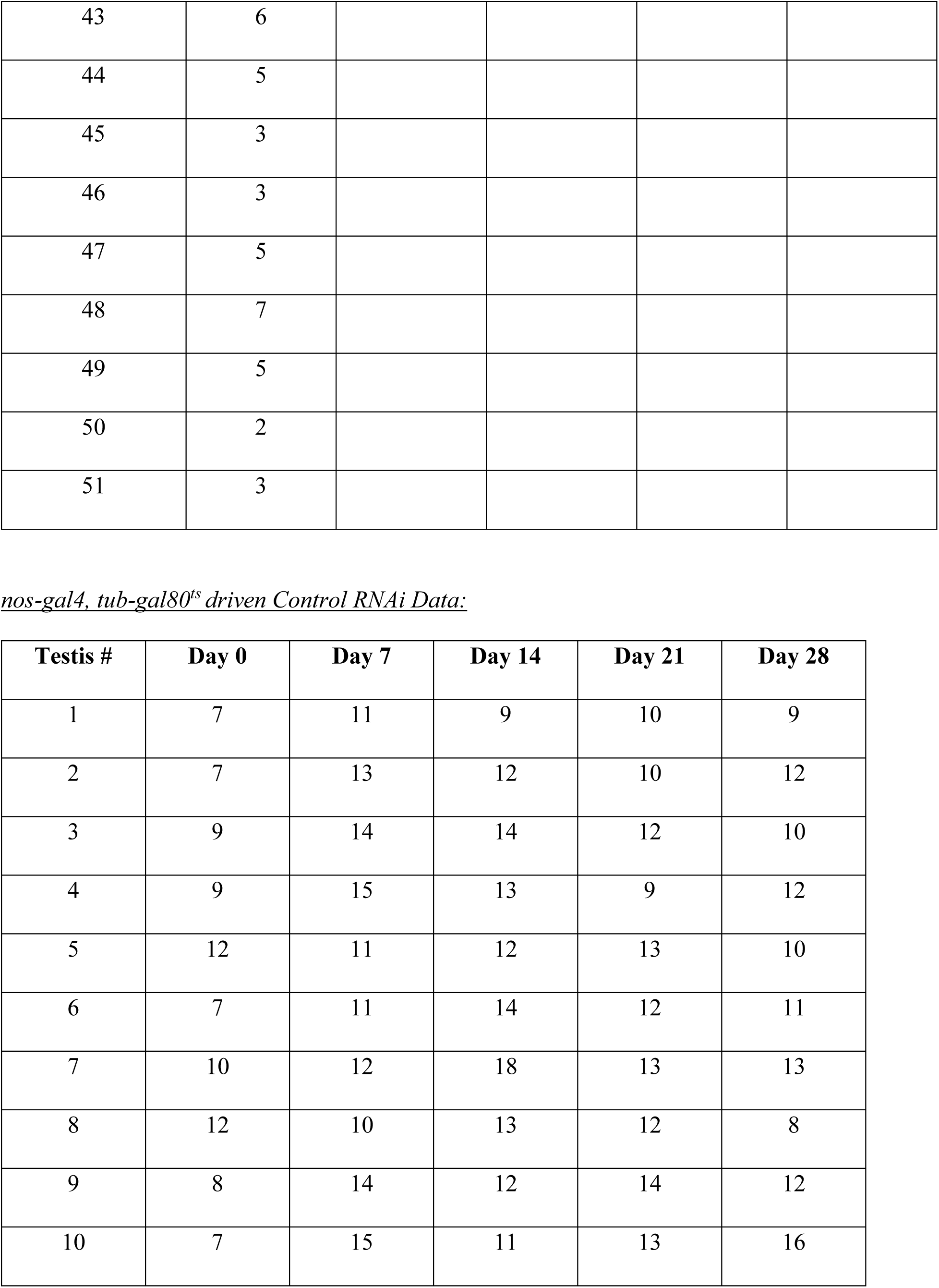
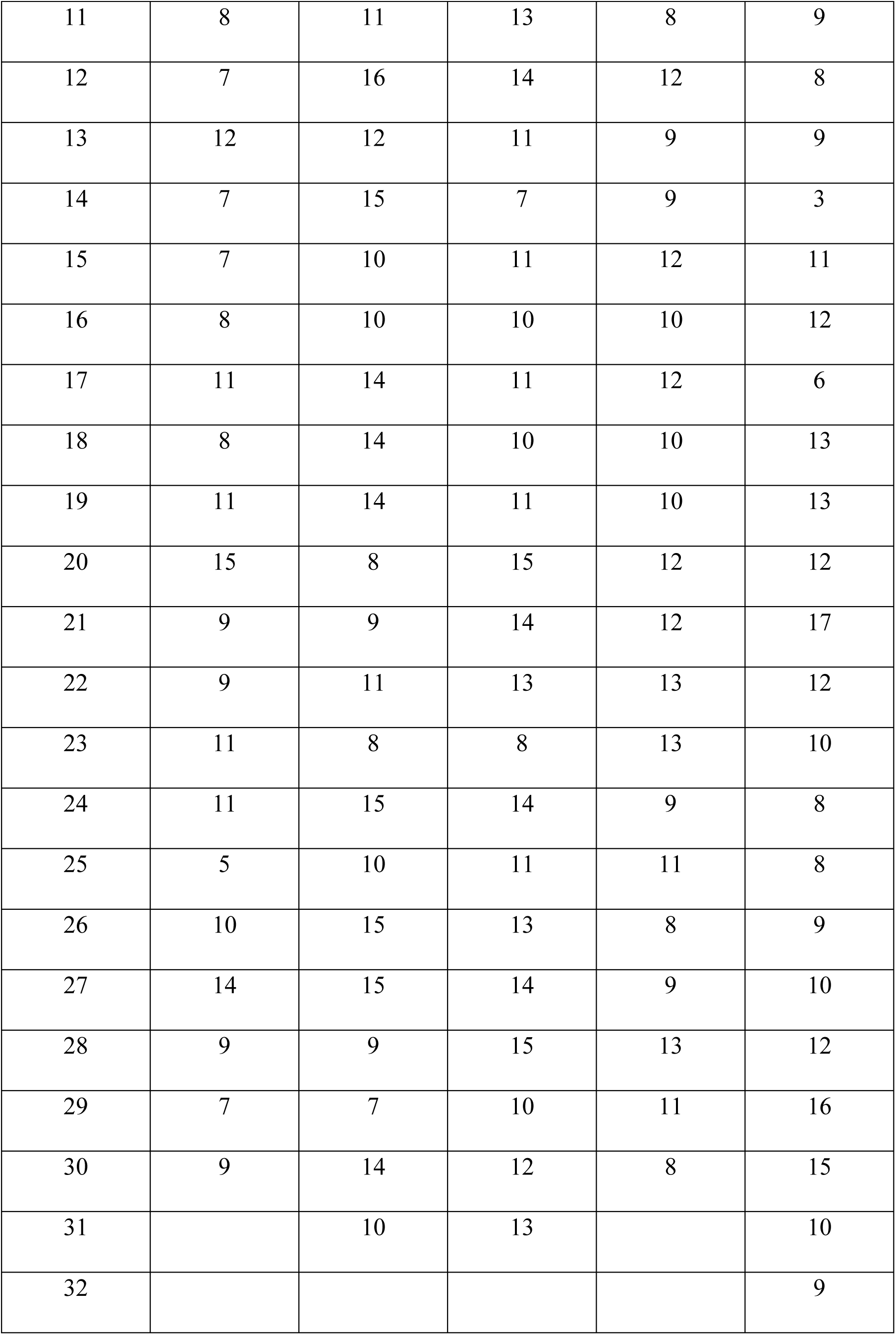
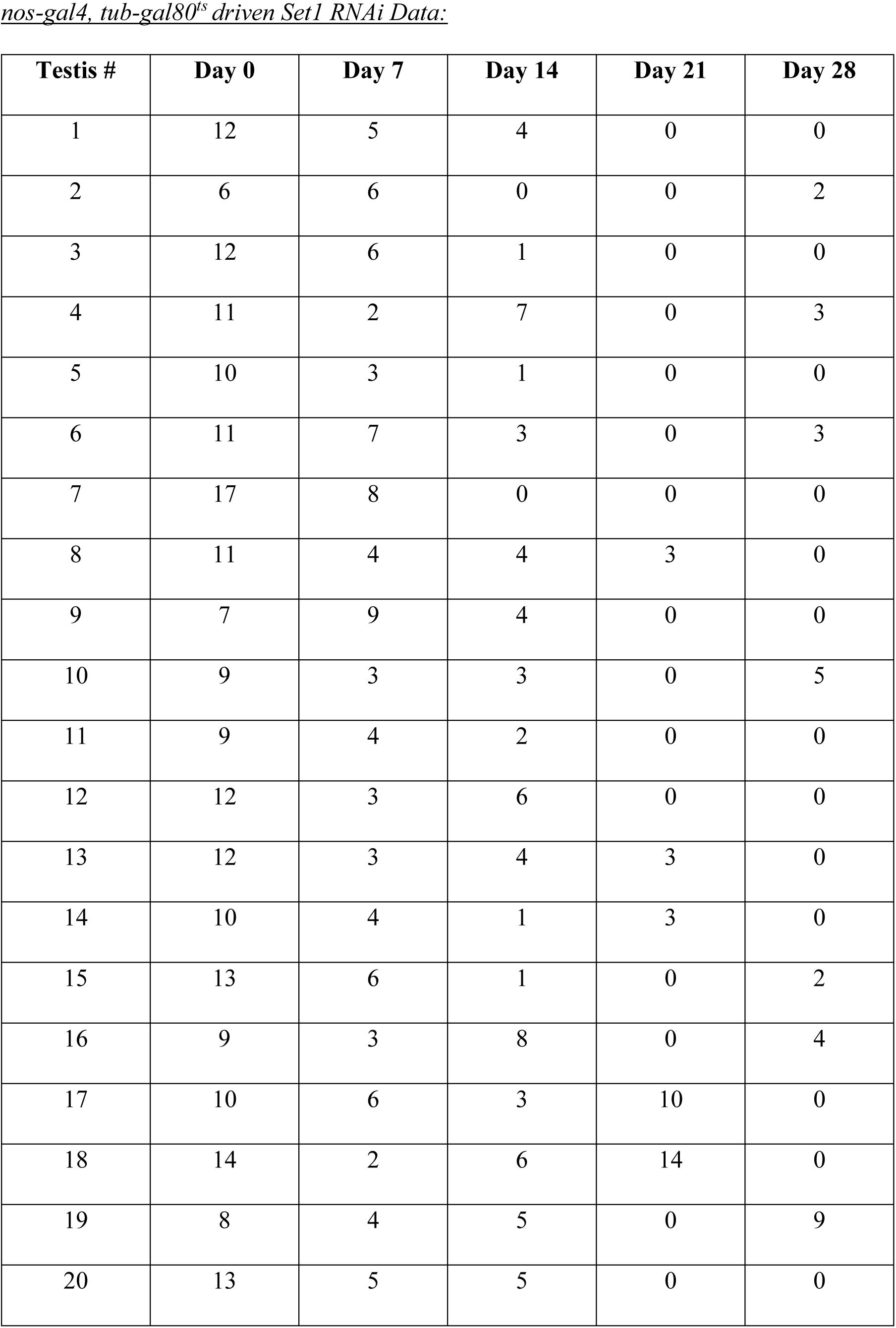
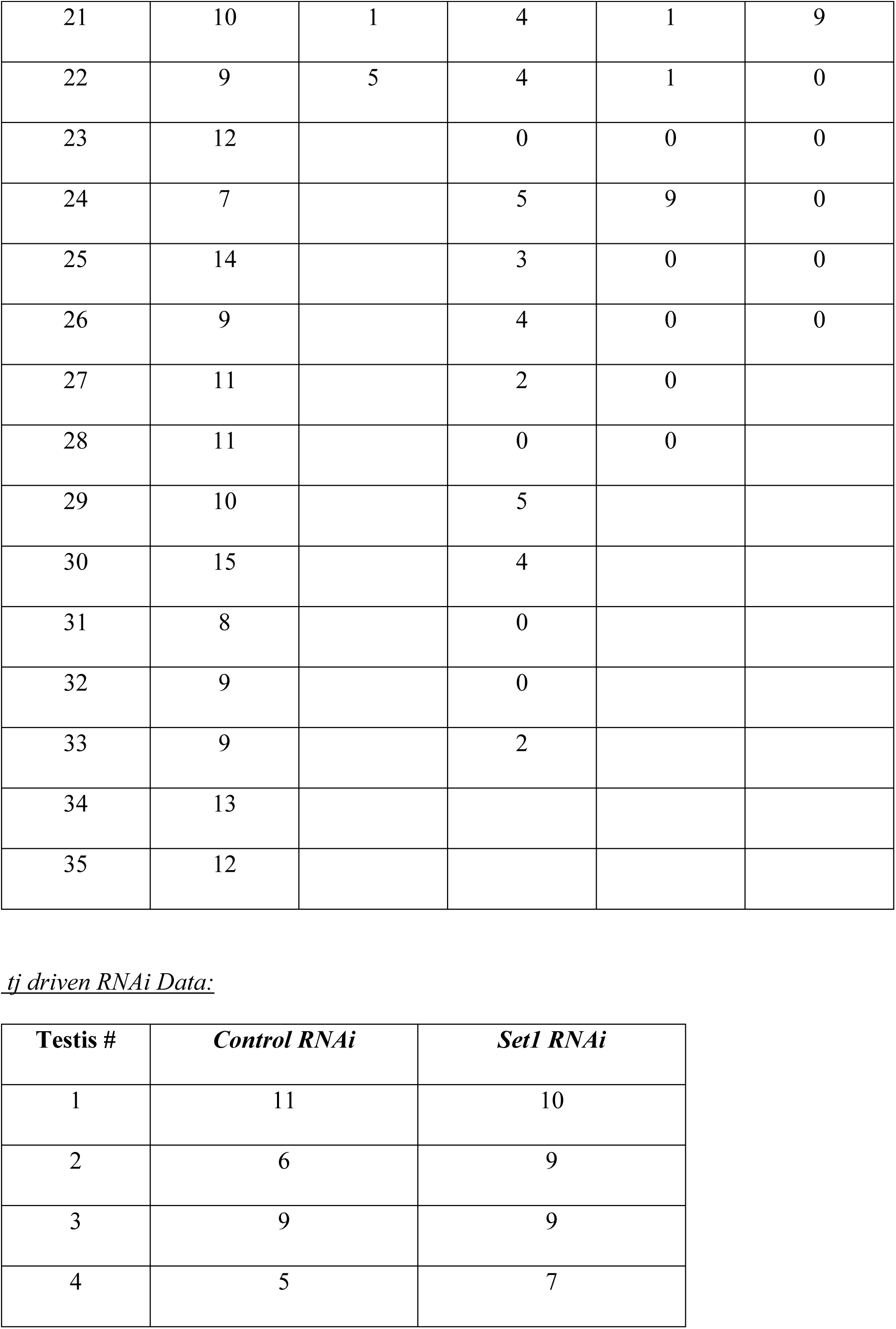
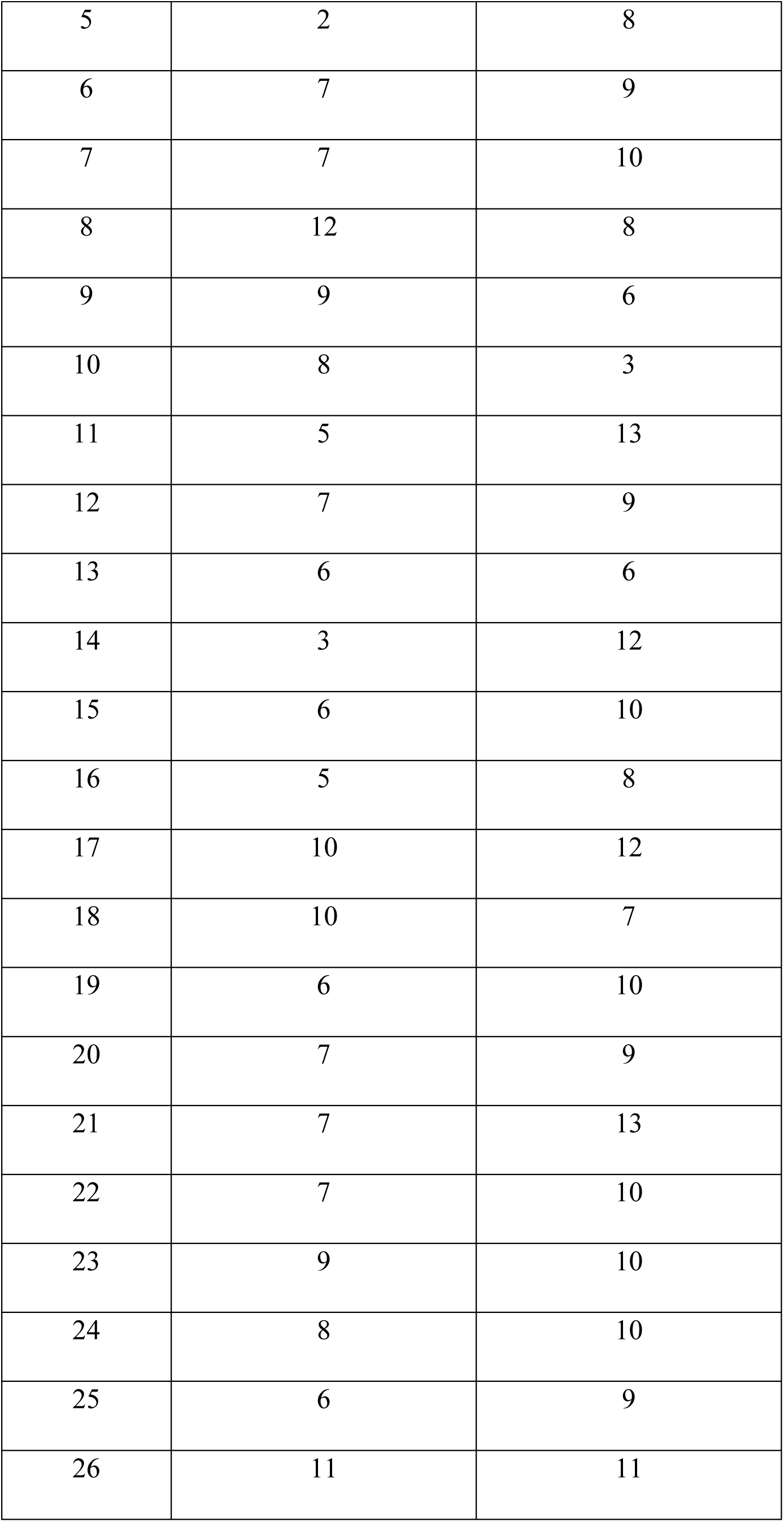
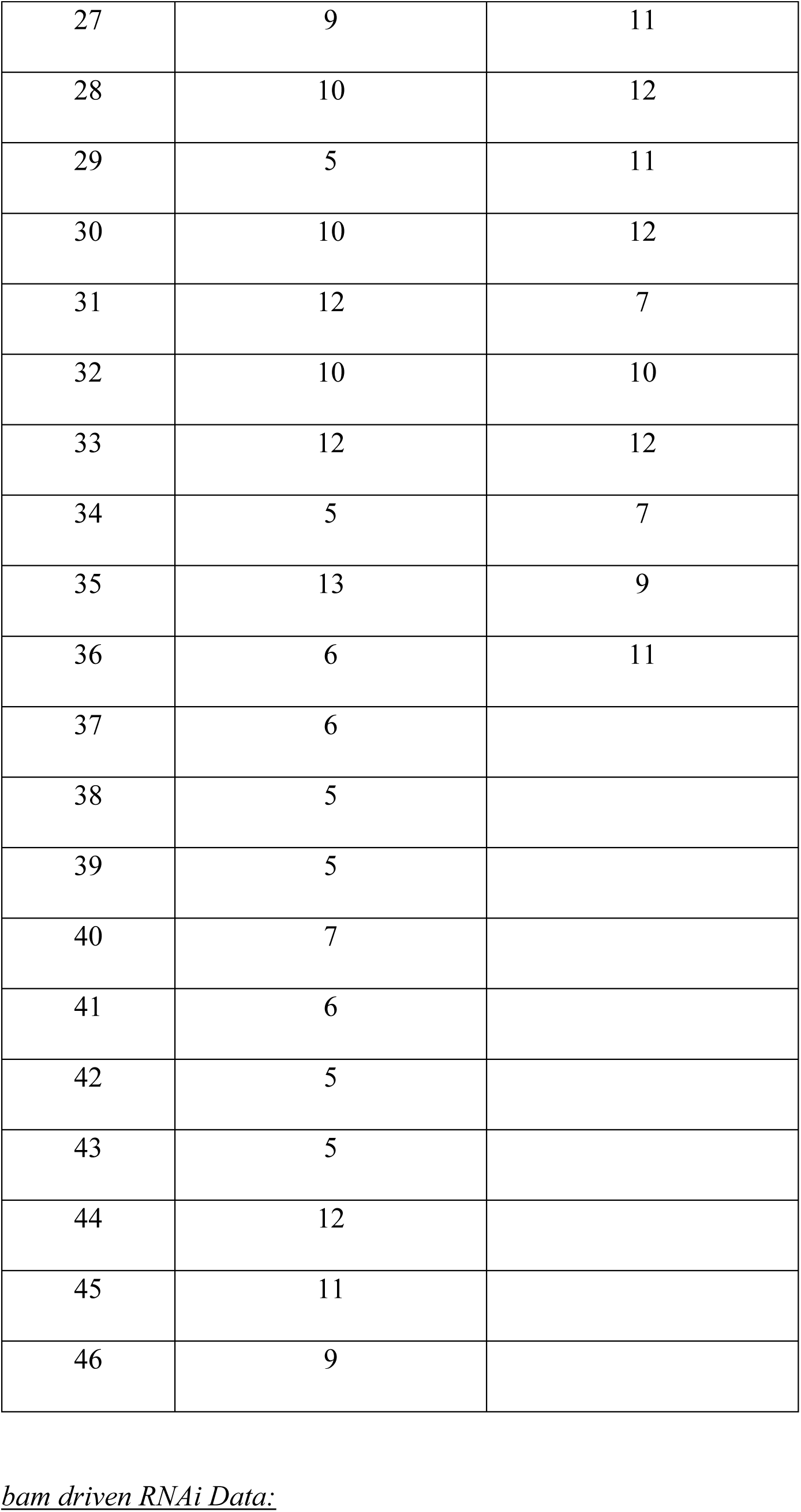
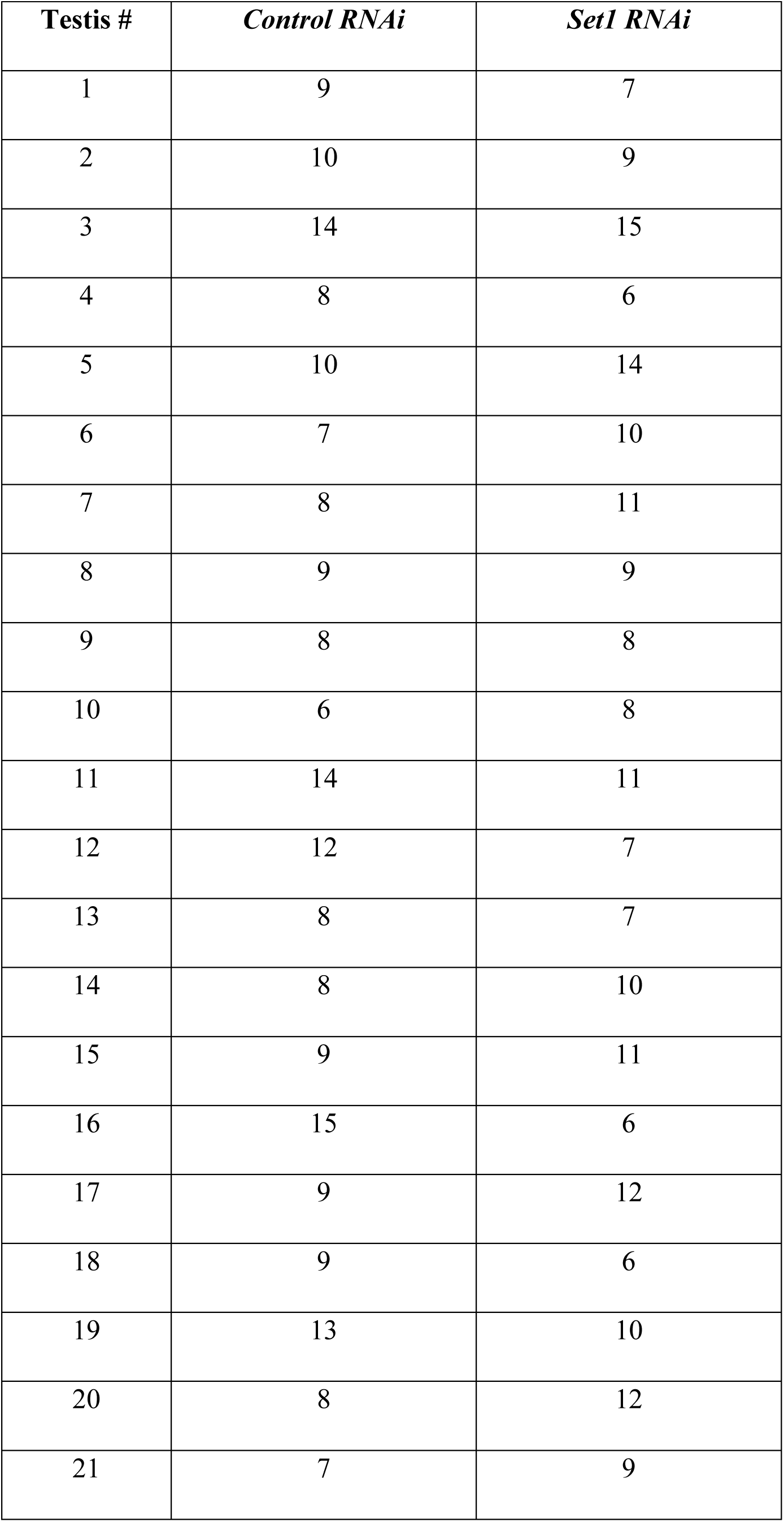
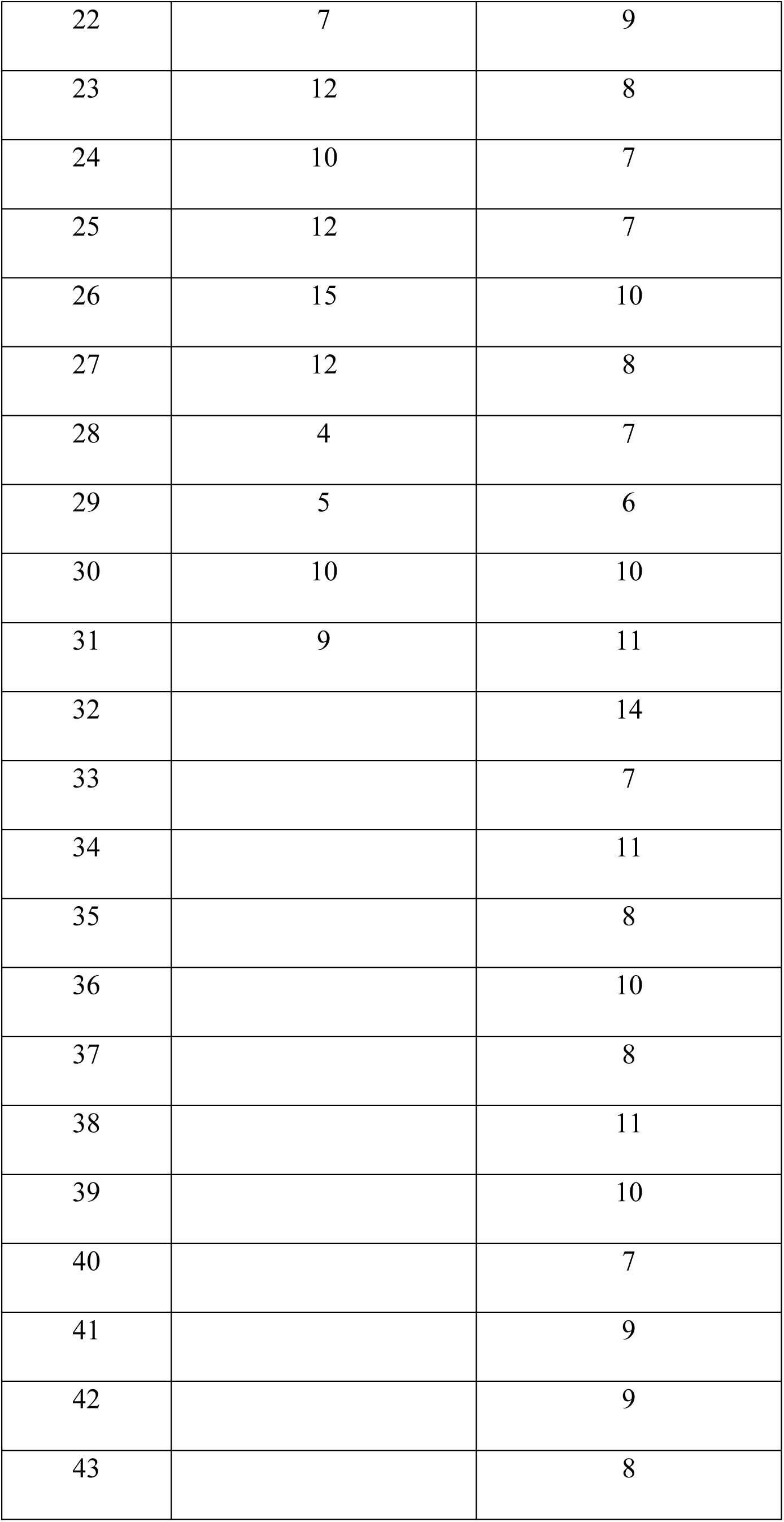
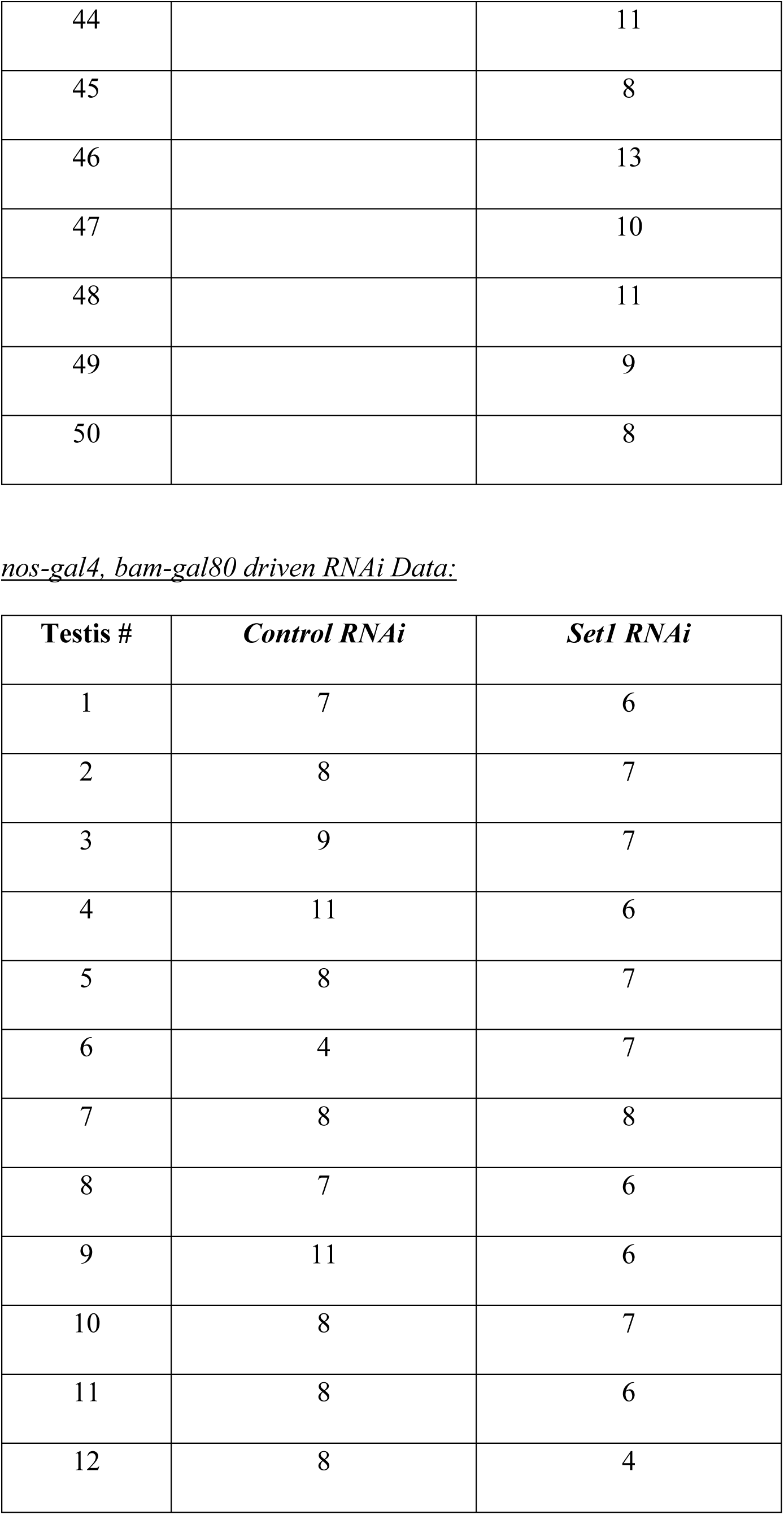
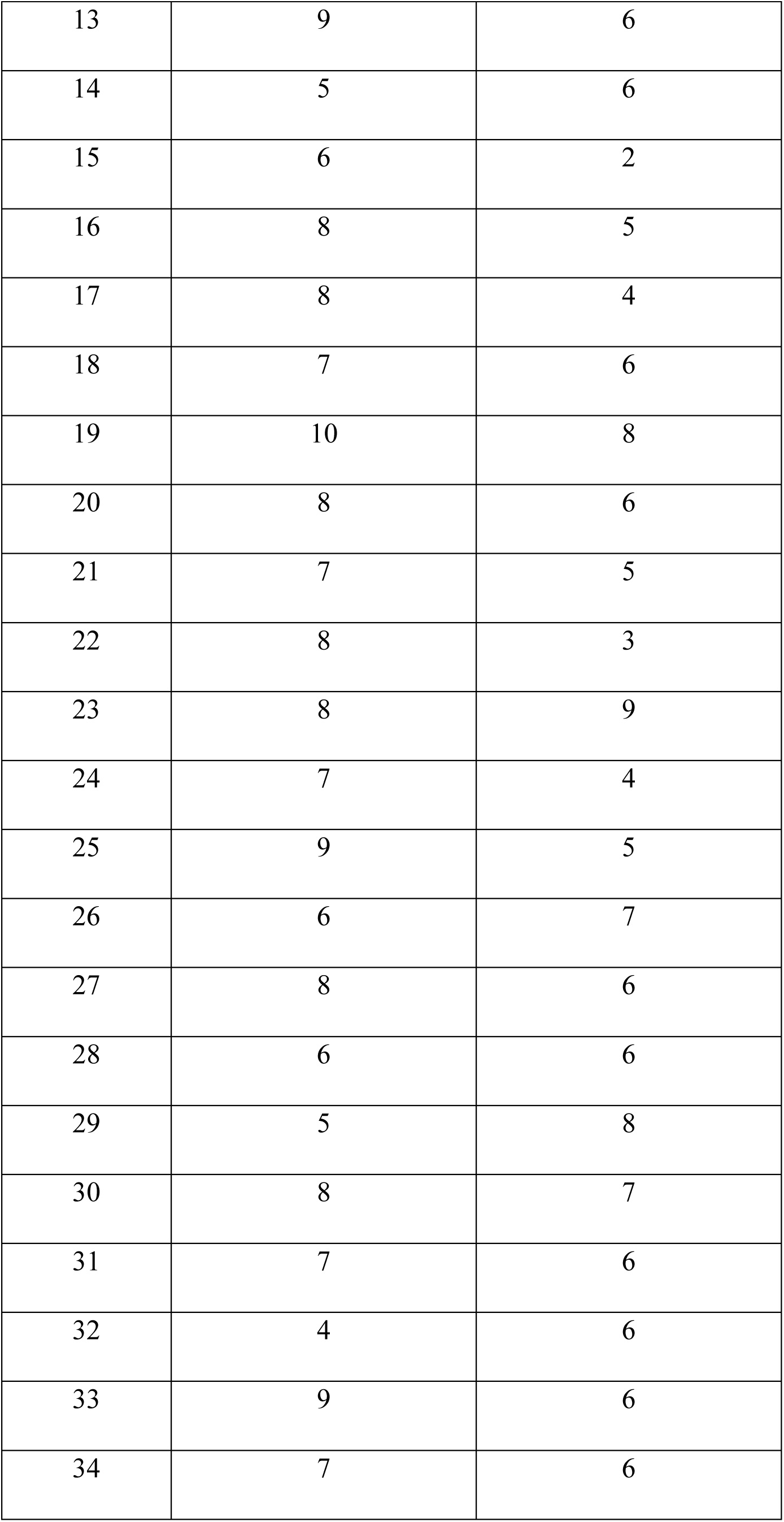
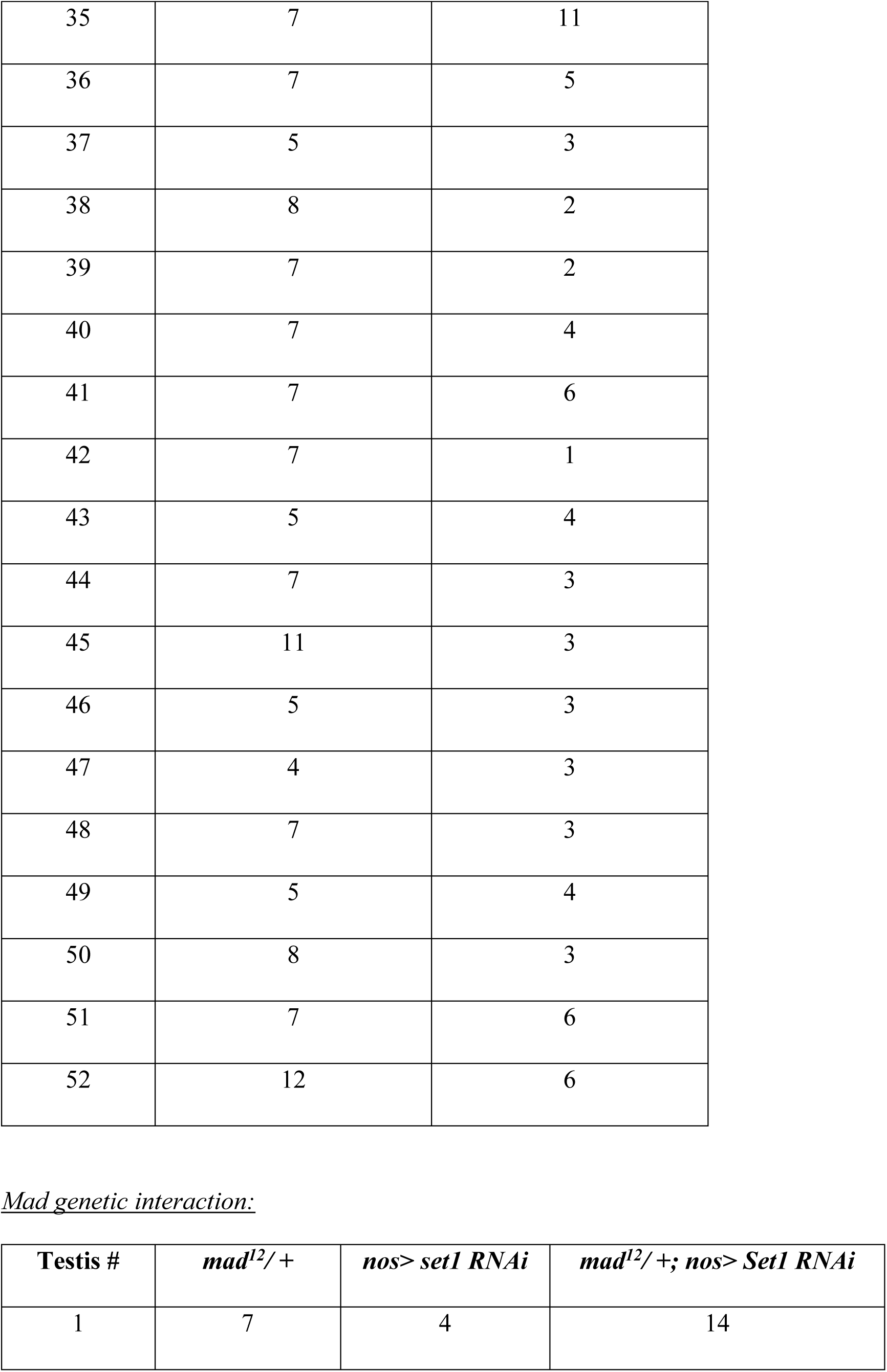
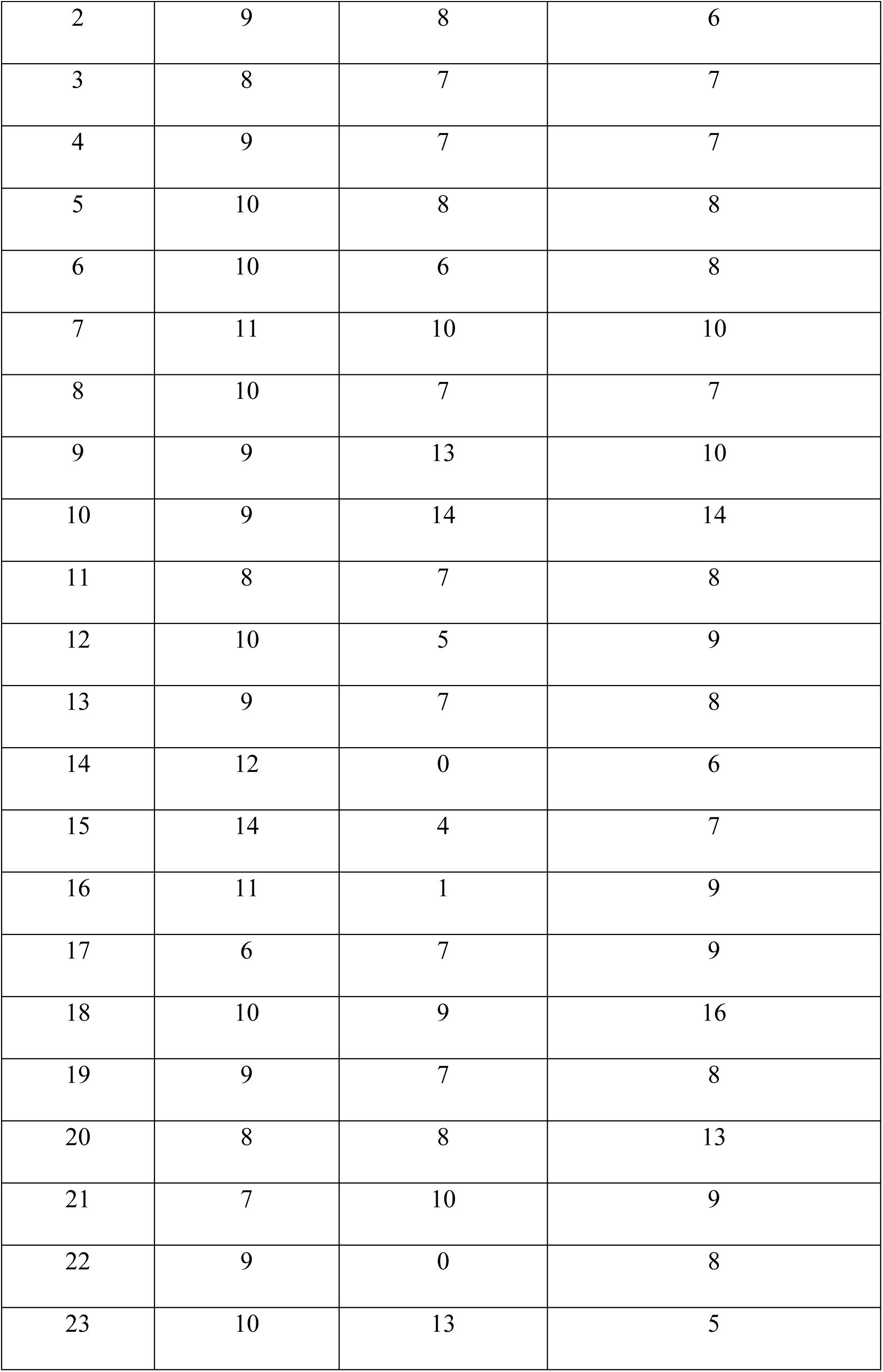
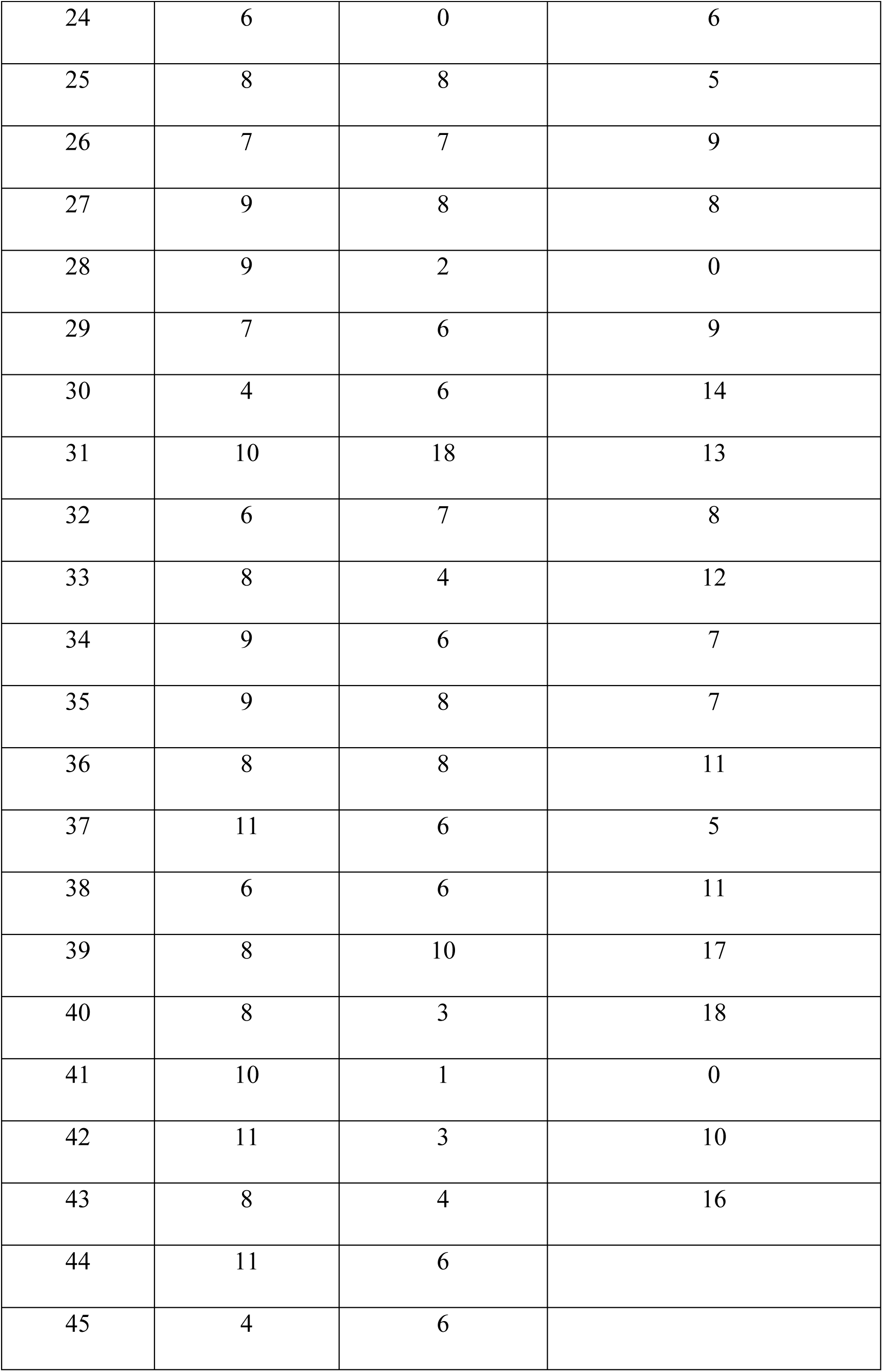
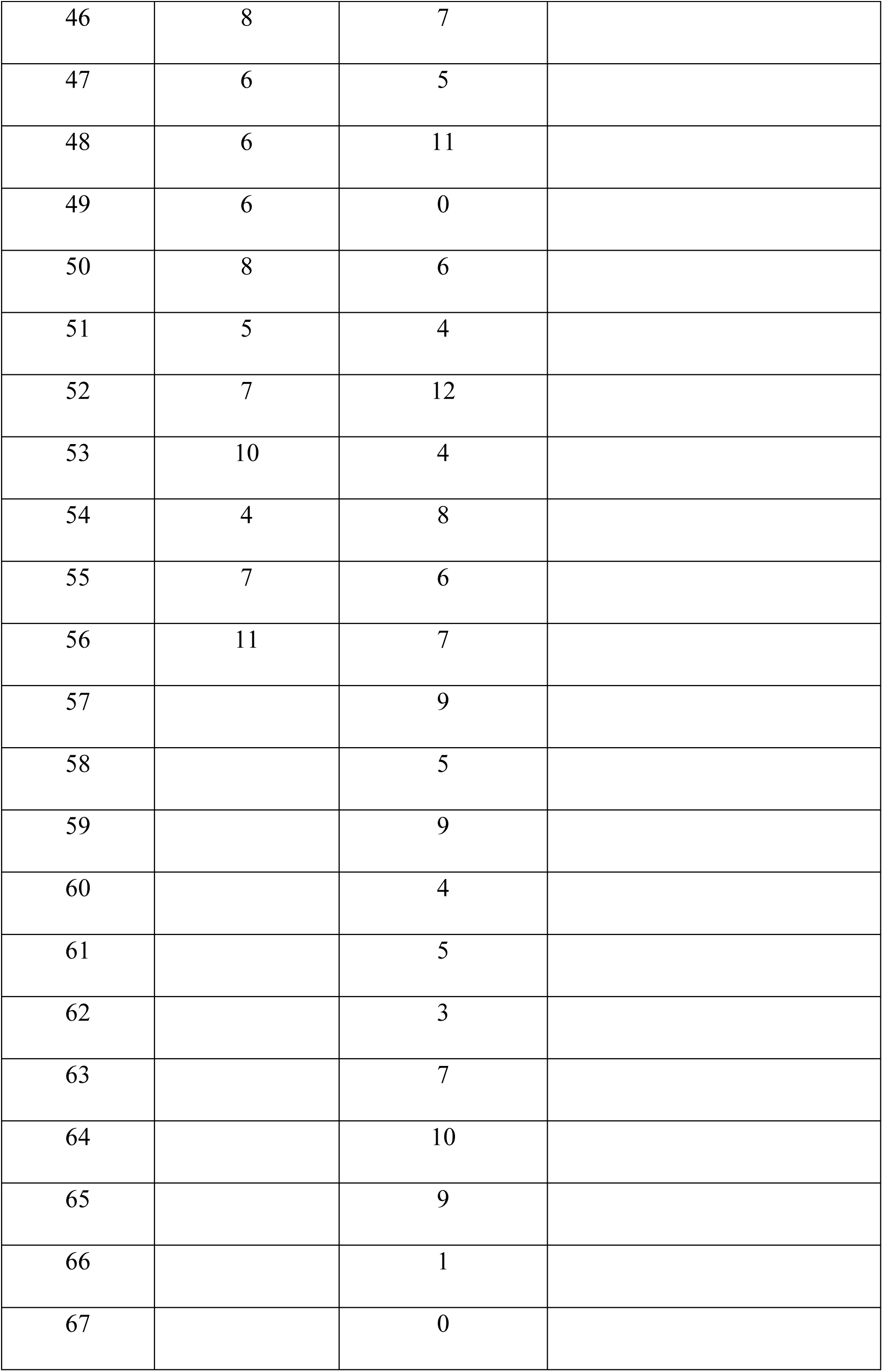
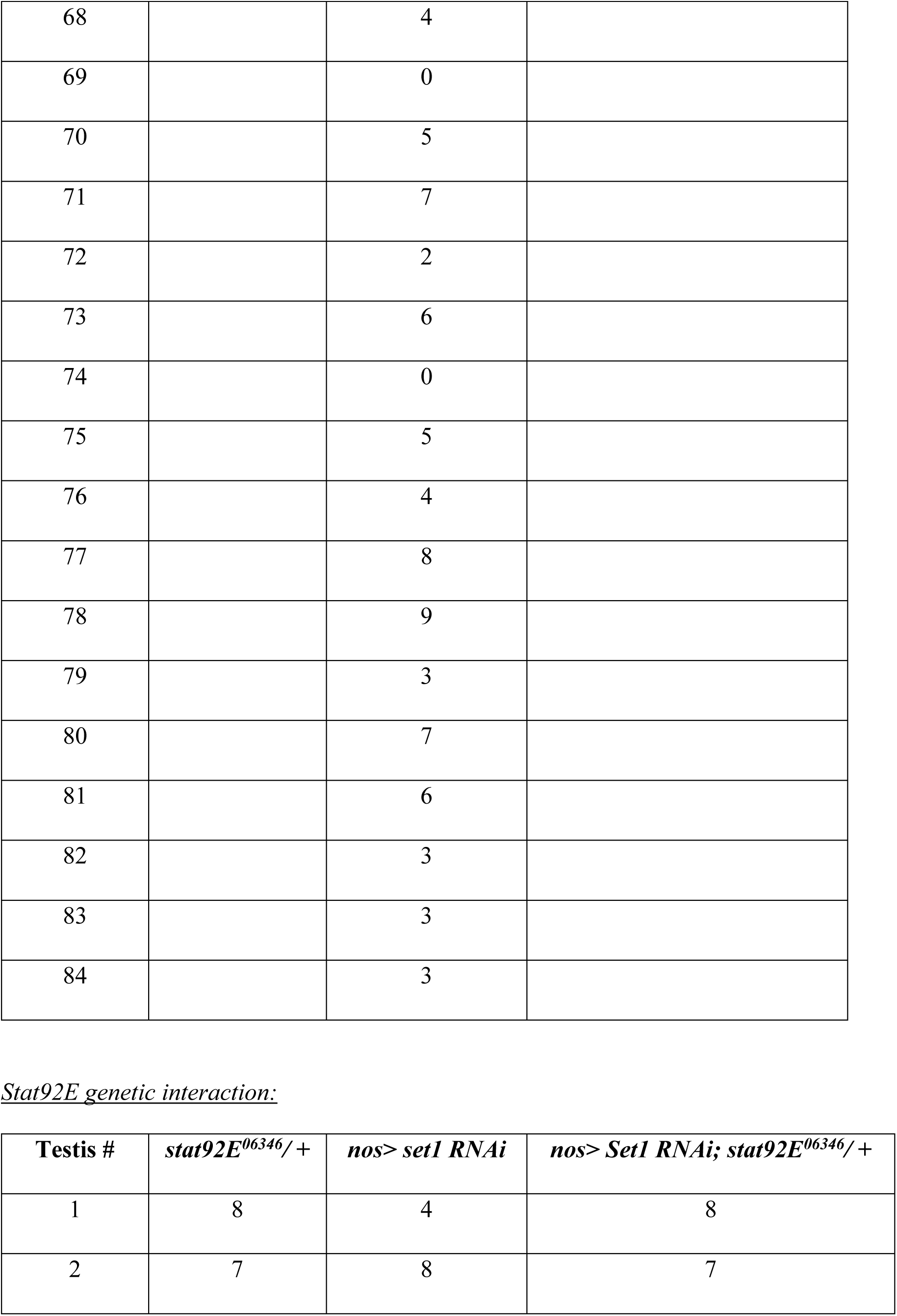
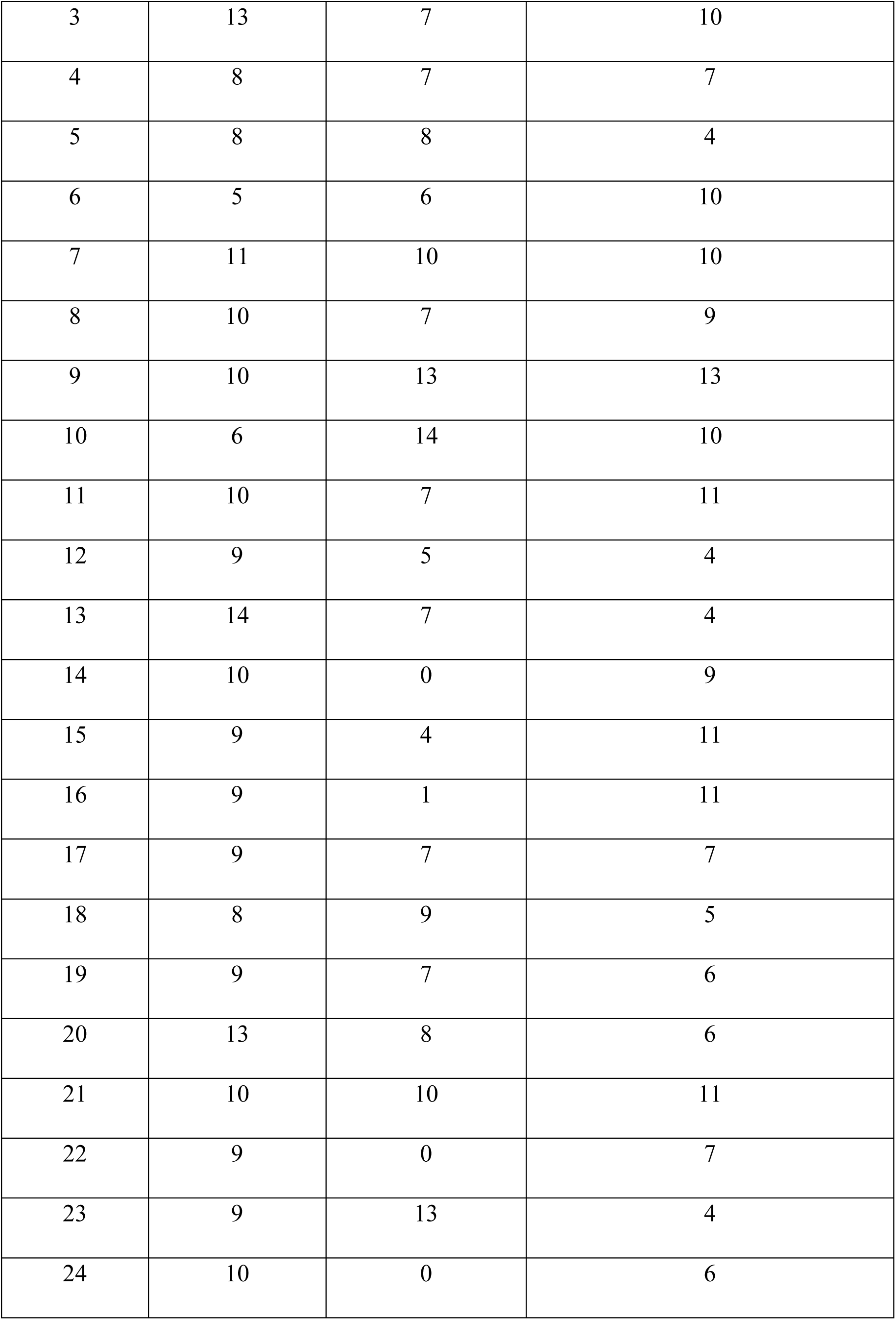
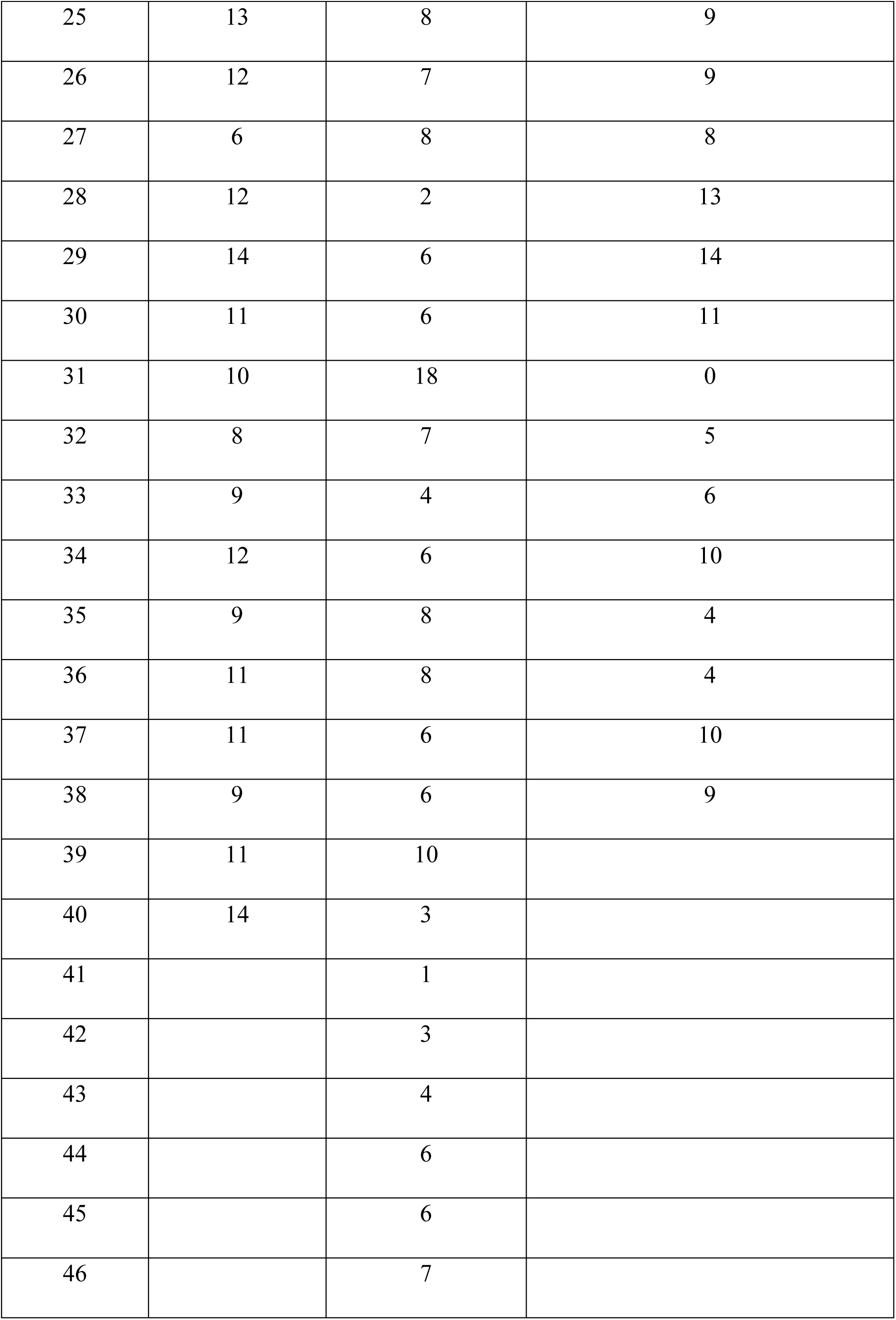
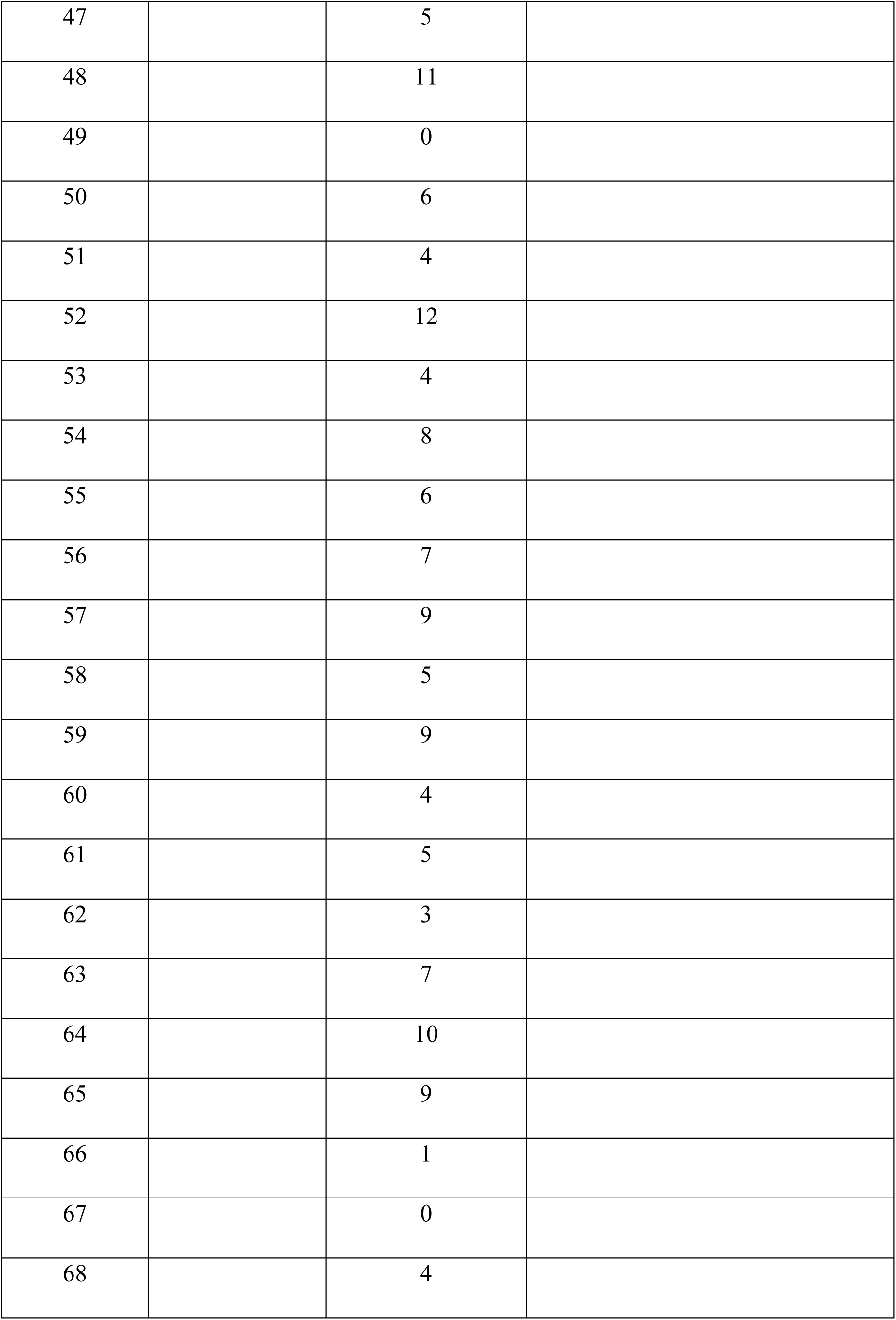
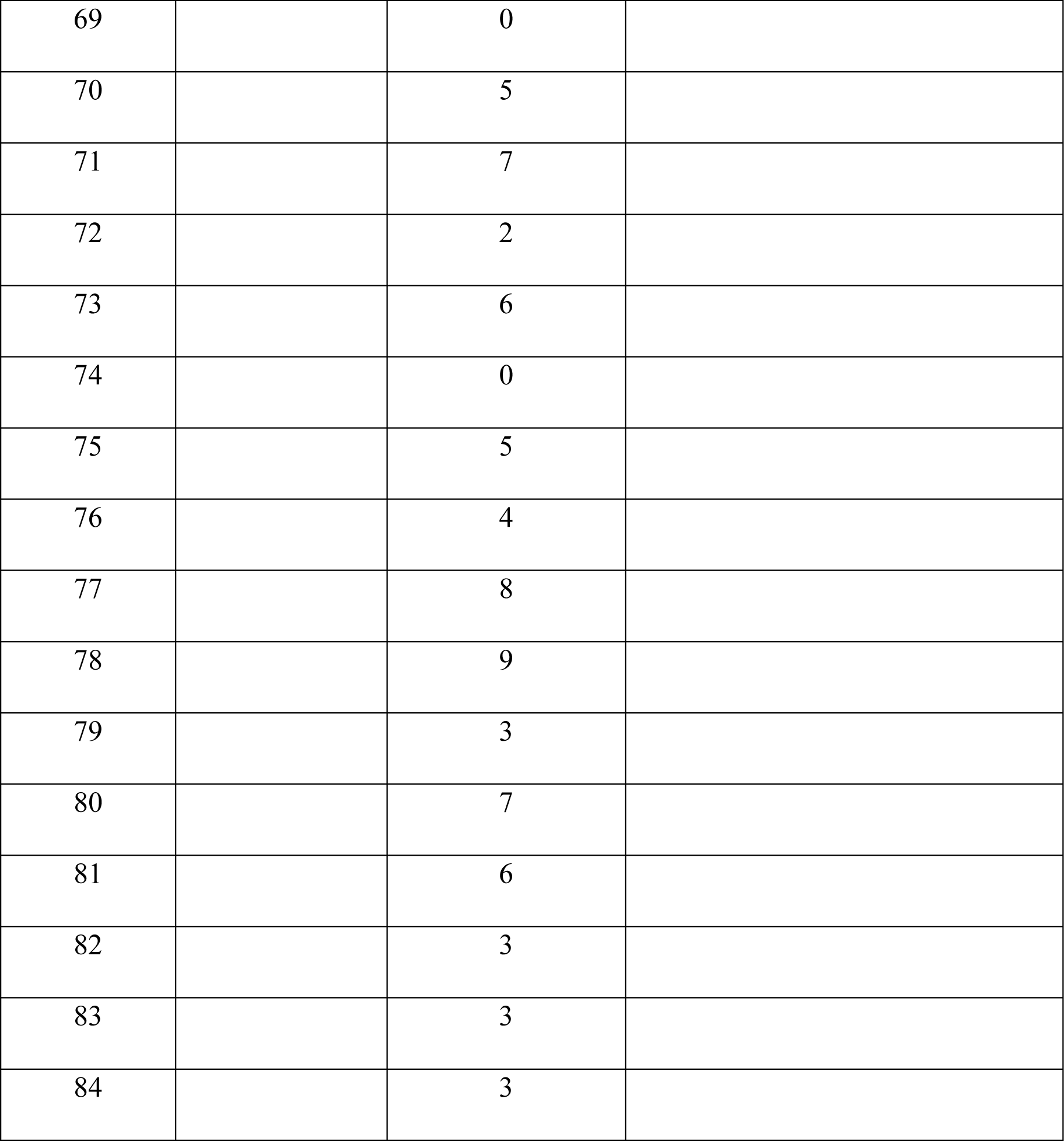
Quantification of Germline Stem Cell number in RNAi knockdown testes.

**Table S2:**
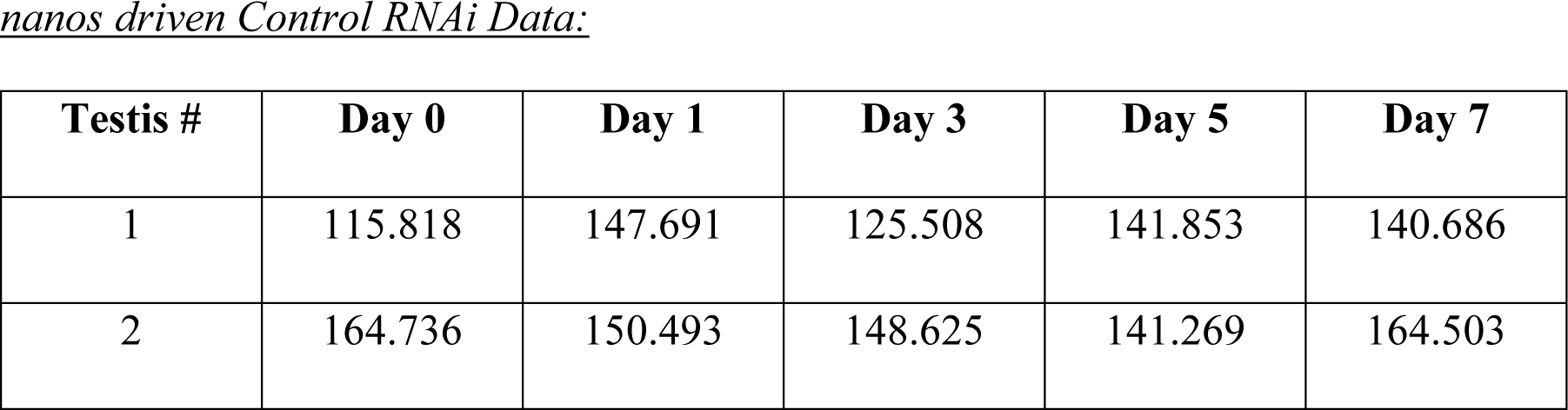
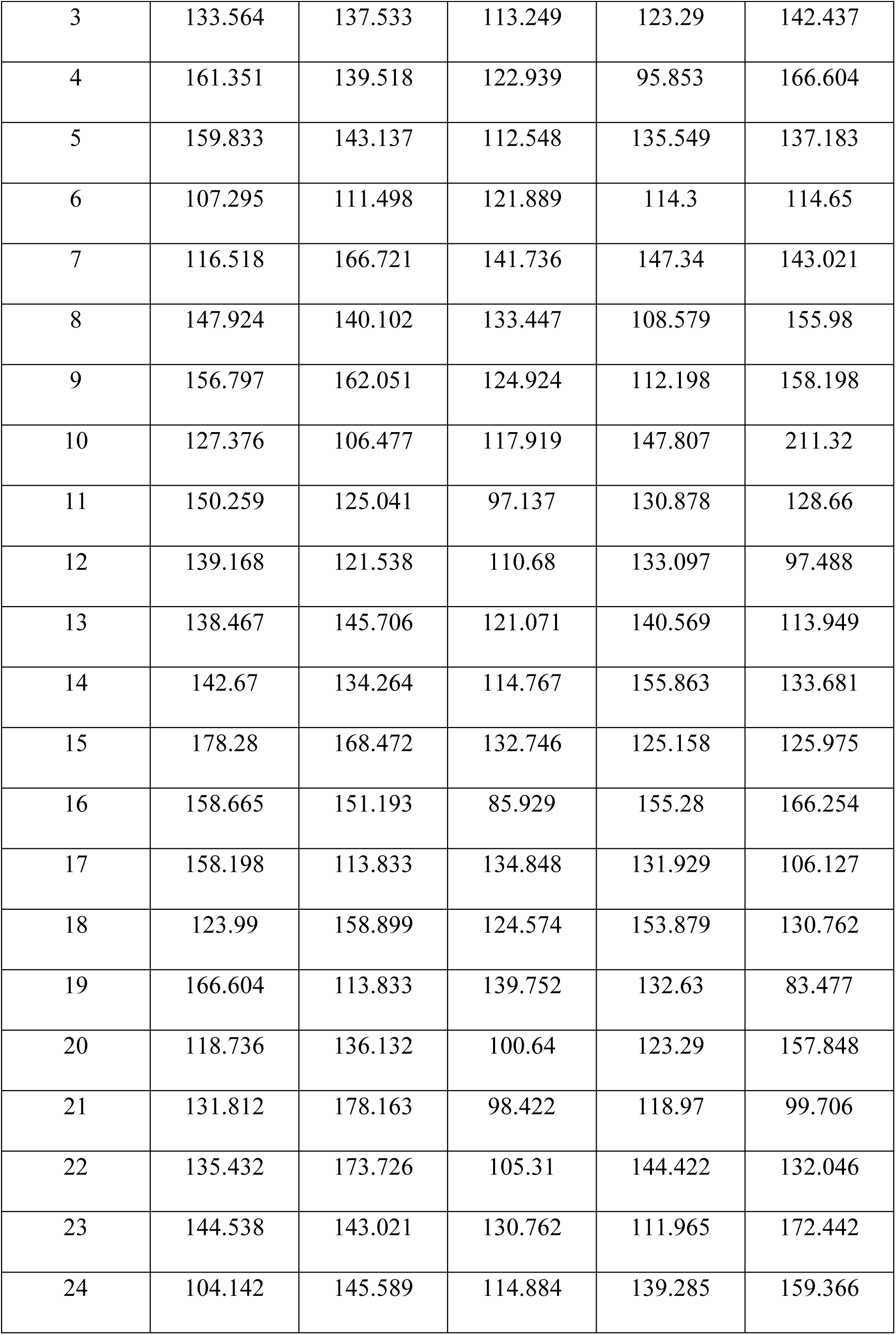
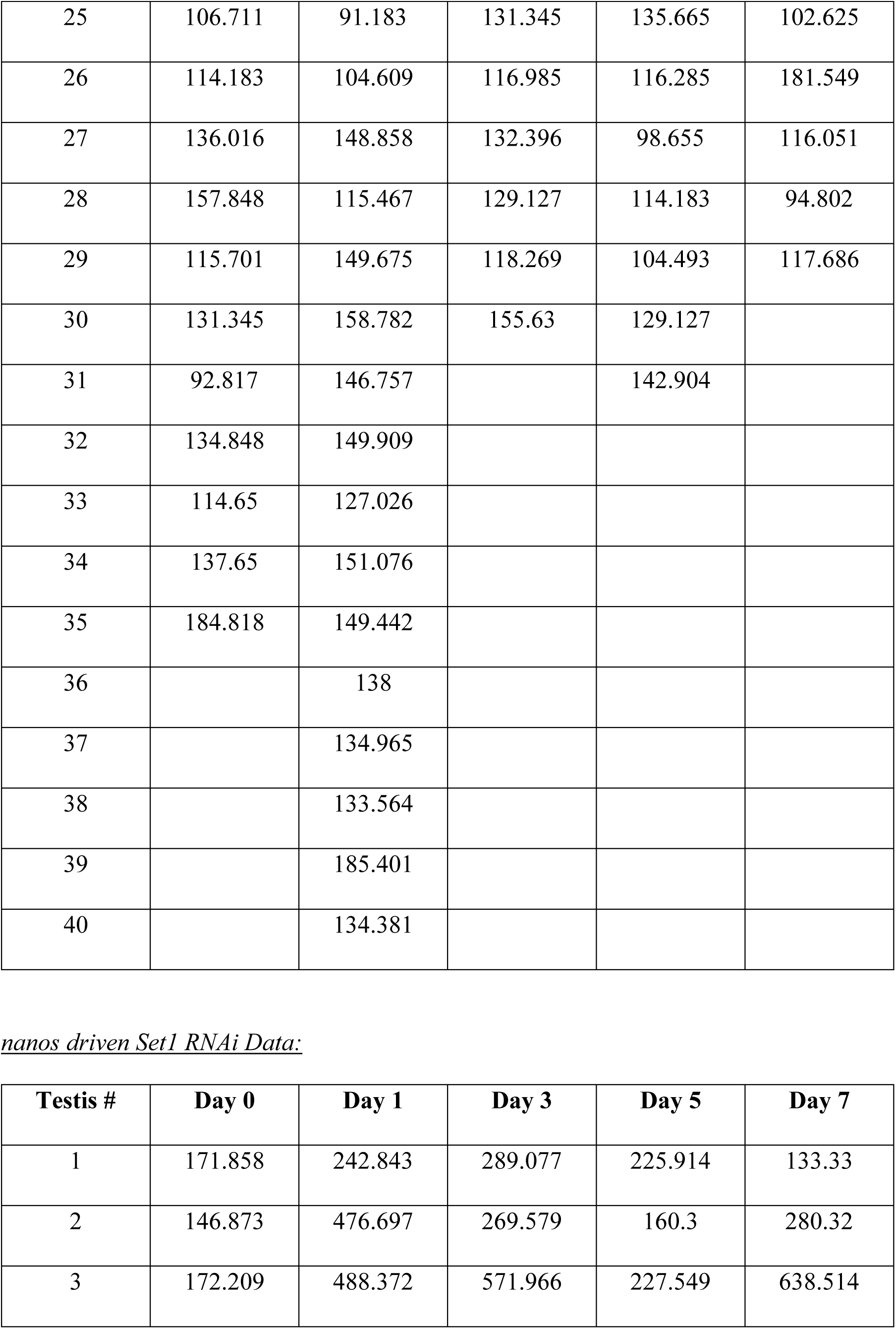
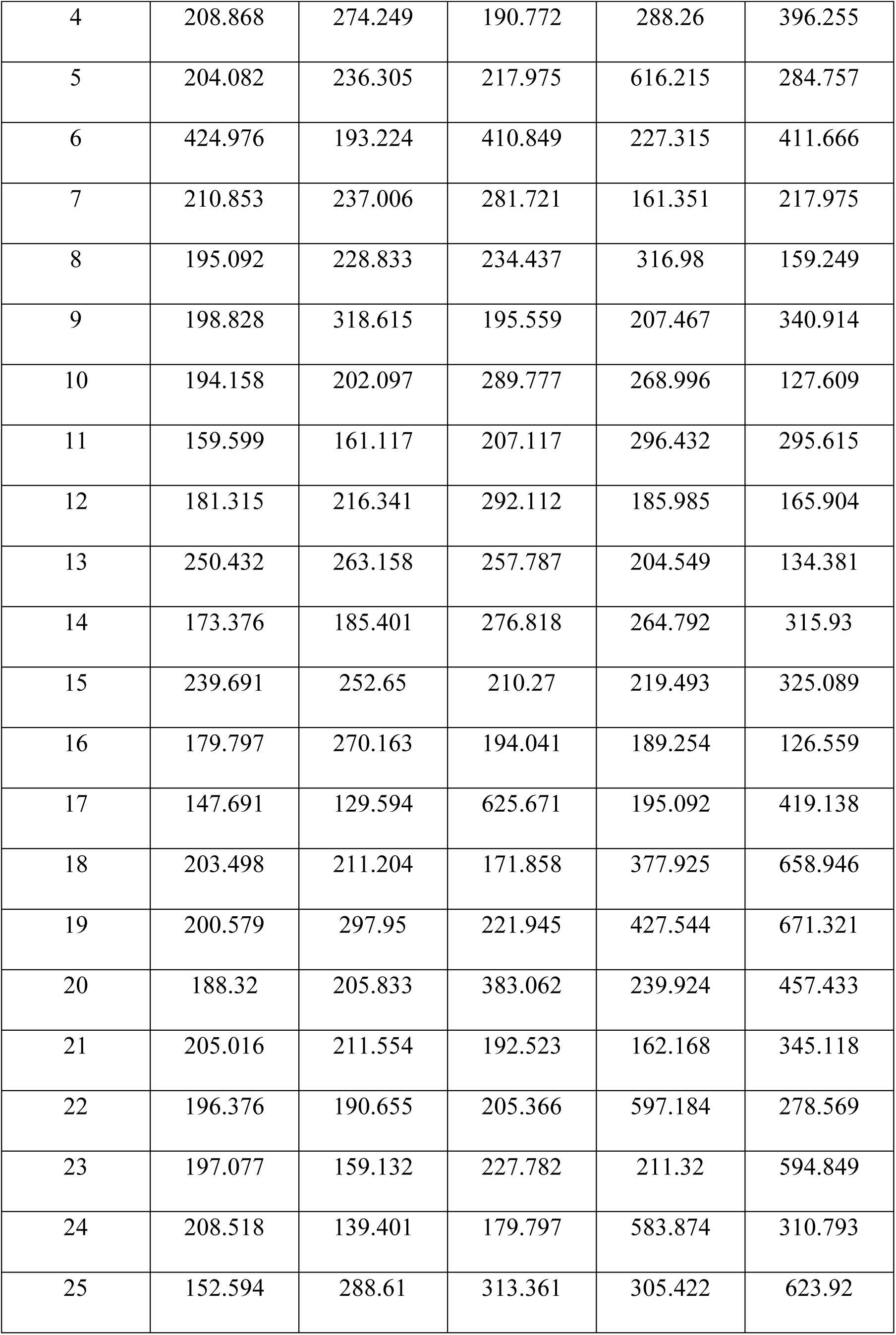
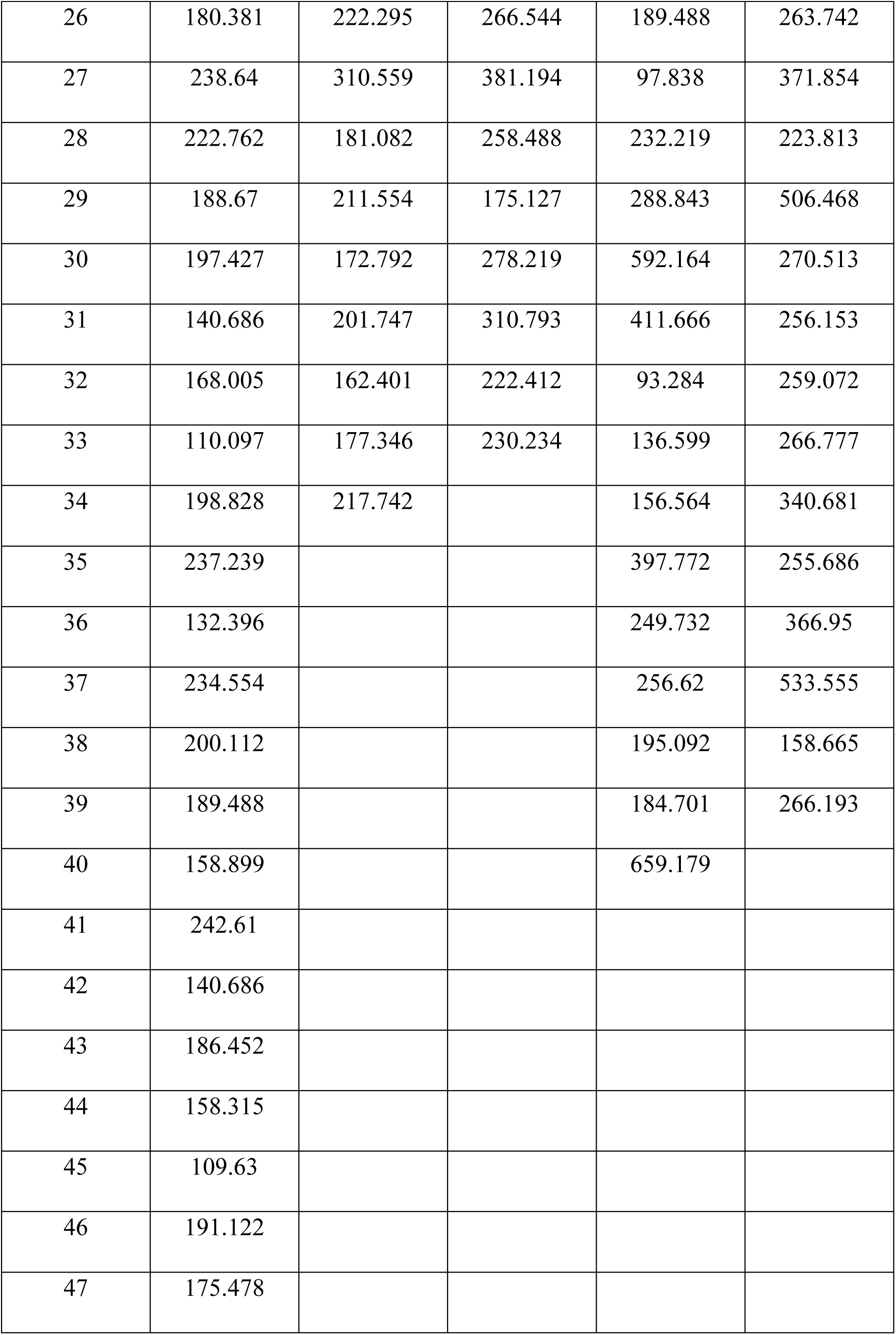
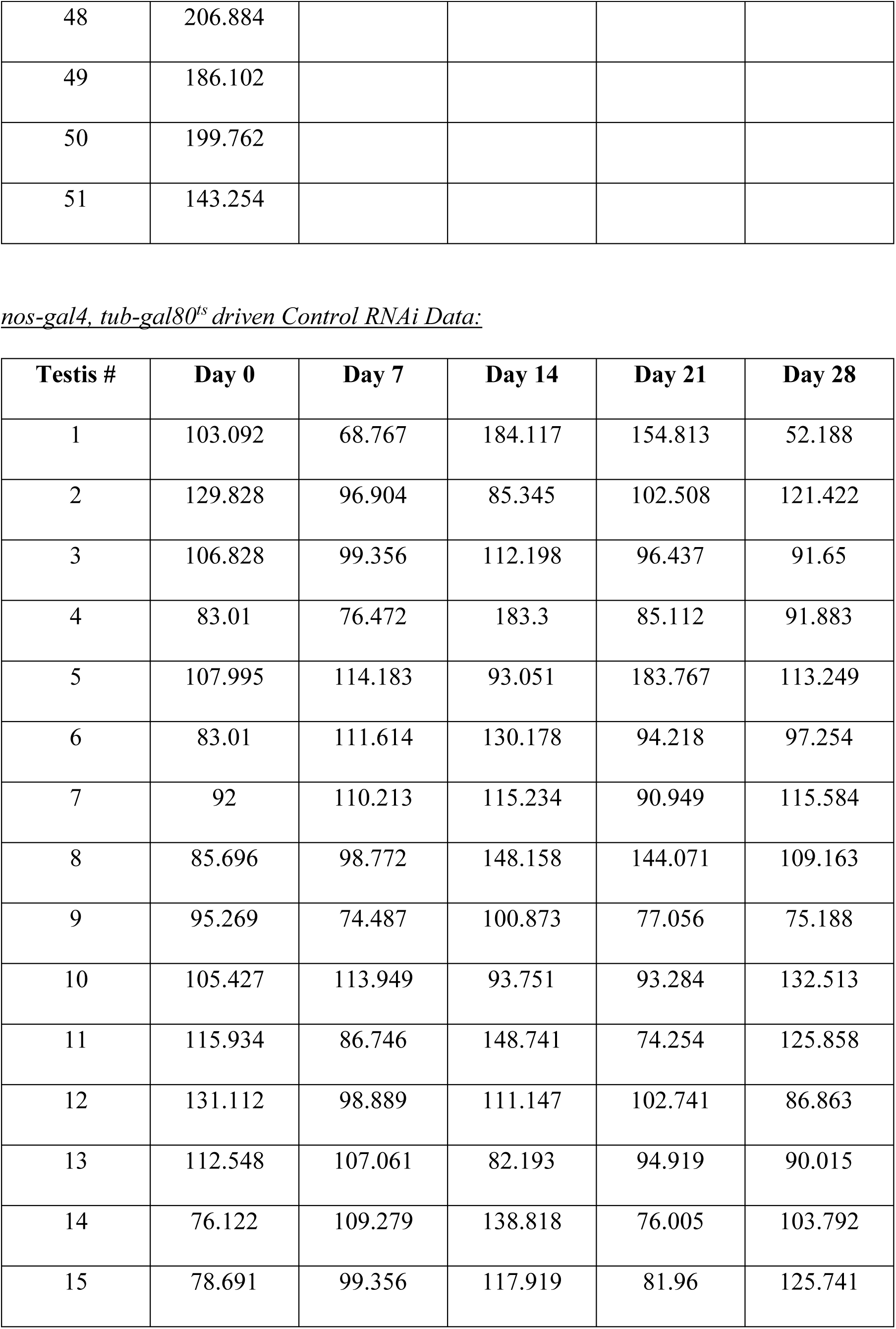
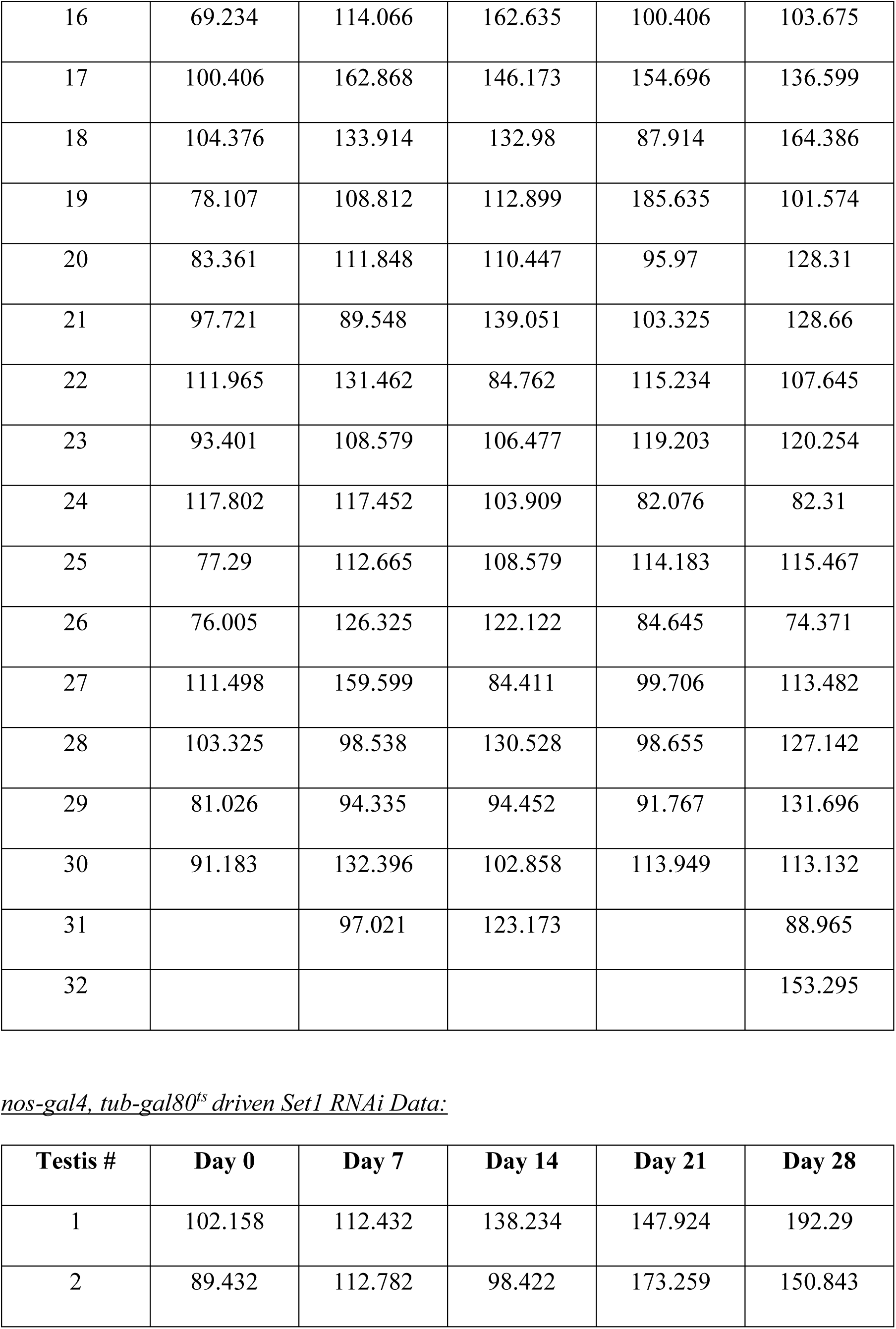
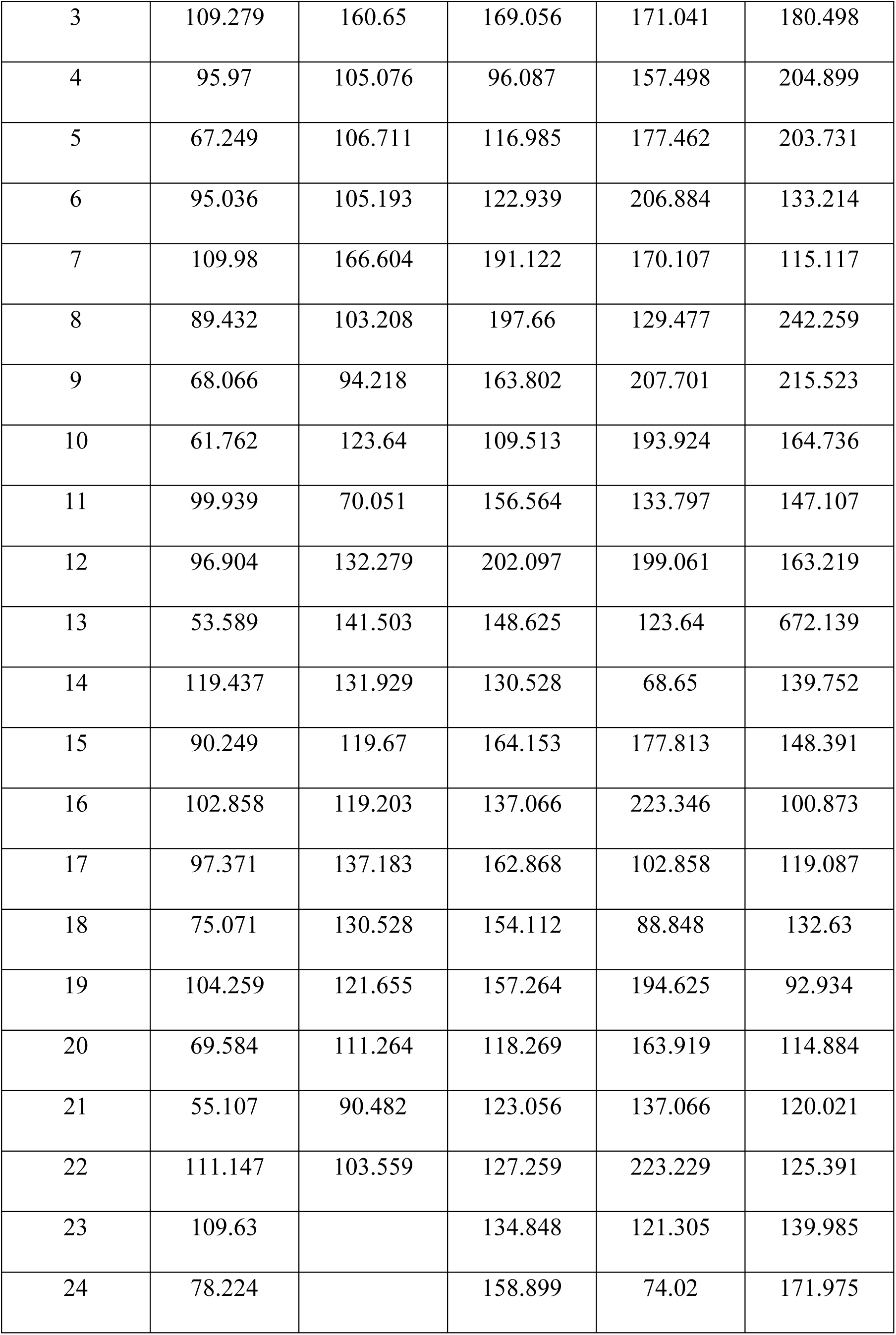
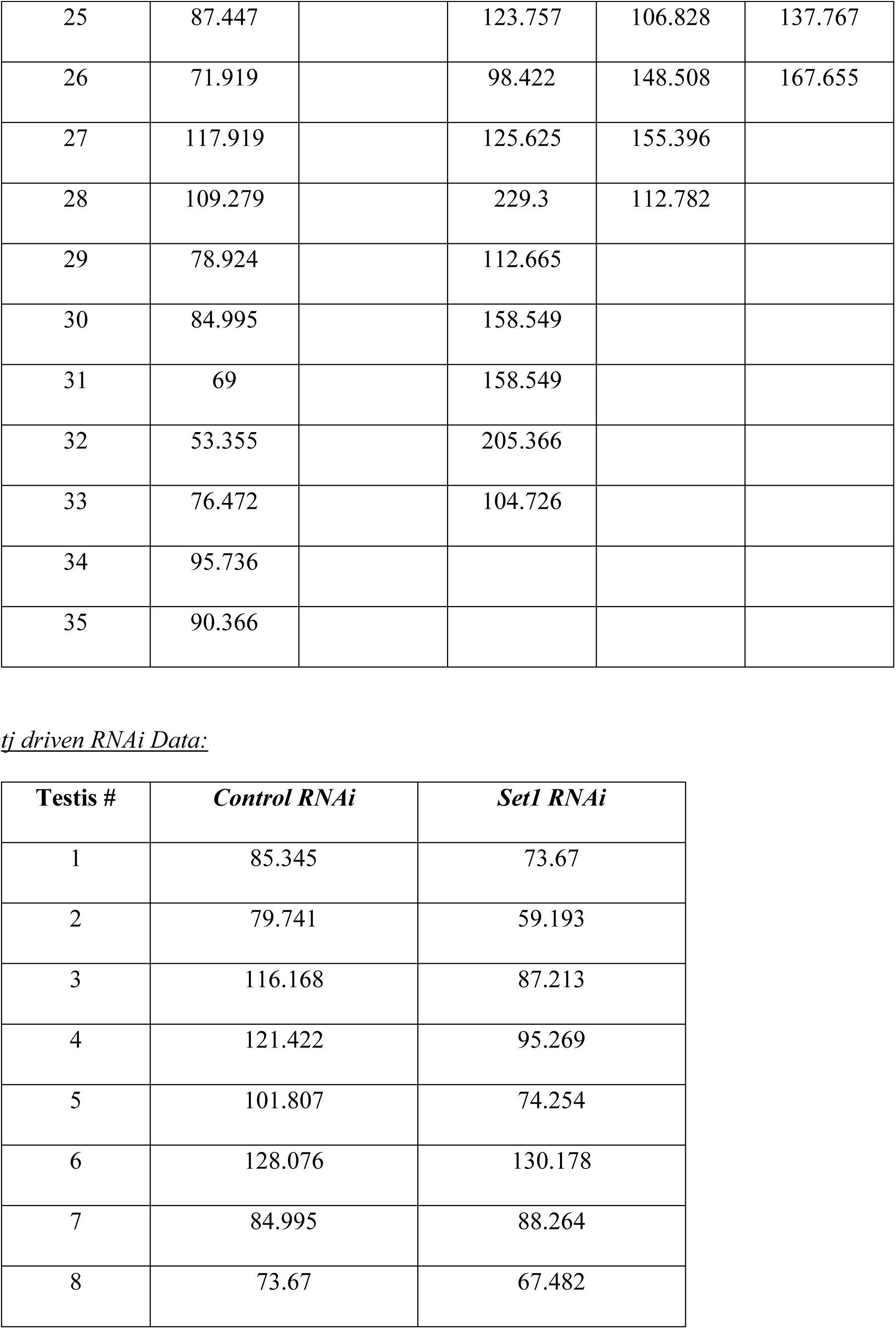
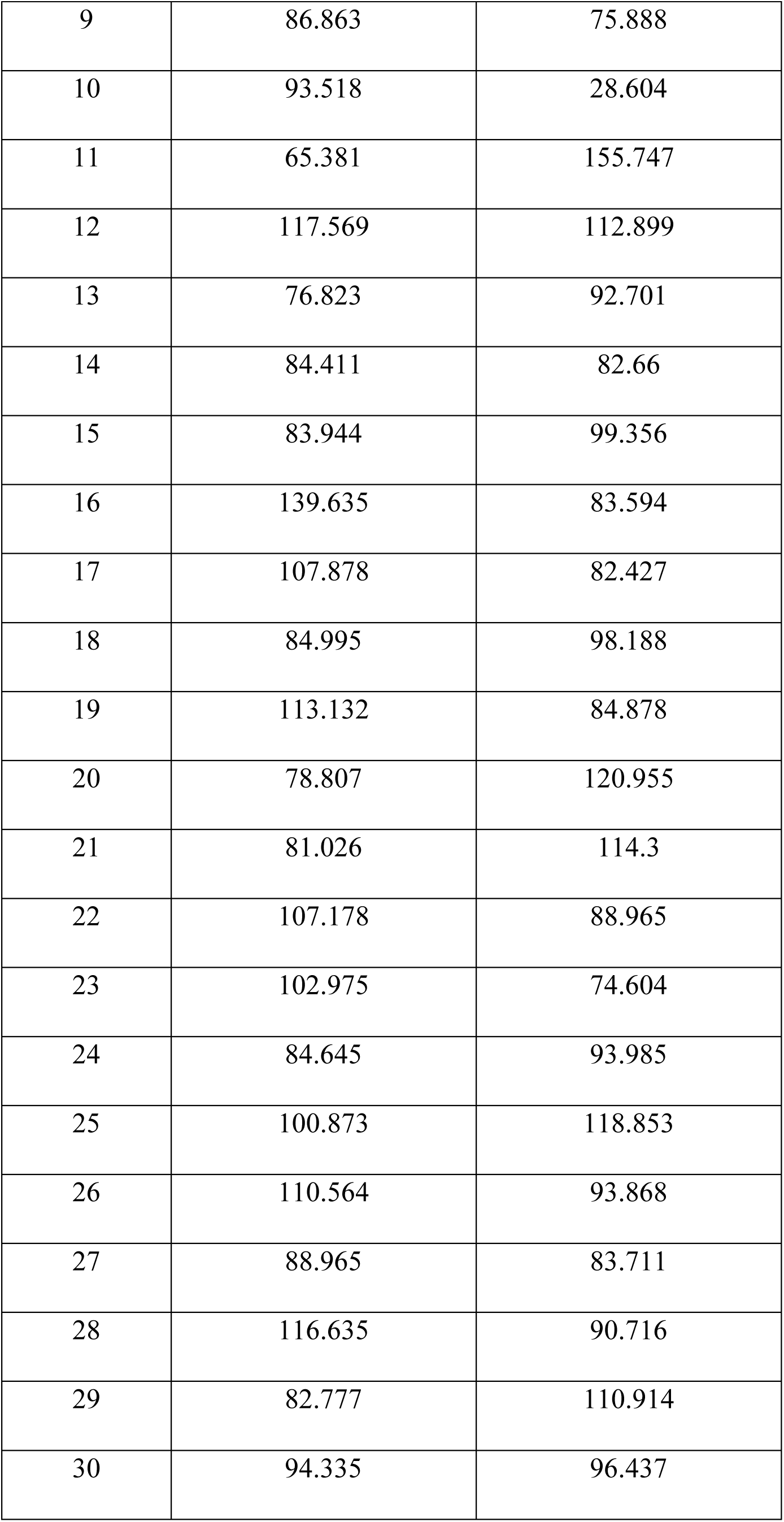
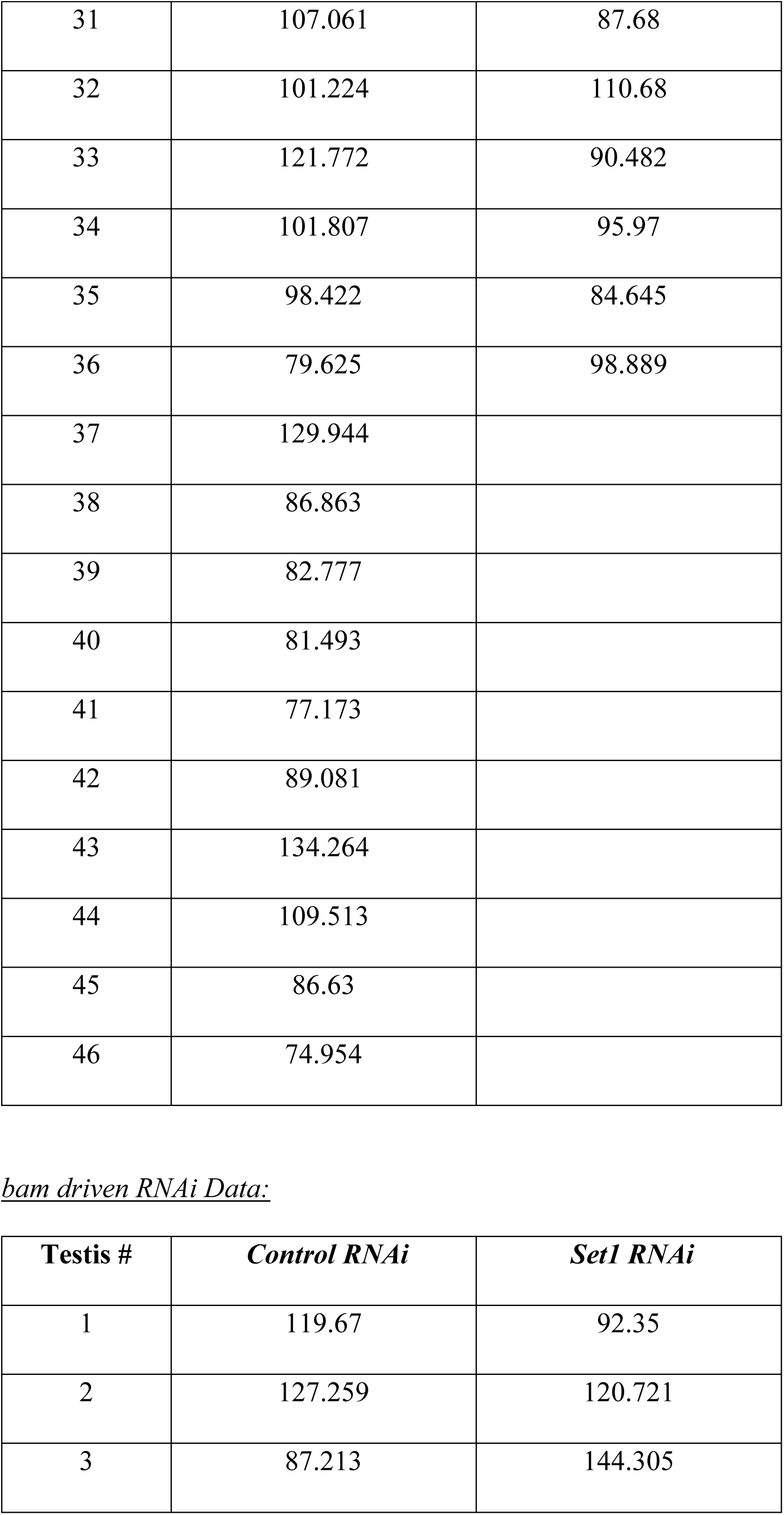
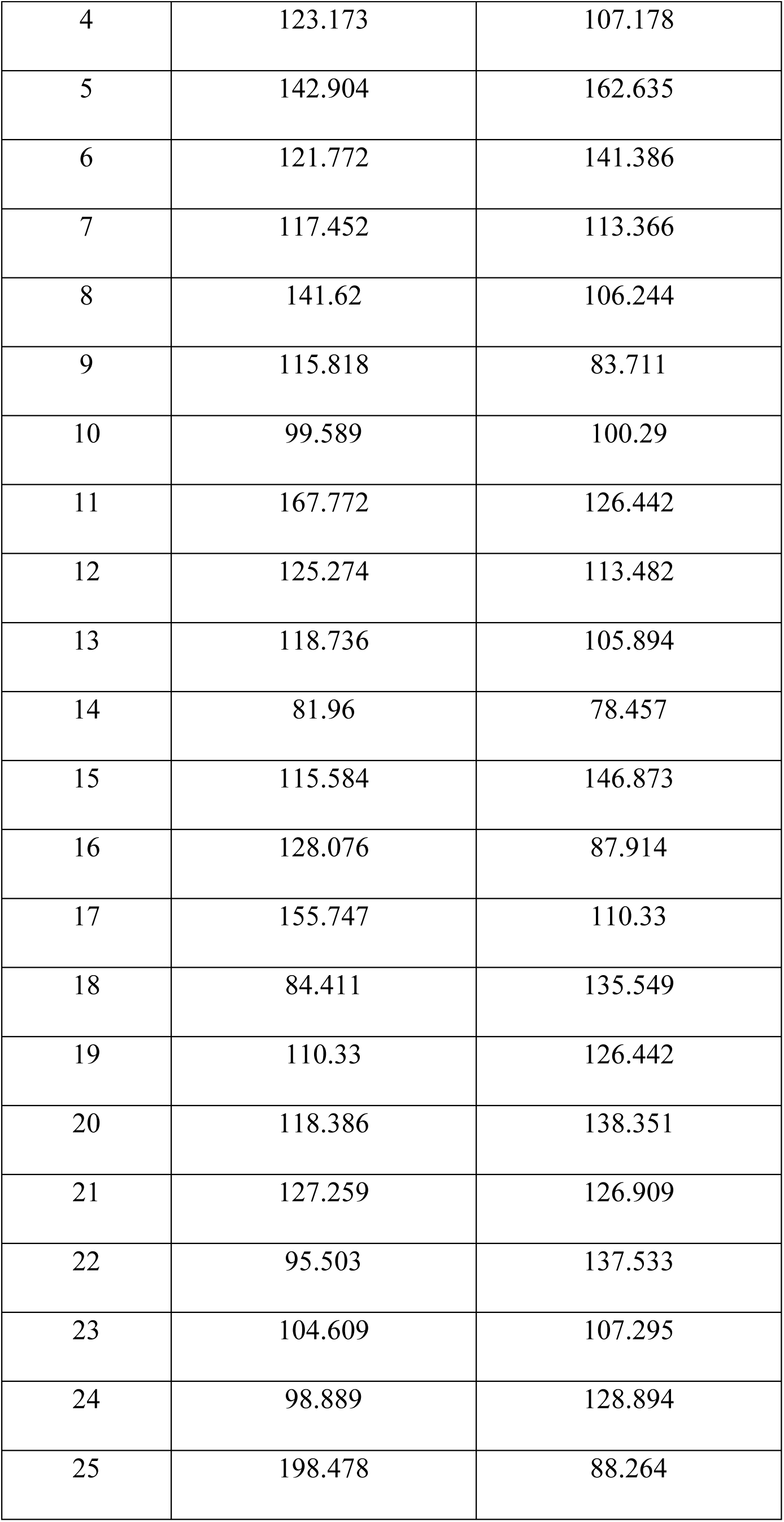
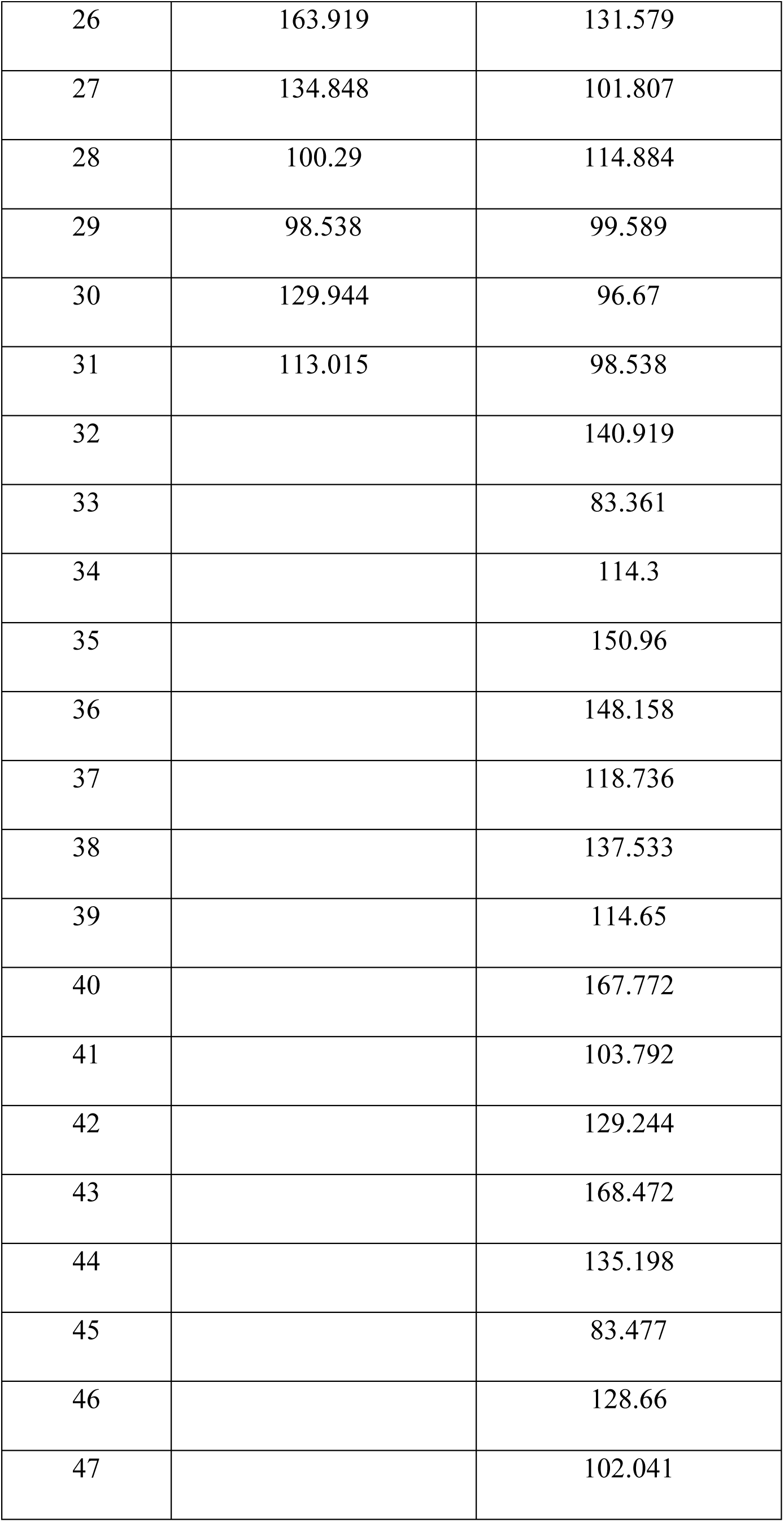
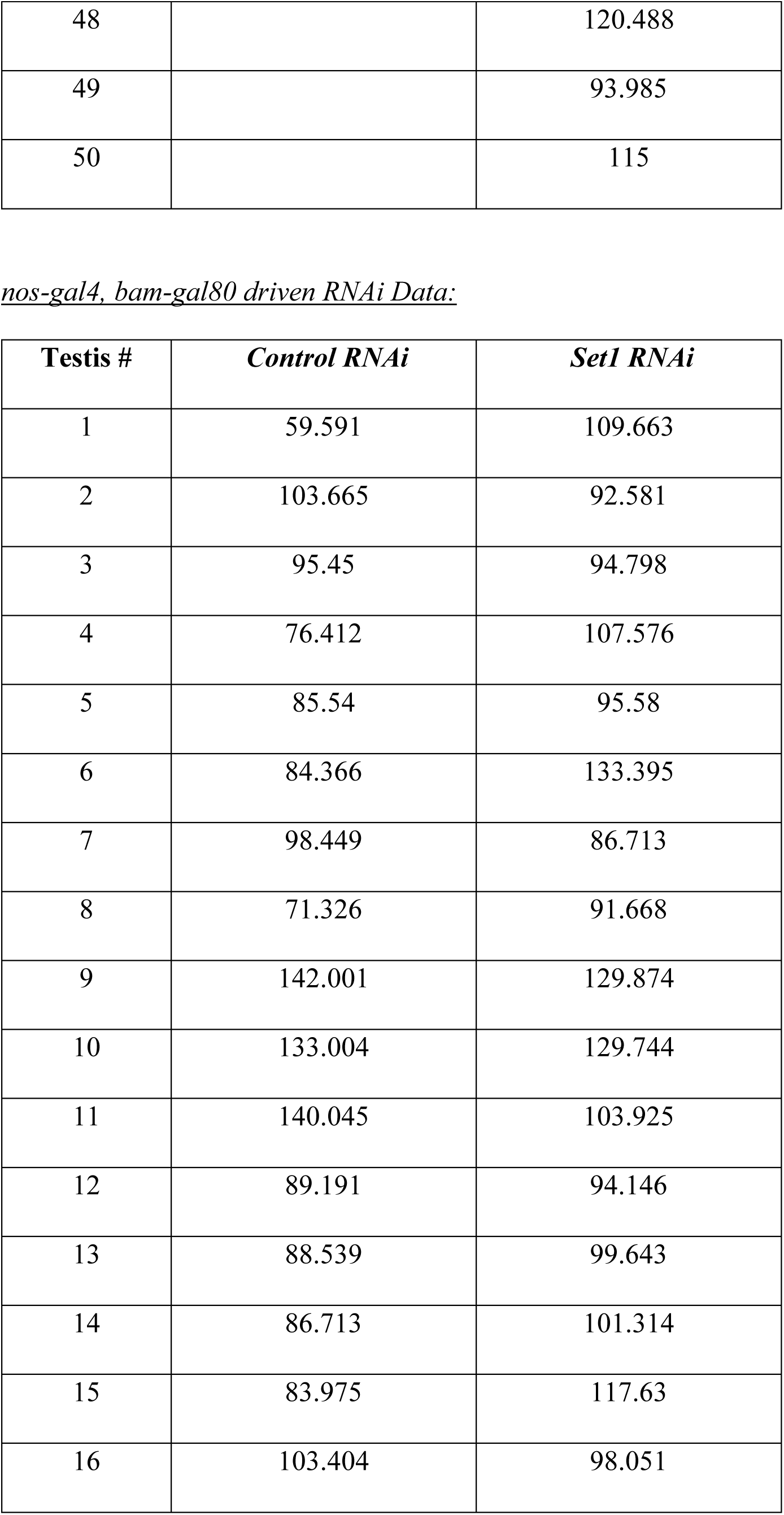
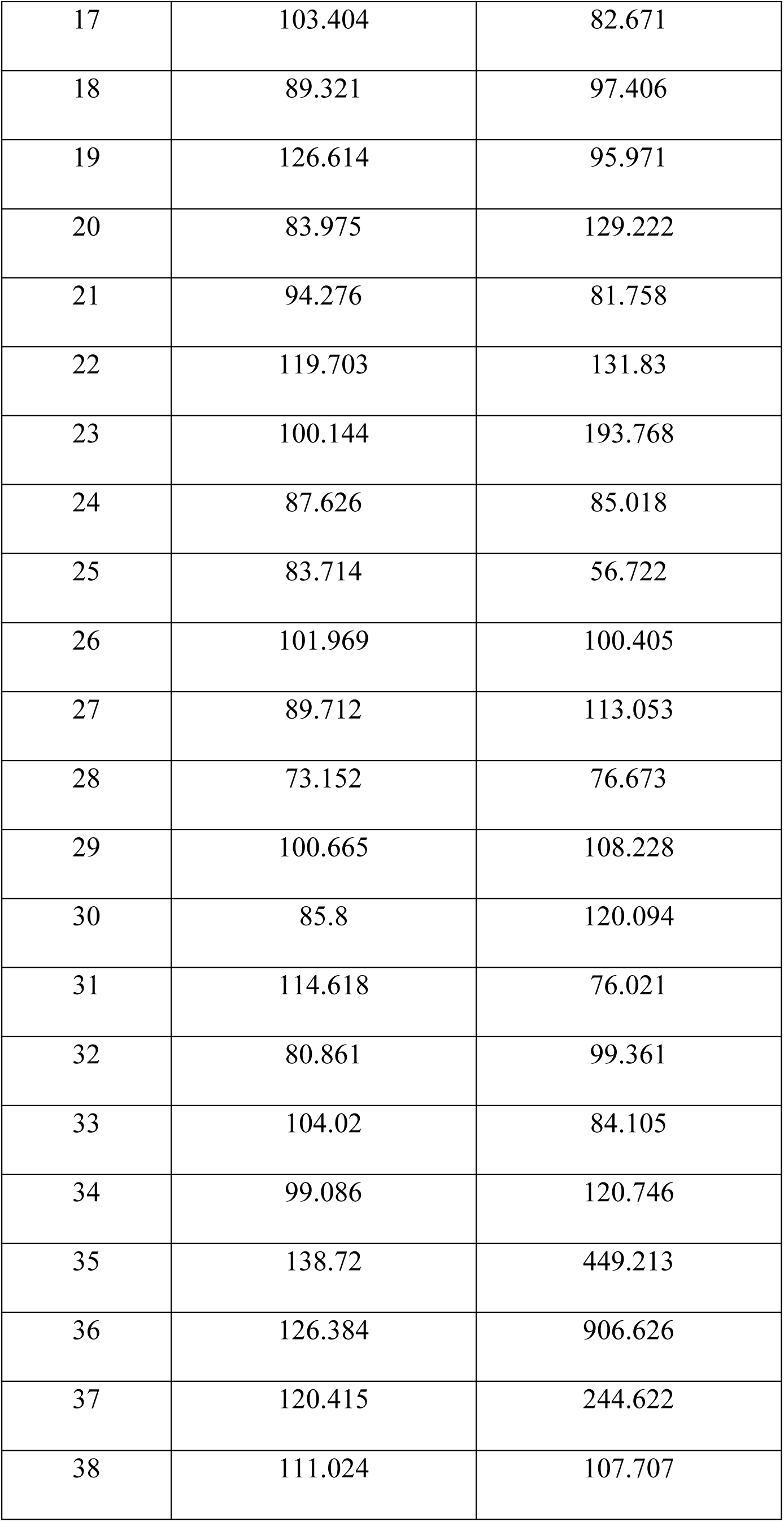
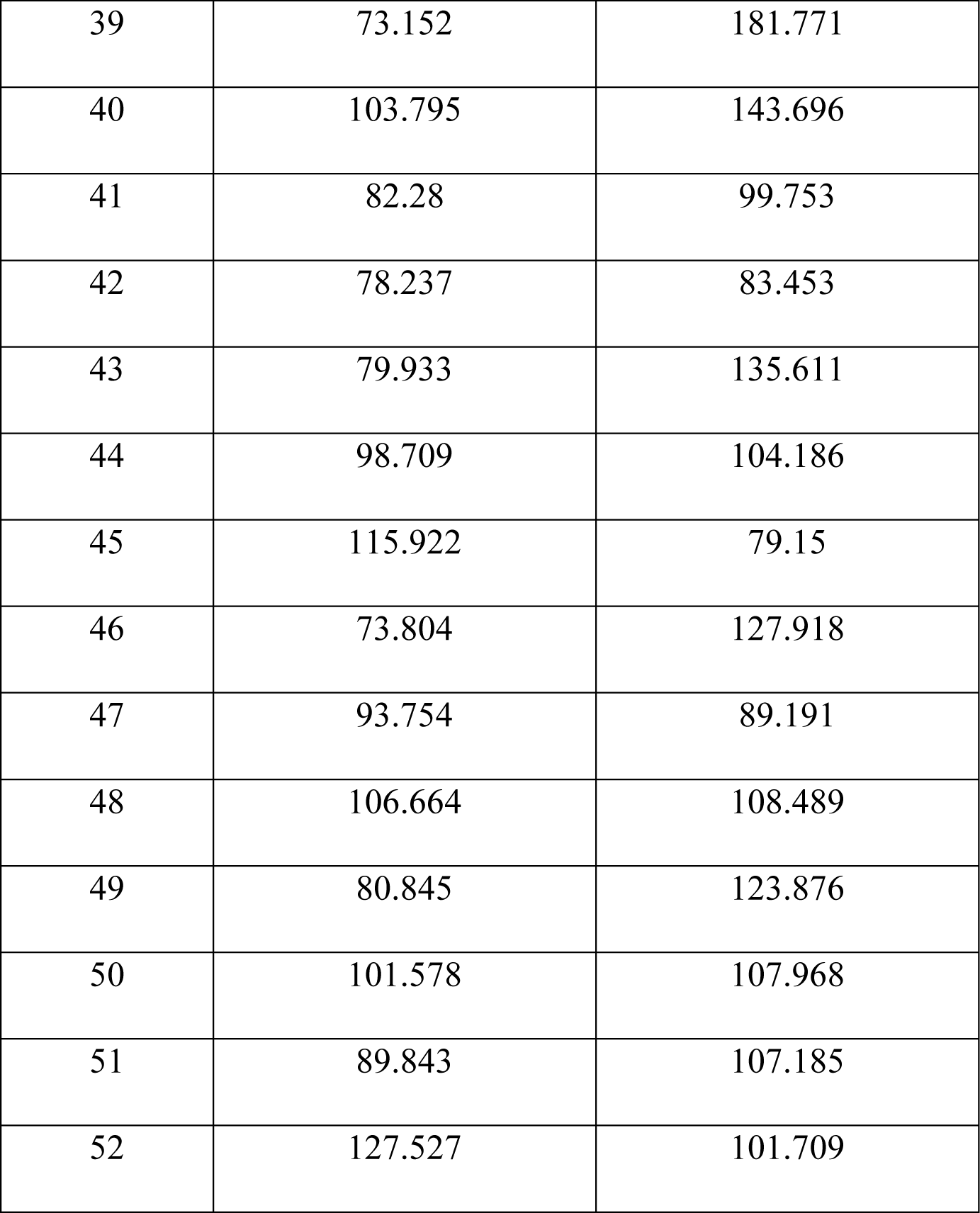
Quantification of Hub area in RNAi knockdown testes.

| Testis # | Day 0 | Day 1 | Day 3 | Day 5 | Day 7 |
| --- | --- | --- | --- | --- | --- |
| 1 | 115.818 | 147.691 | 125.508 | 141.853 | 140.686 |
| 2 | 164.736 | 150.493 | 148.625 | 141.269 | 164.503 |
| 3 | 133.564 | 137.533 | 113.249 | 123.29 | 142.437 |
| 4 | 161.351 | 139.518 | 122.939 | 95.853 | 166.604 |
| 5 | 159.833 | 143.137 | 112.548 | 135.549 | 137.183 |
| 6 | 107.295 | 111.498 | 121.889 | 114.3 | 114.65 |
| 7 | 116.518 | 166.721 | 141.736 | 147.34 | 143.021 |
| 8 | 147.924 | 140.102 | 133.447 | 108.579 | 155.98 |
| 9 | 156.797 | 162.051 | 124.924 | 112.198 | 158.198 |
| 10 | 127.376 | 106.477 | 117.919 | 147.807 | 211.32 |
| 11 | 150.259 | 125.041 | 97.137 | 130.878 | 128.66 |
| 12 | 139.168 | 121.538 | 110.68 | 133.097 | 97.488 |
| 13 | 138.467 | 145.706 | 121.071 | 140.569 | 113.949 |
| 14 | 142.67 | 134.264 | 114.767 | 155.863 | 133.681 |
| 15 | 178.28 | 168.472 | 132.746 | 125.158 | 125.975 |
| 16 | 158.665 | 151.193 | 85.929 | 155.28 | 166.254 |
| 17 | 158.198 | 113.833 | 134.848 | 131.929 | 106.127 |
| 18 | 123.99 | 158.899 | 124.574 | 153.879 | 130.762 |
| 19 | 166.604 | 113.833 | 139.752 | 132.63 | 83.477 |
| 20 | 118.736 | 136.132 | 100.64 | 123.29 | 157.848 |
| 21 | 131.812 | 178.163 | 98.422 | 118.97 | 99.706 |
| 22 | 135.432 | 173.726 | 105.31 | 144.422 | 132.046 |
| 23 | 144.538 | 143.021 | 130.762 | 111.965 | 172.442 |
| 24 | 104.142 | 145.589 | 114.884 | 139.285 | 159.366 |
| 25 | 106.711 | 91.183 | 131.345 | 135.665 | 102.625 |
| 26 | 114.183 | 104.609 | 116.985 | 116.285 | 181.549 |
| 27 | 136.016 | 148.858 | 132.396 | 98.655 | 116.051 |
| 28 | 157.848 | 115.467 | 129.127 | 114.183 | 94.802 |
| 29 | 115.701 | 149.675 | 118.269 | 104.493 | 117.686 |
| 30 | 131.345 | 158.782 | 155.63 | 129.127 |  |
| 31 | 92.817 | 146.757 |  | 142.904 |  |
| 32 | 134.848 | 149.909 |  |  |  |
| 33 | 114.65 | 127.026 |  |  |  |
| 34 | 137.65 | 151.076 |  |  |  |
| 35 | 184.818 | 149.442 |  |  |  |
| 36 |  | 138 |  |  |  |
| 37 |  | 134.965 |  |  |  |
| 38 |  | 133.564 |  |  |  |
| 39 |  | 185.401 |  |  |  |
| 40 |  | 134.381 |  |  |  |

*nanos driven Set1 RNAi Data:*
| **Testis #** | **Day 0** | **Day 1** | **Day 3** | **Day 5** | **Day 7** |
| --- | --- | --- | --- | --- | --- |
| 1 | 171.858 | 242.843 | 289.077 | 225.914 | 133.33 |
| 2 | 146.873 | 476.697 | 269.579 | 160.3 | 280.32 |
| 3 | 172.209 | 488.372 | 571.966 | 227.549 | 638.514 |
| 4 | 208.868 | 274.249 | 190.772 | 288.26 | 396.255 |
| 5 | 204.082 | 236.305 | 217.975 | 616.215 | 284.757 |
| 6 | 424.976 | 193.224 | 410.849 | 227.315 | 411.666 |
| 7 | 210.853 | 237.006 | 281.721 | 161.351 | 217.975 |
| 8 | 195.092 | 228.833 | 234.437 | 316.98 | 159.249 |
| 9 | 198.828 | 318.615 | 195.559 | 207.467 | 340.914 |
| 10 | 194.158 | 202.097 | 289.777 | 268.996 | 127.609 |
| 11 | 159.599 | 161.117 | 207.117 | 296.432 | 295.615 |
| 12 | 181.315 | 216.341 | 292.112 | 185.985 | 165.904 |
| 13 | 250.432 | 263.158 | 257.787 | 204.549 | 134.381 |
| 14 | 173.376 | 185.401 | 276.818 | 264.792 | 315.93 |
| 15 | 239.691 | 252.65 | 210.27 | 219.493 | 325.089 |
| 16 | 179.797 | 270.163 | 194.041 | 189.254 | 126.559 |
| 17 | 147.691 | 129.594 | 625.671 | 195.092 | 419.138 |
| 18 | 203.498 | 211.204 | 171.858 | 377.925 | 658.946 |
| 19 | 200.579 | 297.95 | 221.945 | 427.544 | 671.321 |
| 20 | 188.32 | 205.833 | 383.062 | 239.924 | 457.433 |
| 21 | 205.016 | 211.554 | 192.523 | 162.168 | 345.118 |
| 22 | 196.376 | 190.655 | 205.366 | 597.184 | 278.569 |
| 23 | 197.077 | 159.132 | 227.782 | 211.32 | 594.849 |
| 24 | 208.518 | 139.401 | 179.797 | 583.874 | 310.793 |
| 25 | 152.594 | 288.61 | 313.361 | 305.422 | 623.92 |
| 26 | 180.381 | 222.295 | 266.544 | 189.488 | 263.742 |
| 27 | 238.64 | 310.559 | 381.194 | 97.838 | 371.854 |
| 28 | 222.762 | 181.082 | 258.488 | 232.219 | 223.813 |
| 29 | 188.67 | 211.554 | 175.127 | 288.843 | 506.468 |
| 30 | 197.427 | 172.792 | 278.219 | 592.164 | 270.513 |
| 31 | 140.686 | 201.747 | 310.793 | 411.666 | 256.153 |
| 32 | 168.005 | 162.401 | 222.412 | 93.284 | 259.072 |
| 33 | 110.097 | 177.346 | 230.234 | 136.599 | 266.777 |
| 34 | 198.828 | 217.742 |  | 156.564 | 340.681 |
| 35 | 237.239 |  |  | 397.772 | 255.686 |
| 36 | 132.396 |  |  | 249.732 | 366.95 |
| 37 | 234.554 |  |  | 256.62 | 533.555 |
| 38 | 200.112 |  |  | 195.092 | 158.665 |
| 39 | 189.488 |  |  | 184.701 | 266.193 |
| 40 | 158.899 |  |  | 659.179 |  |
| 41 | 242.61 |  |  |  |  |
| 42 | 140.686 |  |  |  |  |
| 43 | 186.452 |  |  |  |  |
| 44 | 158.315 |  |  |  |  |
| 45 | 109.63 |  |  |  |  |
| 46 | 191.122 |  |  |  |  |
| 47 | 175.478 |  |  |  |  |
| 48 | 206.884 |  |  |  |  |
| 49 | 186.102 |  |  |  |  |
| 50 | 199.762 |  |  |  |  |
| 51 | 143.254 |  |  |  |  |

*nos-gal4, tub-gal80<sup>ts</sup> driven Control RNAi Data:*
| **Testis #** | **Day 0** | **Day 7** | **Day 14** | **Day 21** | **Day 28** |
| --- | --- | --- | --- | --- | --- |
| 1 | 103.092 | 68.767 | 184.117 | 154.813 | 52.188 |
| 2 | 129.828 | 96.904 | 85.345 | 102.508 | 121.422 |
| 3 | 106.828 | 99.356 | 112.198 | 96.437 | 91.65 |
| 4 | 83.01 | 76.472 | 183.3 | 85.112 | 91.883 |
| 5 | 107.995 | 114.183 | 93.051 | 183.767 | 113.249 |
| 6 | 83.01 | 111.614 | 130.178 | 94.218 | 97.254 |
| 7 | 92 | 110.213 | 115.234 | 90.949 | 115.584 |
| 8 | 85.696 | 98.772 | 148.158 | 144.071 | 109.163 |
| 9 | 95.269 | 74.487 | 100.873 | 77.056 | 75.188 |
| 10 | 105.427 | 113.949 | 93.751 | 93.284 | 132.513 |
| 11 | 115.934 | 86.746 | 148.741 | 74.254 | 125.858 |
| 12 | 131.112 | 98.889 | 111.147 | 102.741 | 86.863 |
| 13 | 112.548 | 107.061 | 82.193 | 94.919 | 90.015 |
| 14 | 76.122 | 109.279 | 138.818 | 76.005 | 103.792 |
| 15 | 78.691 | 99.356 | 117.919 | 81.96 | 125.741 |
| 16 | 69.234 | 114.066 | 162.635 | 100.406 | 103.675 |
| 17 | 100.406 | 162.868 | 146.173 | 154.696 | 136.599 |
| 18 | 104.376 | 133.914 | 132.98 | 87.914 | 164.386 |
| 19 | 78.107 | 108.812 | 112.899 | 185.635 | 101.574 |
| 20 | 83.361 | 111.848 | 110.447 | 95.97 | 128.31 |
| 21 | 97.721 | 89.548 | 139.051 | 103.325 | 128.66 |
| 22 | 111.965 | 131.462 | 84.762 | 115.234 | 107.645 |
| 23 | 93.401 | 108.579 | 106.477 | 119.203 | 120.254 |
| 24 | 117.802 | 117.452 | 103.909 | 82.076 | 82.31 |
| 25 | 77.29 | 112.665 | 108.579 | 114.183 | 115.467 |
| 26 | 76.005 | 126.325 | 122.122 | 84.645 | 74.371 |
| 27 | 111.498 | 159.599 | 84.411 | 99.706 | 113.482 |
| 28 | 103.325 | 98.538 | 130.528 | 98.655 | 127.142 |
| 29 | 81.026 | 94.335 | 94.452 | 91.767 | 131.696 |
| 30 | 91.183 | 132.396 | 102.858 | 113.949 | 113.132 |
| 31 |  | 97.021 | 123.173 |  | 88.965 |
| 32 |  |  |  |  | 153.295 |

*nos-gal4, tub-gal80<sup>ts</sup> driven Set1 RNAi Data:*
| **Testis #** | **Day 0** | **Day 7** | **Day 14** | **Day 21** | **Day 28** |
| --- | --- | --- | --- | --- | --- |
| 1 | 102.158 | 112.432 | 138.234 | 147.924 | 192.29 |
| 2 | 89.432 | 112.782 | 98.422 | 173.259 | 150.843 |
| 3 | 109.279 | 160.65 | 169.056 | 171.041 | 180.498 |
| 4 | 95.97 | 105.076 | 96.087 | 157.498 | 204.899 |
| 5 | 67.249 | 106.711 | 116.985 | 177.462 | 203.731 |
| 6 | 95.036 | 105.193 | 122.939 | 206.884 | 133.214 |
| 7 | 109.98 | 166.604 | 191.122 | 170.107 | 115.117 |
| 8 | 89.432 | 103.208 | 197.66 | 129.477 | 242.259 |
| 9 | 68.066 | 94.218 | 163.802 | 207.701 | 215.523 |
| 10 | 61.762 | 123.64 | 109.513 | 193.924 | 164.736 |
| 11 | 99.939 | 70.051 | 156.564 | 133.797 | 147.107 |
| 12 | 96.904 | 132.279 | 202.097 | 199.061 | 163.219 |
| 13 | 53.589 | 141.503 | 148.625 | 123.64 | 672.139 |
| 14 | 119.437 | 131.929 | 130.528 | 68.65 | 139.752 |
| 15 | 90.249 | 119.67 | 164.153 | 177.813 | 148.391 |
| 16 | 102.858 | 119.203 | 137.066 | 223.346 | 100.873 |
| 17 | 97.371 | 137.183 | 162.868 | 102.858 | 119.087 |
| 18 | 75.071 | 130.528 | 154.112 | 88.848 | 132.63 |
| 19 | 104.259 | 121.655 | 157.264 | 194.625 | 92.934 |
| 20 | 69.584 | 111.264 | 118.269 | 163.919 | 114.884 |
| 21 | 55.107 | 90.482 | 123.056 | 137.066 | 120.021 |
| 22 | 111.147 | 103.559 | 127.259 | 223.229 | 125.391 |
| 23 | 109.63 |  | 134.848 | 121.305 | 139.985 |
| 24 | 78.224 |  | 158.899 | 74.02 | 171.975 |
| 25 | 87.447 |  | 123.757 | 106.828 | 137.767 |
| 26 | 71.919 |  | 98.422 | 148.508 | 167.655 |
| 27 | 117.919 |  | 125.625 | 155.396 |  |
| 28 | 109.279 |  | 229.3 | 112.782 |  |
| 29 | 78.924 |  | 112.665 |  |  |
| 30 | 84.995 |  | 158.549 |  |  |
| 31 | 69 |  | 158.549 |  |  |
| 32 | 53.355 |  | 205.366 |  |  |
| 33 | 76.472 |  | 104.726 |  |  |
| 34 | 95.736 |  |  |  |  |
| 35 | 90.366 |  |  |  |  |

*tj driven RNAi Data:*
| **Testis #** | ***Control RNAi*** | ***Set1 RNAi*** |
| --- | --- | --- |
| 1 | 85.345 | 73.67 |
| 2 | 79.741 | 59.193 |
| 3 | 116.168 | 87.213 |
| 4 | 121.422 | 95.269 |
| 5 | 101.807 | 74.254 |
| 6 | 128.076 | 130.178 |
| 7 | 84.995 | 88.264 |
| 8 | 73.67 | 67.482 |
| 9 | 86.863 | 75.888 |
| 10 | 93.518 | 28.604 |
| 11 | 65.381 | 155.747 |
| 12 | 117.569 | 112.899 |
| 13 | 76.823 | 92.701 |
| 14 | 84.411 | 82.66 |
| 15 | 83.944 | 99.356 |
| 16 | 139.635 | 83.594 |
| 17 | 107.878 | 82.427 |
| 18 | 84.995 | 98.188 |
| 19 | 113.132 | 84.878 |
| 20 | 78.807 | 120.955 |
| 21 | 81.026 | 114.3 |
| 22 | 107.178 | 88.965 |
| 23 | 102.975 | 74.604 |
| 24 | 84.645 | 93.985 |
| 25 | 100.873 | 118.853 |
| 26 | 110.564 | 93.868 |
| 27 | 88.965 | 83.711 |
| 28 | 116.635 | 90.716 |
| 29 | 82.777 | 110.914 |
| 30 | 94.335 | 96.437 |
| 31 | 107.061 | 87.68 |
| 32 | 101.224 | 110.68 |
| 33 | 121.772 | 90.482 |
| 34 | 101.807 | 95.97 |
| 35 | 98.422 | 84.645 |
| 36 | 79.625 | 98.889 |
| 37 | 129.944 |  |
| 38 | 86.863 |  |
| 39 | 82.777 |  |
| 40 | 81.493 |  |
| 41 | 77.173 |  |
| 42 | 89.081 |  |
| 43 | 134.264 |  |
| 44 | 109.513 |  |
| 45 | 86.63 |  |
| 46 | 74.954 |  |

*bam driven RNAi Data:*
| **Testis #** | ***Control RNAi*** | ***Set1 RNAi*** |
| --- | --- | --- |
| 1 | 119.67 | 92.35 |
| 2 | 127.259 | 120.721 |
| 3 | 87.213 | 144.305 |
| 4 | 123.173 | 107.178 |
| 5 | 142.904 | 162.635 |
| 6 | 121.772 | 141.386 |
| 7 | 117.452 | 113.366 |
| 8 | 141.62 | 106.244 |
| 9 | 115.818 | 83.711 |
| 10 | 99.589 | 100.29 |
| 11 | 167.772 | 126.442 |
| 12 | 125.274 | 113.482 |
| 13 | 118.736 | 105.894 |
| 14 | 81.96 | 78.457 |
| 15 | 115.584 | 146.873 |
| 16 | 128.076 | 87.914 |
| 17 | 155.747 | 110.33 |
| 18 | 84.411 | 135.549 |
| 19 | 110.33 | 126.442 |
| 20 | 118.386 | 138.351 |
| 21 | 127.259 | 126.909 |
| 22 | 95.503 | 137.533 |
| 23 | 104.609 | 107.295 |
| 24 | 98.889 | 128.894 |
| 25 | 198.478 | 88.264 |
| 26 | 163.919 | 131.579 |
| 27 | 134.848 | 101.807 |
| 28 | 100.29 | 114.884 |
| 29 | 98.538 | 99.589 |
| 30 | 129.944 | 96.67 |
| 31 | 113.015 | 98.538 |
| 32 |  | 140.919 |
| 33 |  | 83.361 |
| 34 |  | 114.3 |
| 35 |  | 150.96 |
| 36 |  | 148.158 |
| 37 |  | 118.736 |
| 38 |  | 137.533 |
| 39 |  | 114.65 |
| 40 |  | 167.772 |
| 41 |  | 103.792 |
| 42 |  | 129.244 |
| 43 |  | 168.472 |
| 44 |  | 135.198 |
| 45 |  | 83.477 |
| 46 |  | 128.66 |
| 47 |  | 102.041 |
| 48 |  | 120.488 |
| 49 |  | 93.985 |
| 50 |  | 115 |

*nos-gal4, bam-gal80 driven RNAi Data:*
| **Testis #** | ***Control RNAi*** | ***Set1 RNAi*** |
| --- | --- | --- |
| 1 | 59.591 | 109.663 |
| 2 | 103.665 | 92.581 |
| 3 | 95.45 | 94.798 |
| 4 | 76.412 | 107.576 |
| 5 | 85.54 | 95.58 |
| 6 | 84.366 | 133.395 |
| 7 | 98.449 | 86.713 |
| 8 | 71.326 | 91.668 |
| 9 | 142.001 | 129.874 |
| 10 | 133.004 | 129.744 |
| 11 | 140.045 | 103.925 |
| 12 | 89.191 | 94.146 |
| 13 | 88.539 | 99.643 |
| 14 | 86.713 | 101.314 |
| 15 | 83.975 | 117.63 |
| 16 | 103.404 | 98.051 |
| 17 | 103.404 | 82.671 |
| 18 | 89.321 | 97.406 |
| 19 | 126.614 | 95.971 |
| 20 | 83.975 | 129.222 |
| 21 | 94.276 | 81.758 |
| 22 | 119.703 | 131.83 |
| 23 | 100.144 | 193.768 |
| 24 | 87.626 | 85.018 |
| 25 | 83.714 | 56.722 |
| 26 | 101.969 | 100.405 |
| 27 | 89.712 | 113.053 |
| 28 | 73.152 | 76.673 |
| 29 | 100.665 | 108.228 |
| 30 | 85.8 | 120.094 |
| 31 | 114.618 | 76.021 |
| 32 | 80.861 | 99.361 |
| 33 | 104.02 | 84.105 |
| 34 | 99.086 | 120.746 |
| 35 | 138.72 | 449.213 |
| 36 | 126.384 | 906.626 |
| 37 | 120.415 | 244.622 |
| 38 | 111.024 | 107.707 |
| 39 | 73.152 | 181.771 |
| 40 | 103.795 | 143.696 |
| 41 | 82.28 | 99.753 |
| 42 | 78.237 | 83.453 |
| 43 | 79.933 | 135.611 |
| 44 | 98.709 | 104.186 |
| 45 | 115.922 | 79.15 |
| 46 | 73.804 | 127.918 |
| 47 | 93.754 | 89.191 |
| 48 | 106.664 | 108.489 |
| 49 | 80.845 | 123.876 |
| 50 | 101.578 | 107.968 |
| 51 | 89.843 | 107.185 |
| 52 | 127.527 | 101.709 |

**Table S3:**
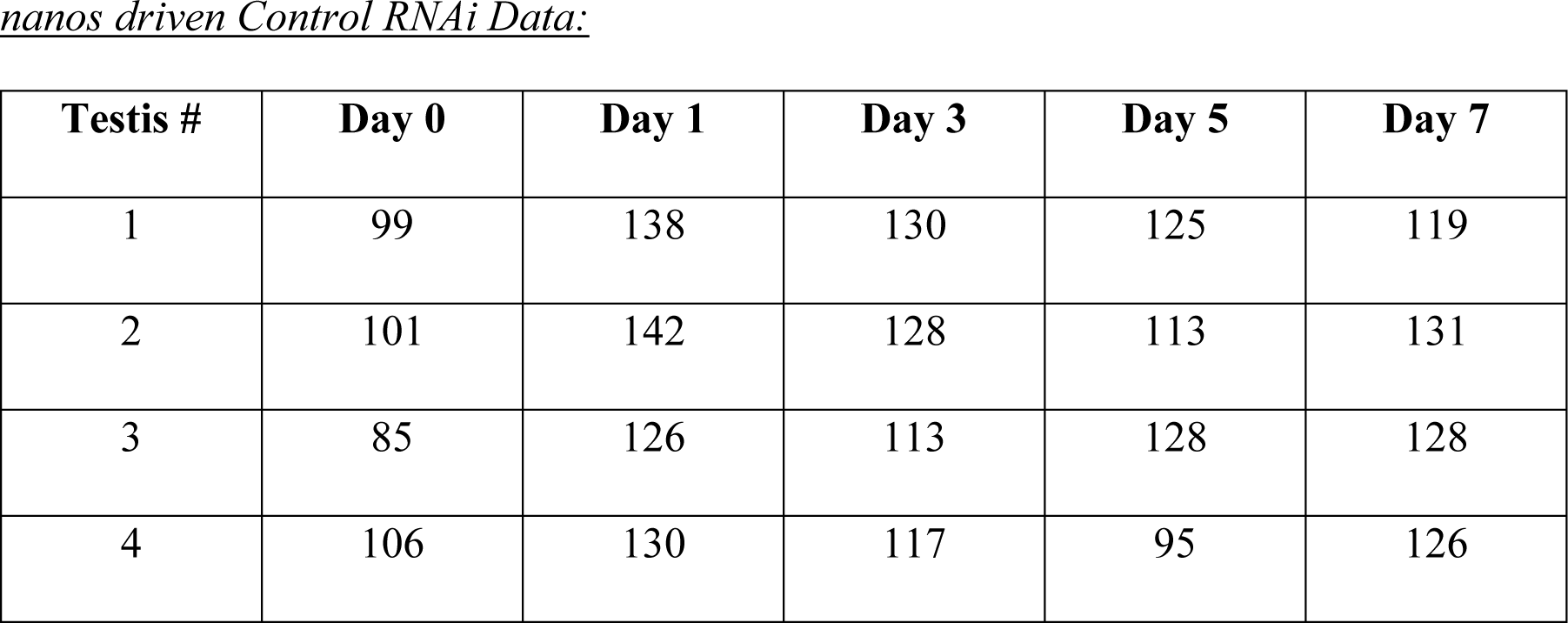
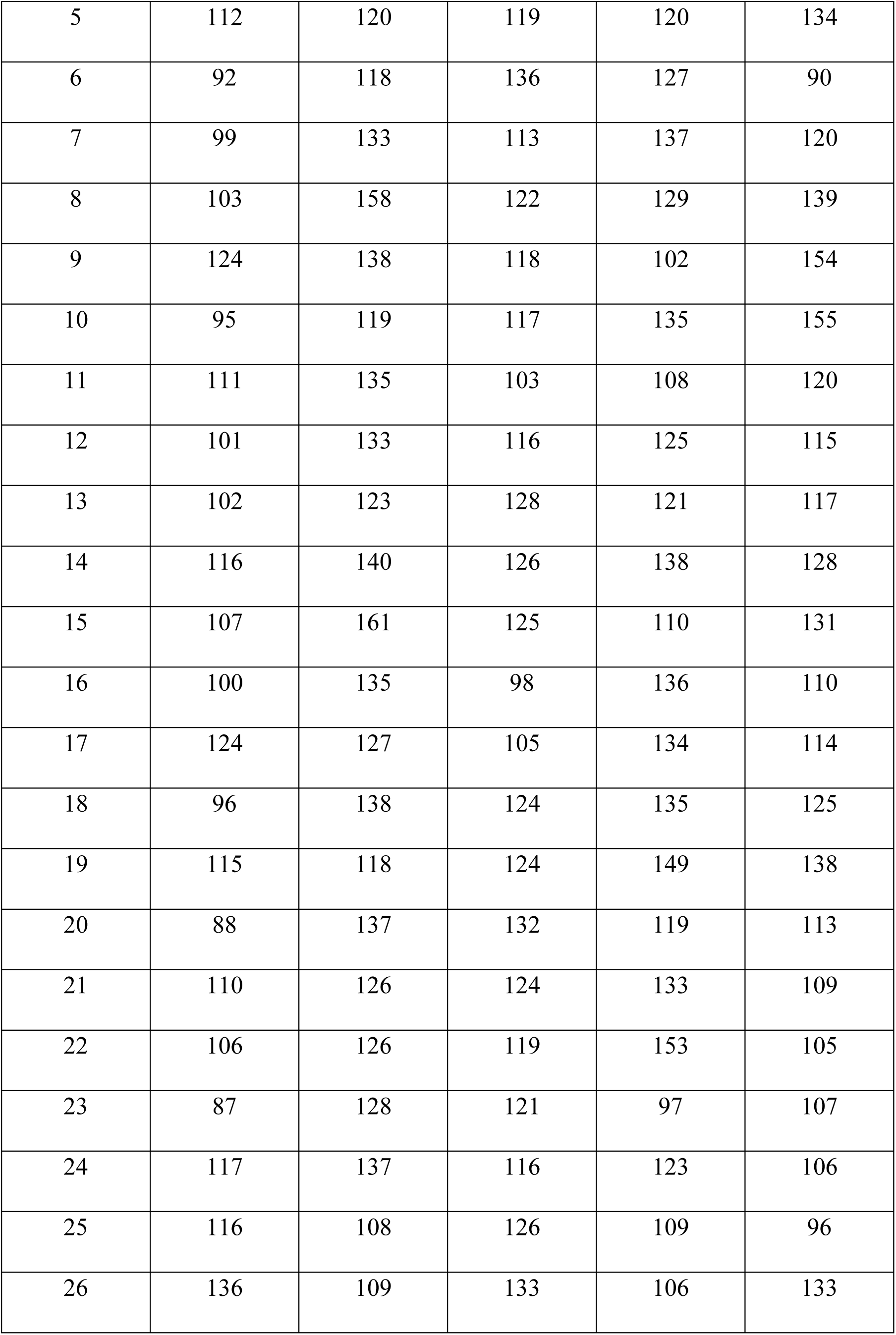
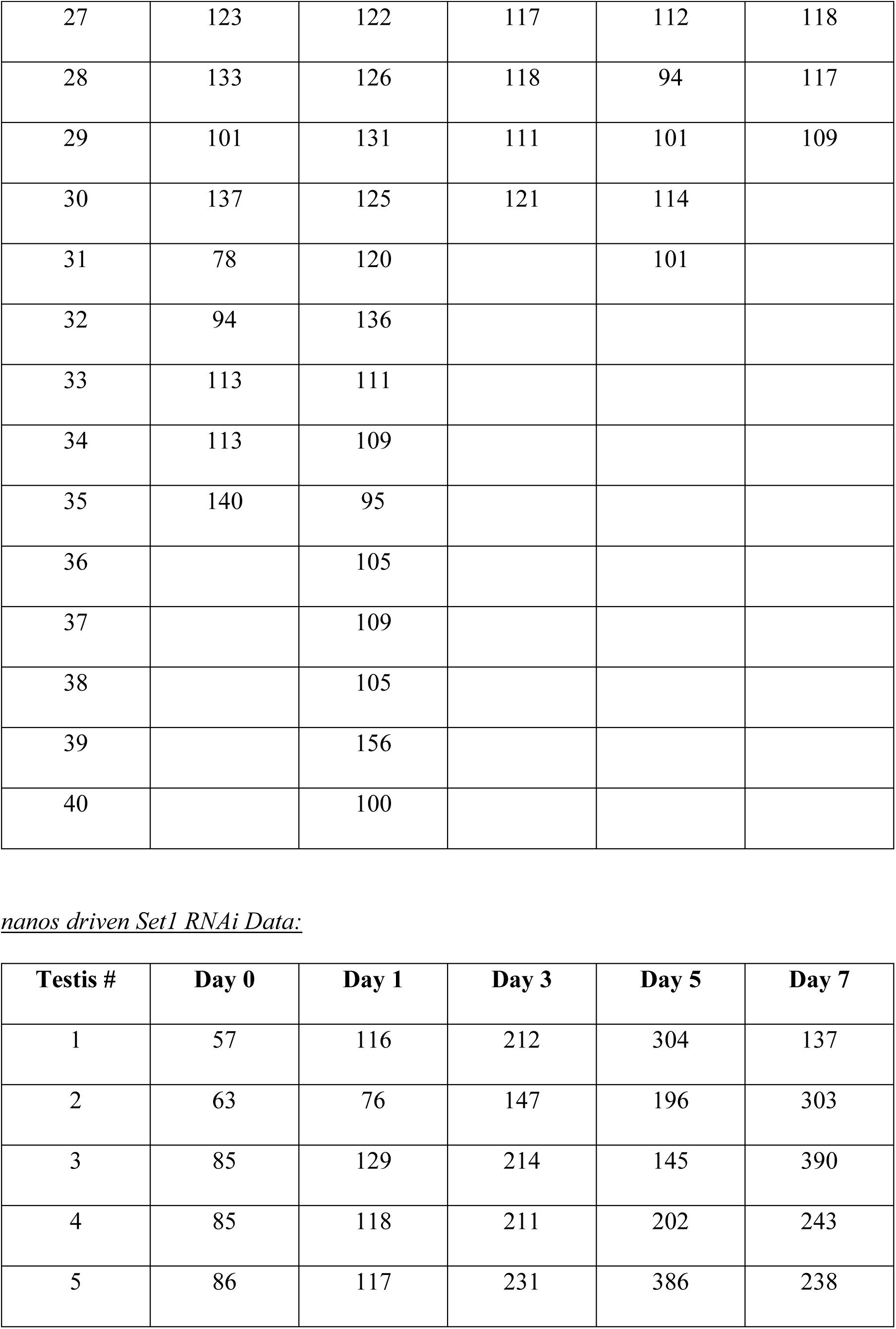
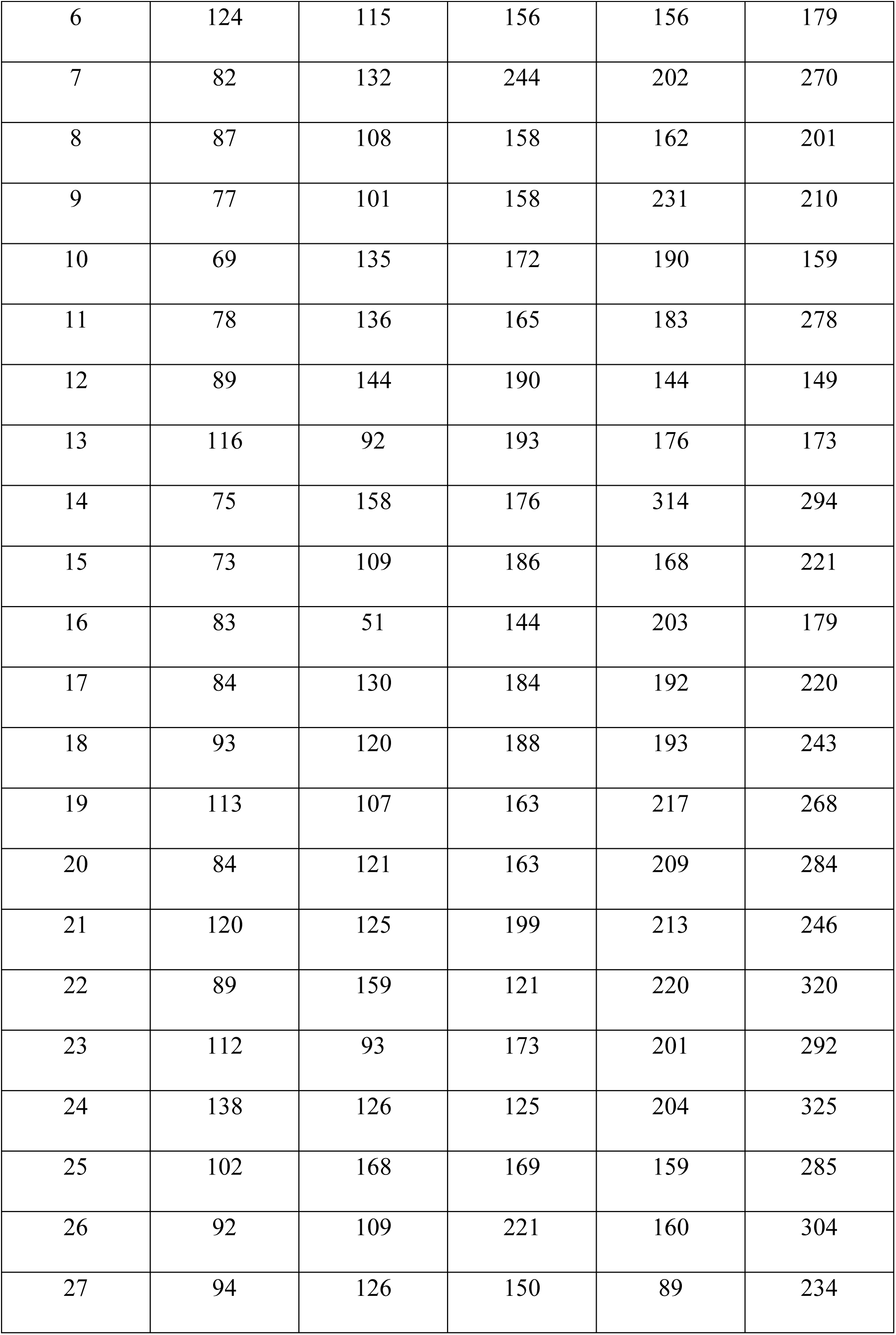
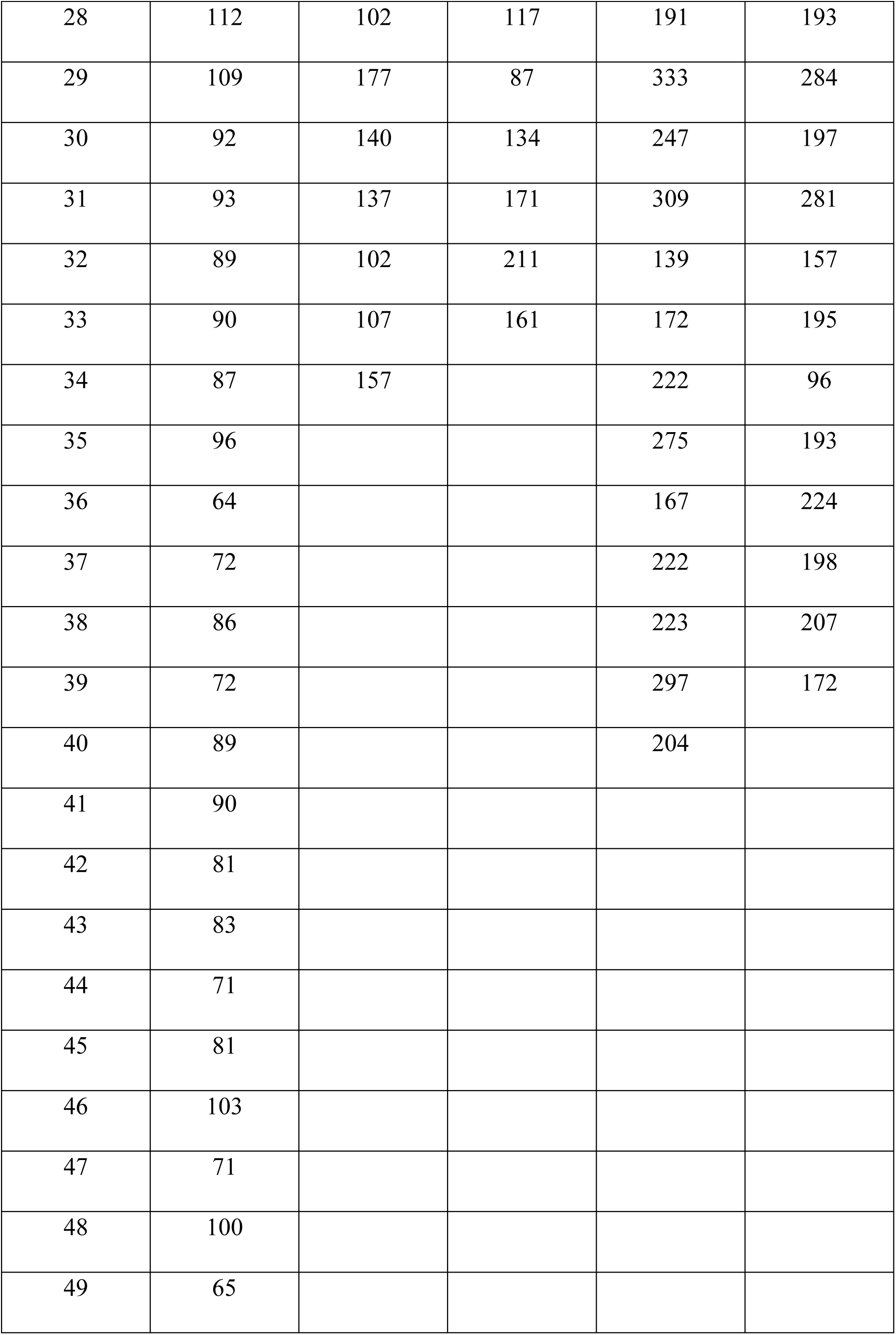
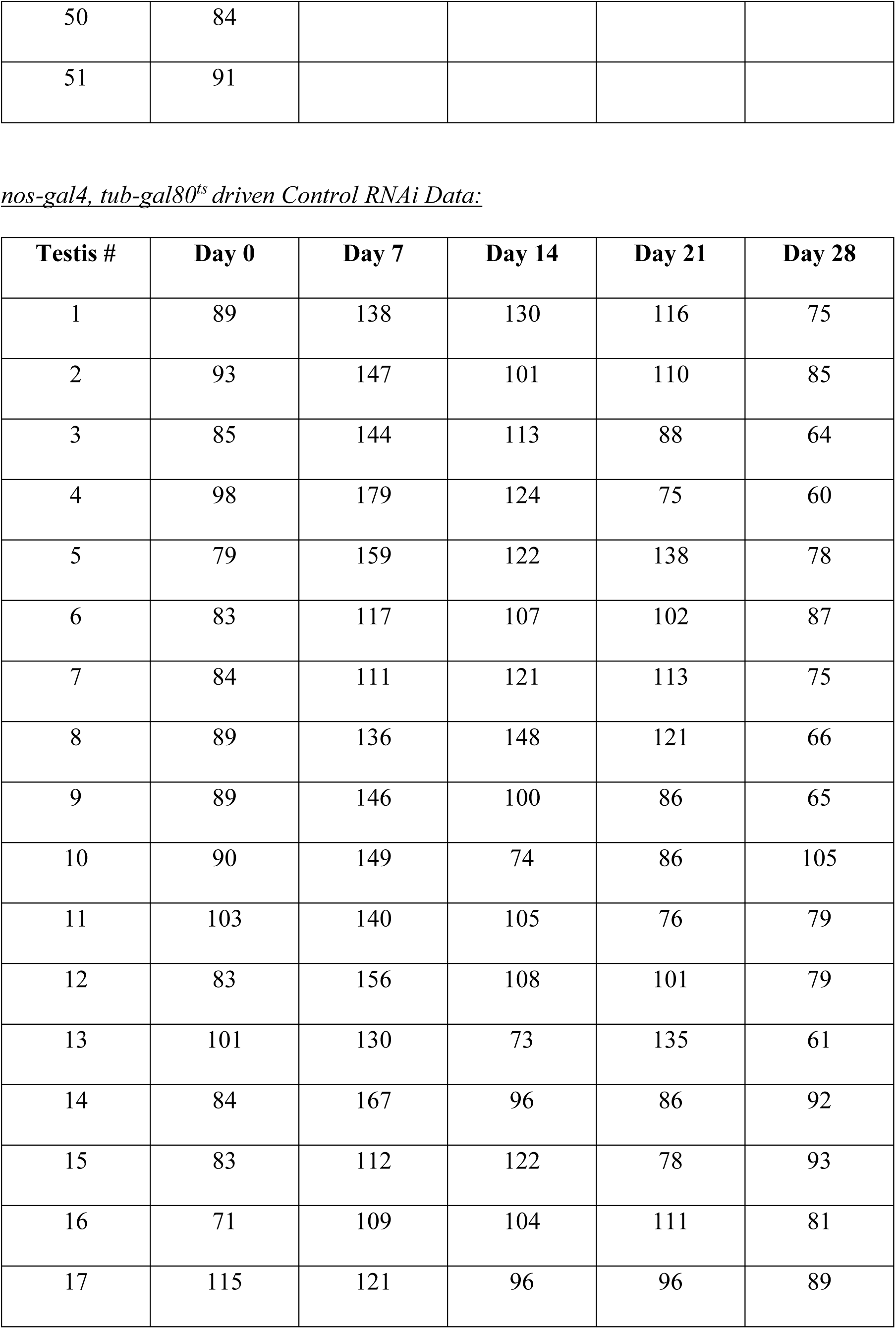
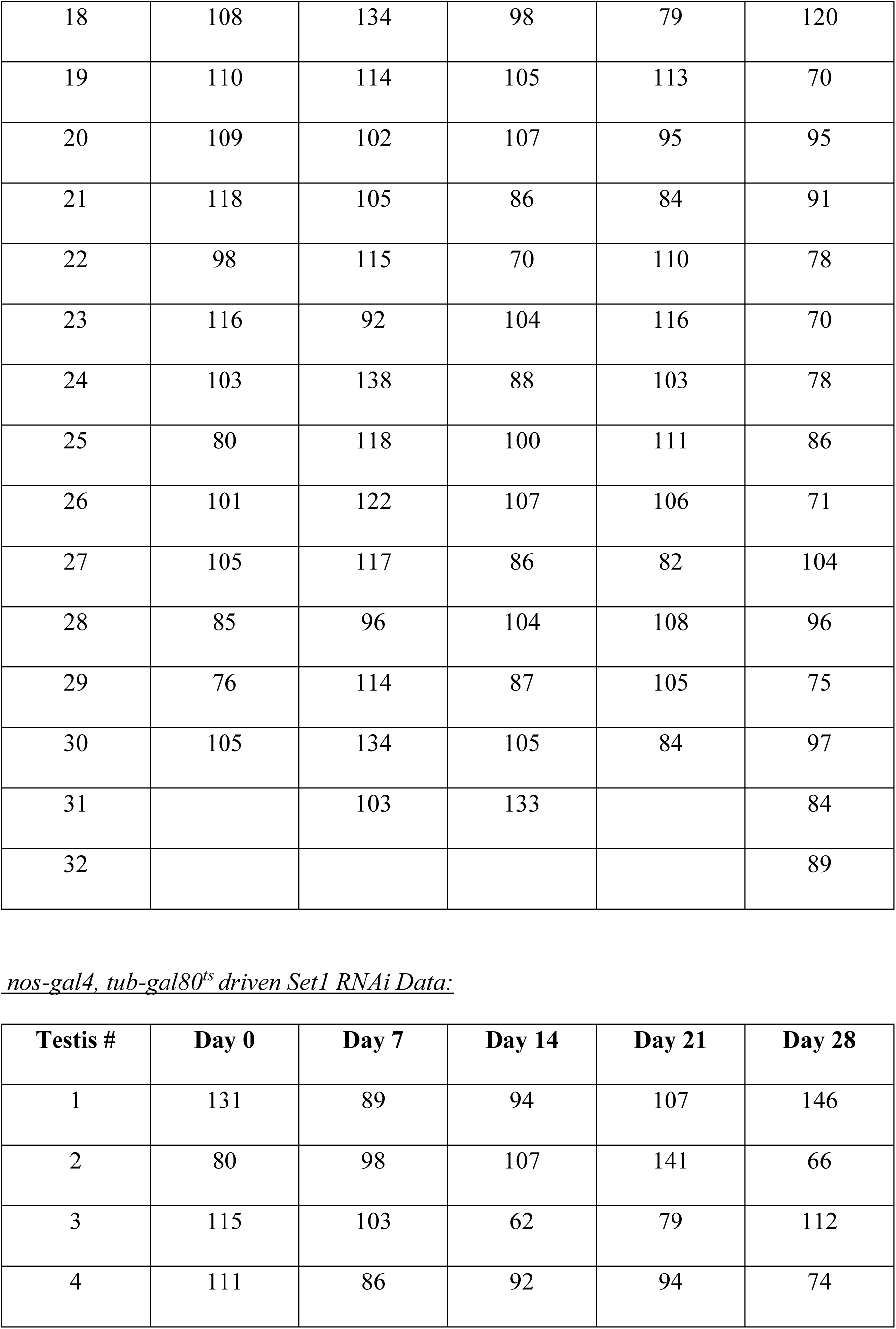
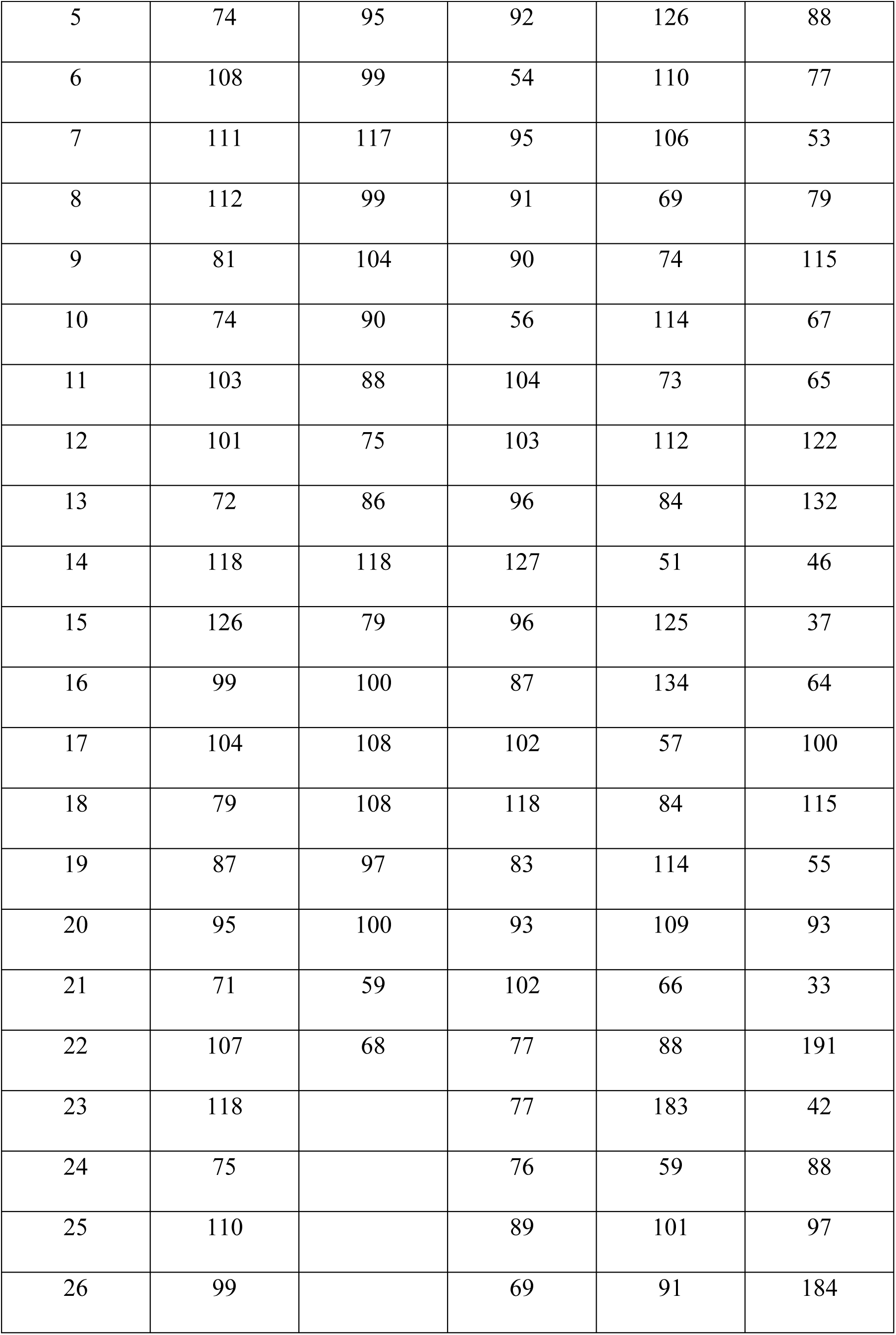
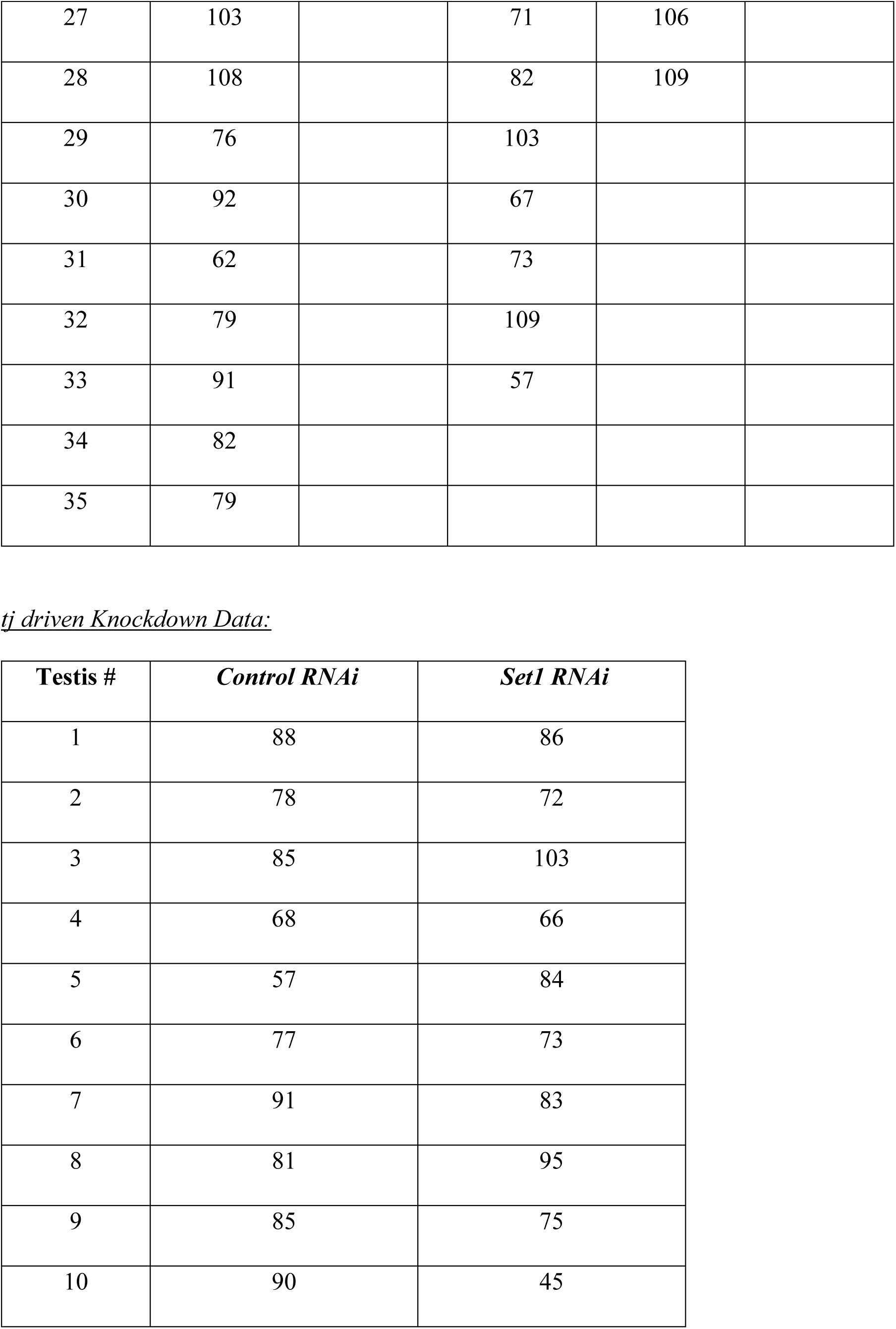
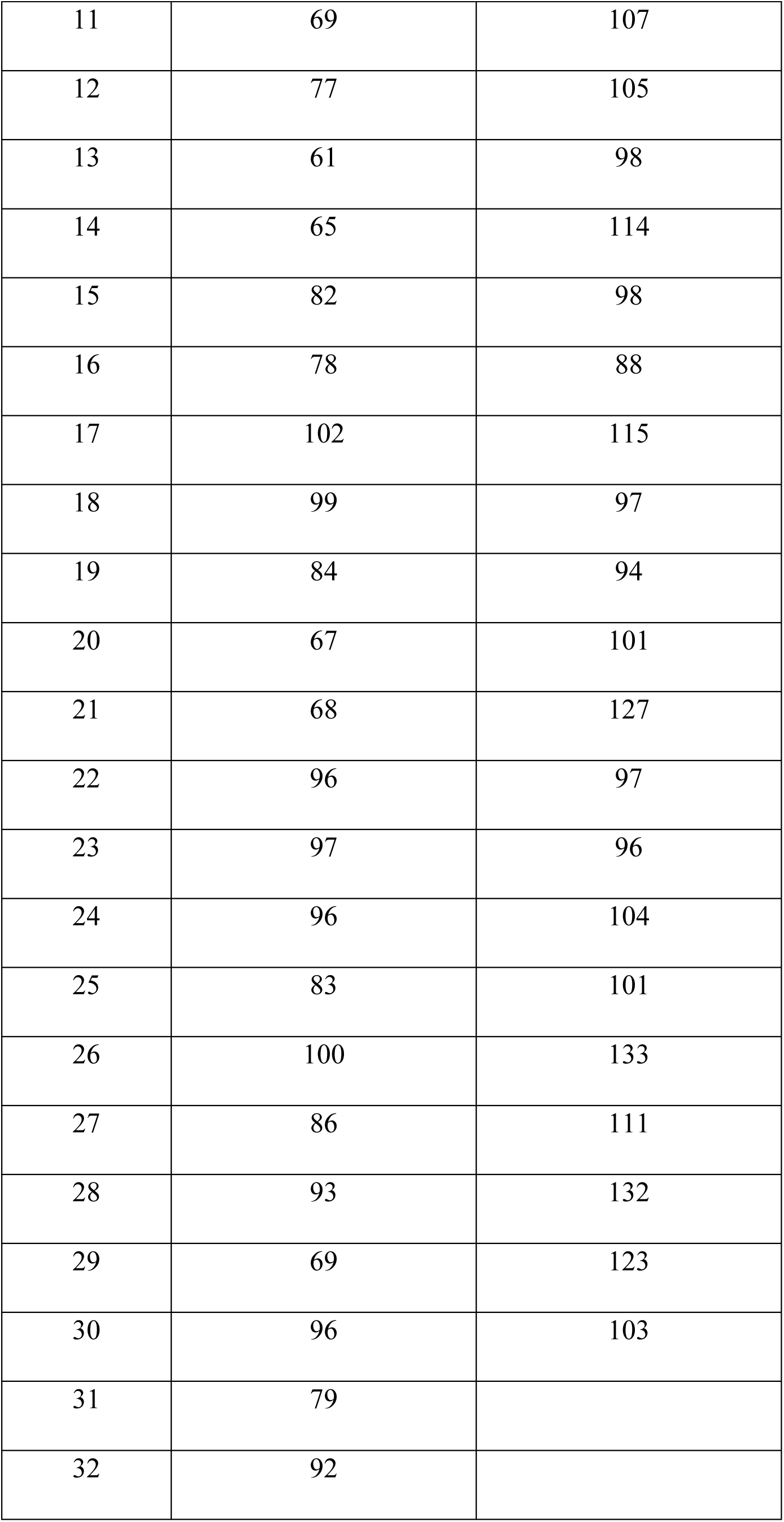
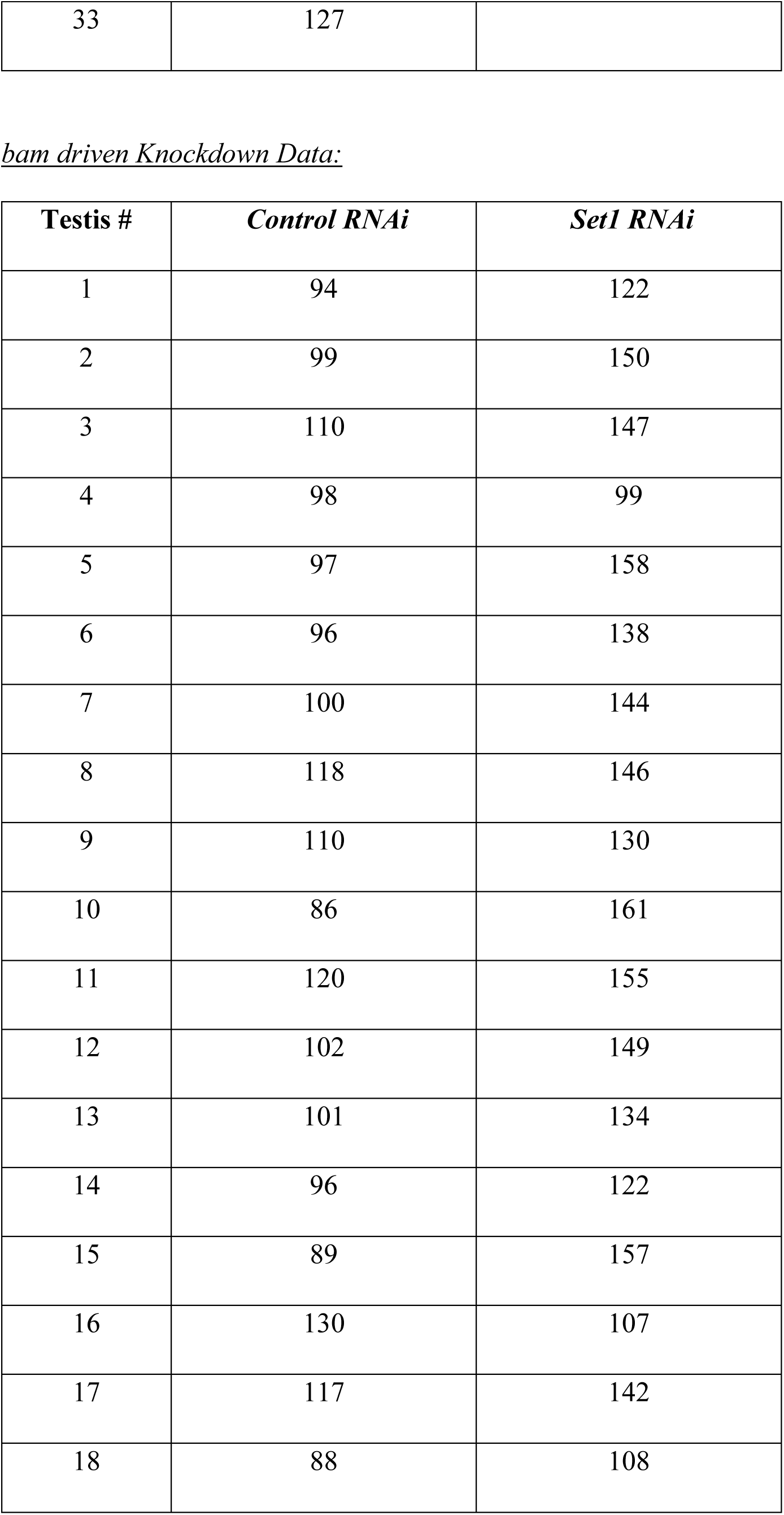
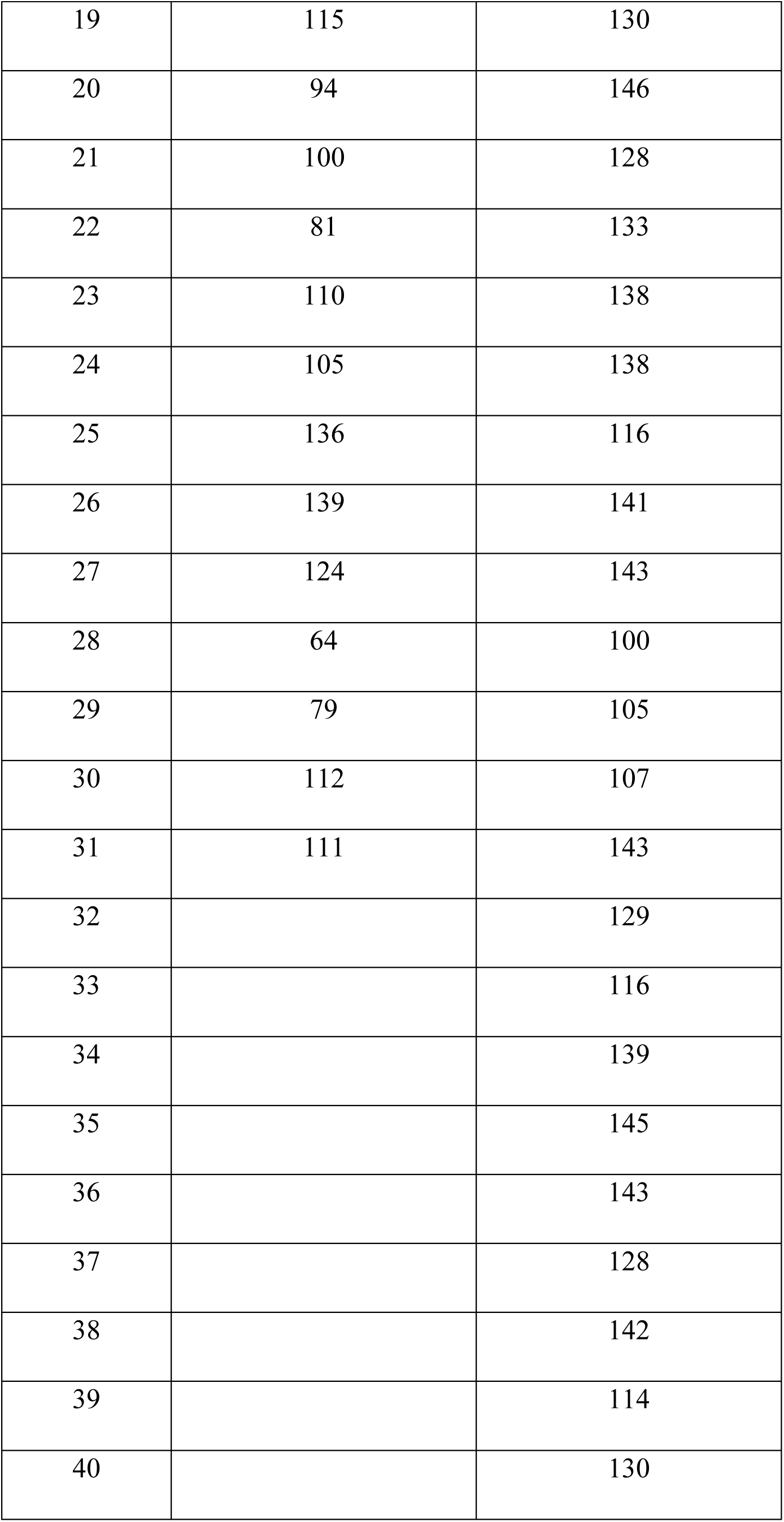
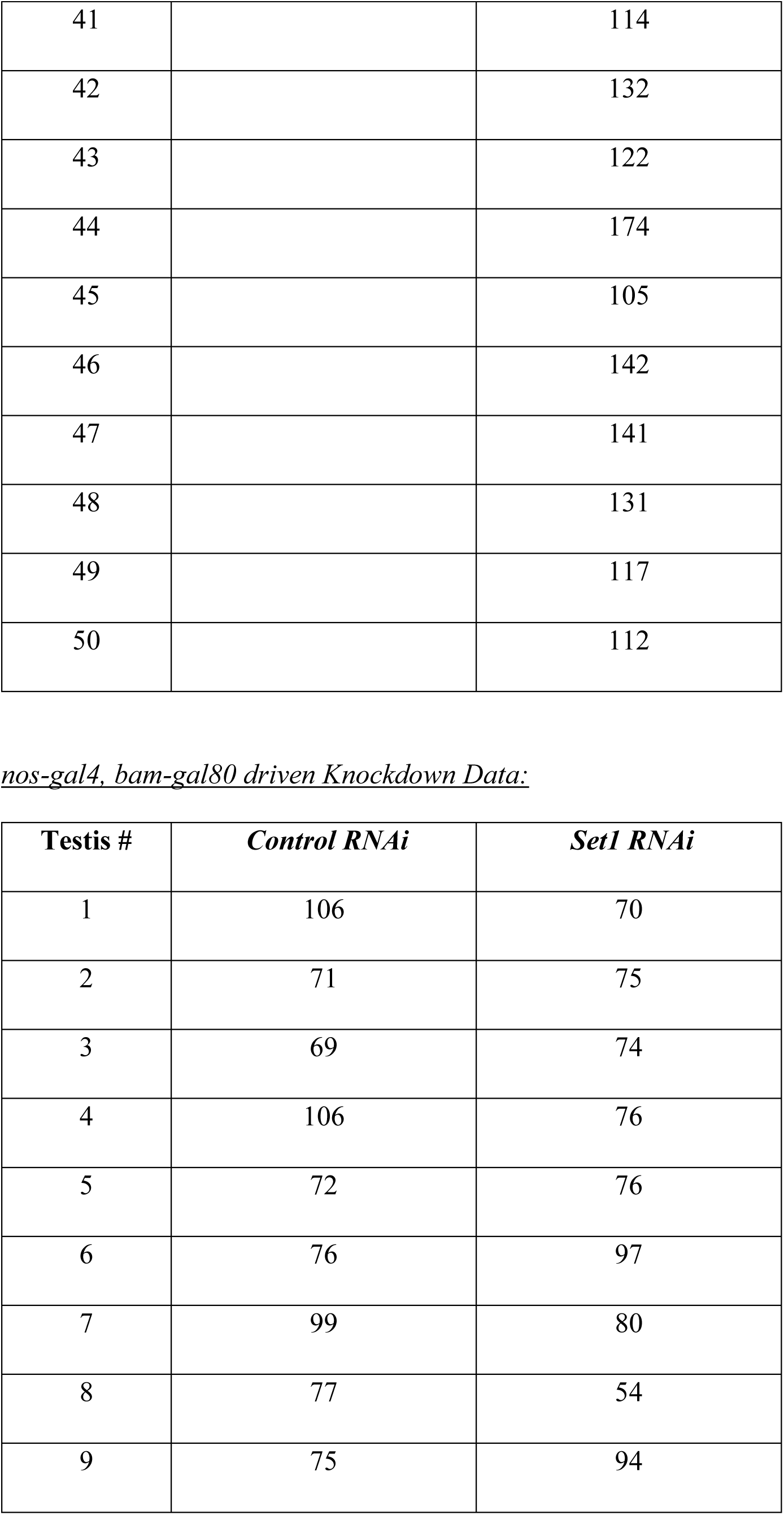
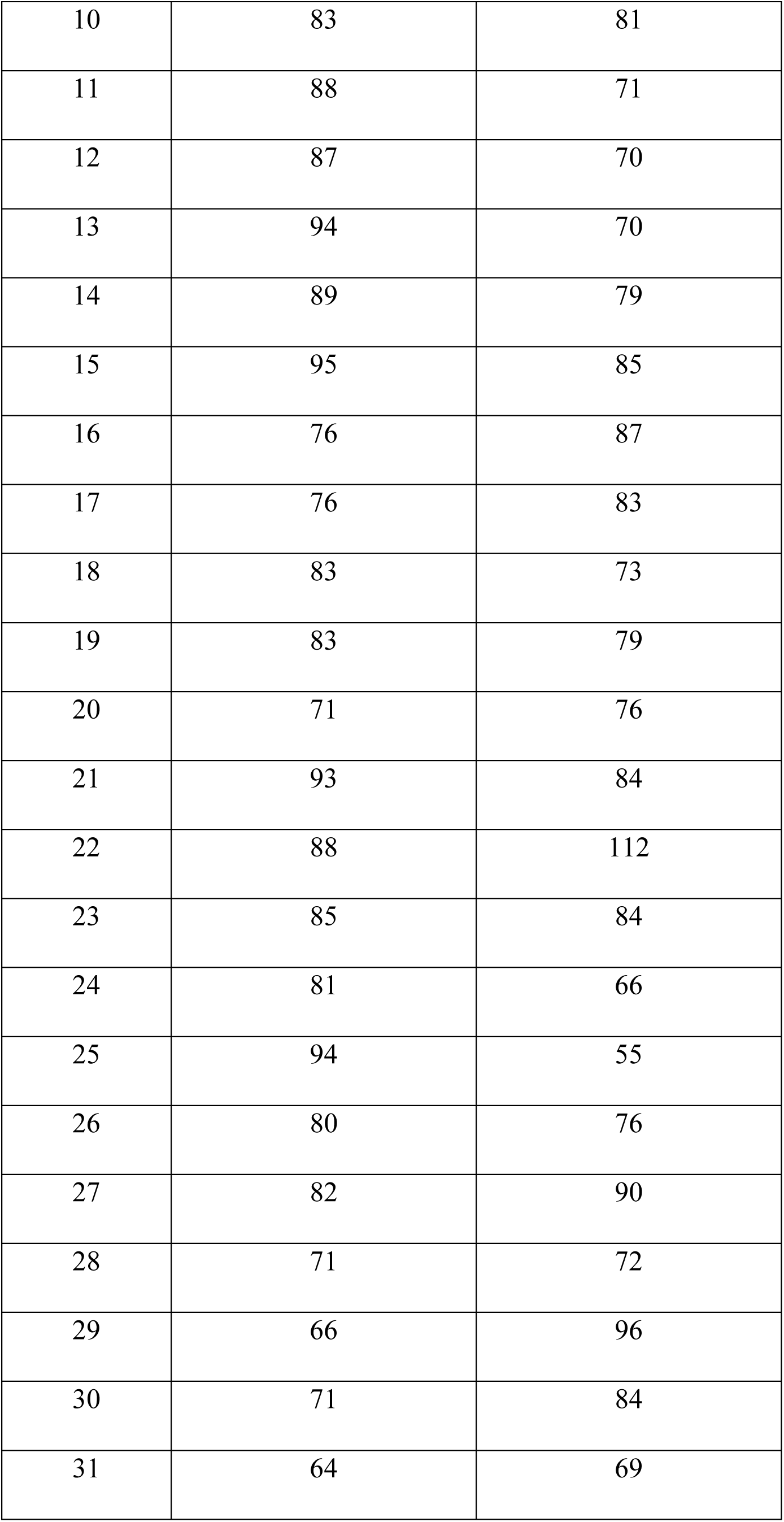
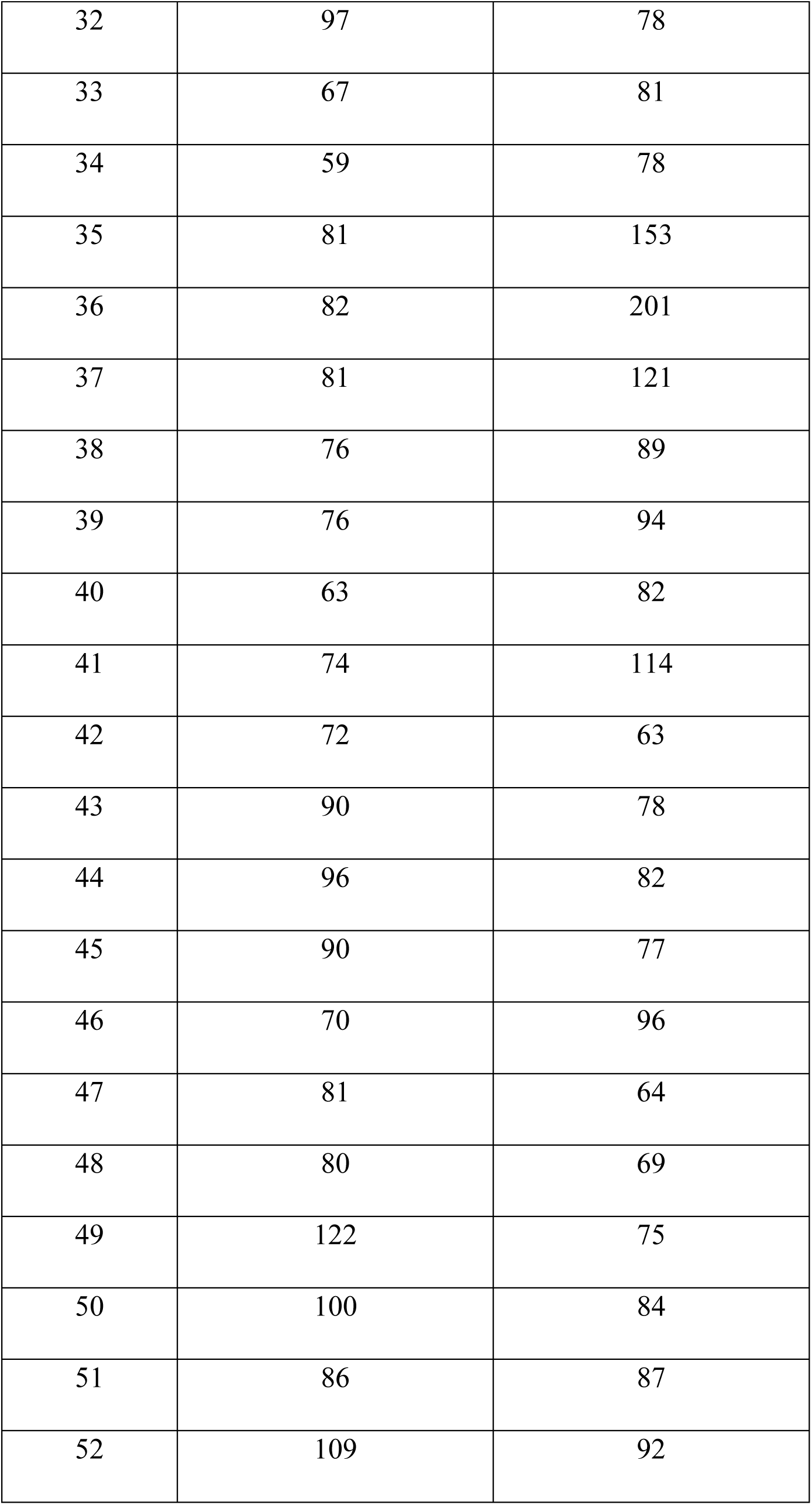
Quantification of Cyst Cell number in RNAi knockdown testes.

| **Testis #** | **Day 0** | **Day 1** | **Day 3** | **Day 5** | **Day 7** |
| --- | --- | --- | --- | --- | --- |
| 1 | 99 | 138 | 130 | 125 | 119 |
| 2 | 101 | 142 | 128 | 113 | 131 |
| 3 | 85 | 126 | 113 | 128 | 128 |
| 4 | 106 | 130 | 117 | 95 | 126 |
| 5 | 112 | 120 | 119 | 120 | 134 |
| 6 | 92 | 118 | 136 | 127 | 90 |
| 7 | 99 | 133 | 113 | 137 | 120 |
| 8 | 103 | 158 | 122 | 129 | 139 |
| 9 | 124 | 138 | 118 | 102 | 154 |
| 10 | 95 | 119 | 117 | 135 | 155 |
| 11 | 111 | 135 | 103 | 108 | 120 |
| 12 | 101 | 133 | 116 | 125 | 115 |
| 13 | 102 | 123 | 128 | 121 | 117 |
| 14 | 116 | 140 | 126 | 138 | 128 |
| 15 | 107 | 161 | 125 | 110 | 131 |
| 16 | 100 | 135 | 98 | 136 | 110 |
| 17 | 124 | 127 | 105 | 134 | 114 |
| 18 | 96 | 138 | 124 | 135 | 125 |
| 19 | 115 | 118 | 124 | 149 | 138 |
| 20 | 88 | 137 | 132 | 119 | 113 |
| 21 | 110 | 126 | 124 | 133 | 109 |
| 22 | 106 | 126 | 119 | 153 | 105 |
| 23 | 87 | 128 | 121 | 97 | 107 |
| 24 | 117 | 137 | 116 | 123 | 106 |
| 25 | 116 | 108 | 126 | 109 | 96 |
| 26 | 136 | 109 | 133 | 106 | 133 |
| 27 | 123 | 122 | 117 | 112 | 118 |
| 28 | 133 | 126 | 118 | 94 | 117 |
| 29 | 101 | 131 | 111 | 101 | 109 |
| 30 | 137 | 125 | 121 | 114 |  |
| 31 | 78 | 120 |  | 101 |  |
| 32 | 94 | 136 |  |  |  |
| 33 | 113 | 111 |  |  |  |
| 34 | 113 | 109 |  |  |  |
| 35 | 140 | 95 |  |  |  |
| 36 |  | 105 |  |  |  |
| 37 |  | 109 |  |  |  |
| 38 |  | 105 |  |  |  |
| 39 |  | 156 |  |  |  |
| 40 |  | 100 |  |  |  |

*nanos driven Set1 RNAi Data:*
| **Testis #** | **Day 0** | **Day 1** | **Day 3** | **Day 5** | **Day 7** |
| --- | --- | --- | --- | --- | --- |
| 1 | 57 | 116 | 212 | 304 | 137 |
| 2 | 63 | 76 | 147 | 196 | 303 |
| 3 | 85 | 129 | 214 | 145 | 390 |
| 4 | 85 | 118 | 211 | 202 | 243 |
| 5 | 86 | 117 | 231 | 386 | 238 |
| 6 | 124 | 115 | 156 | 156 | 179 |
| 7 | 82 | 132 | 244 | 202 | 270 |
| 8 | 87 | 108 | 158 | 162 | 201 |
| 9 | 77 | 101 | 158 | 231 | 210 |
| 10 | 69 | 135 | 172 | 190 | 159 |
| 11 | 78 | 136 | 165 | 183 | 278 |
| 12 | 89 | 144 | 190 | 144 | 149 |
| 13 | 116 | 92 | 193 | 176 | 173 |
| 14 | 75 | 158 | 176 | 314 | 294 |
| 15 | 73 | 109 | 186 | 168 | 221 |
| 16 | 83 | 51 | 144 | 203 | 179 |
| 17 | 84 | 130 | 184 | 192 | 220 |
| 18 | 93 | 120 | 188 | 193 | 243 |
| 19 | 113 | 107 | 163 | 217 | 268 |
| 20 | 84 | 121 | 163 | 209 | 284 |
| 21 | 120 | 125 | 199 | 213 | 246 |
| 22 | 89 | 159 | 121 | 220 | 320 |
| 23 | 112 | 93 | 173 | 201 | 292 |
| 24 | 138 | 126 | 125 | 204 | 325 |
| 25 | 102 | 168 | 169 | 159 | 285 |
| 26 | 92 | 109 | 221 | 160 | 304 |
| 27 | 94 | 126 | 150 | 89 | 234 |
| 28 | 112 | 102 | 117 | 191 | 193 |
| 29 | 109 | 177 | 87 | 333 | 284 |
| 30 | 92 | 140 | 134 | 247 | 197 |
| 31 | 93 | 137 | 171 | 309 | 281 |
| 32 | 89 | 102 | 211 | 139 | 157 |
| 33 | 90 | 107 | 161 | 172 | 195 |
| 34 | 87 | 157 |  | 222 | 96 |
| 35 | 96 |  |  | 275 | 193 |
| 36 | 64 |  |  | 167 | 224 |
| 37 | 72 |  |  | 222 | 198 |
| 38 | 86 |  |  | 223 | 207 |
| 39 | 72 |  |  | 297 | 172 |
| 40 | 89 |  |  | 204 |  |
| 41 | 90 |  |  |  |  |
| 42 | 81 |  |  |  |  |
| 43 | 83 |  |  |  |  |
| 44 | 71 |  |  |  |  |
| 45 | 81 |  |  |  |  |
| 46 | 103 |  |  |  |  |
| 47 | 71 |  |  |  |  |
| 48 | 100 |  |  |  |  |
| 49 | 65 |  |  |  |  |
| 50 | 84 |  |  |  |  |
| 51 | 91 |  |  |  |  |

*nos-gal4, tub-gal80<sup>ts</sup> driven Control RNAi Data:*
| **Testis #** | **Day 0** | **Day 7** | **Day 14** | **Day 21** | **Day 28** |
| --- | --- | --- | --- | --- | --- |
| 1 | 89 | 138 | 130 | 116 | 75 |
| 2 | 93 | 147 | 101 | 110 | 85 |
| 3 | 85 | 144 | 113 | 88 | 64 |
| 4 | 98 | 179 | 124 | 75 | 60 |
| 5 | 79 | 159 | 122 | 138 | 78 |
| 6 | 83 | 117 | 107 | 102 | 87 |
| 7 | 84 | 111 | 121 | 113 | 75 |
| 8 | 89 | 136 | 148 | 121 | 66 |
| 9 | 89 | 146 | 100 | 86 | 65 |
| 10 | 90 | 149 | 74 | 86 | 105 |
| 11 | 103 | 140 | 105 | 76 | 79 |
| 12 | 83 | 156 | 108 | 101 | 79 |
| 13 | 101 | 130 | 73 | 135 | 61 |
| 14 | 84 | 167 | 96 | 86 | 92 |
| 15 | 83 | 112 | 122 | 78 | 93 |
| 16 | 71 | 109 | 104 | 111 | 81 |
| 17 | 115 | 121 | 96 | 96 | 89 |
| 18 | 108 | 134 | 98 | 79 | 120 |
| 19 | 110 | 114 | 105 | 113 | 70 |
| 20 | 109 | 102 | 107 | 95 | 95 |
| 21 | 118 | 105 | 86 | 84 | 91 |
| 22 | 98 | 115 | 70 | 110 | 78 |
| 23 | 116 | 92 | 104 | 116 | 70 |
| 24 | 103 | 138 | 88 | 103 | 78 |
| 25 | 80 | 118 | 100 | 111 | 86 |
| 26 | 101 | 122 | 107 | 106 | 71 |
| 27 | 105 | 117 | 86 | 82 | 104 |
| 28 | 85 | 96 | 104 | 108 | 96 |
| 29 | 76 | 114 | 87 | 105 | 75 |
| 30 | 105 | 134 | 105 | 84 | 97 |
| 31 |  | 103 | 133 |  | 84 |
| 32 |  |  |  |  | 89 |

*nos-gal4, tub-gal80<sup>ts</sup> driven Set1 RNAi Data:*
| **Testis #** | **Day 0** | **Day 7** | **Day 14** | **Day 21** | **Day 28** |
| --- | --- | --- | --- | --- | --- |
| 1 | 131 | 89 | 94 | 107 | 146 |
| 2 | 80 | 98 | 107 | 141 | 66 |
| 3 | 115 | 103 | 62 | 79 | 112 |
| 4 | 111 | 86 | 92 | 94 | 74 |
| 5 | 74 | 95 | 92 | 126 | 88 |
| 6 | 108 | 99 | 54 | 110 | 77 |
| 7 | 111 | 117 | 95 | 106 | 53 |
| 8 | 112 | 99 | 91 | 69 | 79 |
| 9 | 81 | 104 | 90 | 74 | 115 |
| 10 | 74 | 90 | 56 | 114 | 67 |
| 11 | 103 | 88 | 104 | 73 | 65 |
| 12 | 101 | 75 | 103 | 112 | 122 |
| 13 | 72 | 86 | 96 | 84 | 132 |
| 14 | 118 | 118 | 127 | 51 | 46 |
| 15 | 126 | 79 | 96 | 125 | 37 |
| 16 | 99 | 100 | 87 | 134 | 64 |
| 17 | 104 | 108 | 102 | 57 | 100 |
| 18 | 79 | 108 | 118 | 84 | 115 |
| 19 | 87 | 97 | 83 | 114 | 55 |
| 20 | 95 | 100 | 93 | 109 | 93 |
| 21 | 71 | 59 | 102 | 66 | 33 |
| 22 | 107 | 68 | 77 | 88 | 191 |
| 23 | 118 |  | 77 | 183 | 42 |
| 24 | 75 |  | 76 | 59 | 88 |
| 25 | 110 |  | 89 | 101 | 97 |
| 26 | 99 |  | 69 | 91 | 184 |
| 27 | 103 |  | 71 | 106 |  |
| 28 | 108 |  | 82 | 109 |  |
| 29 | 76 |  | 103 |  |  |
| 30 | 92 |  | 67 |  |  |
| 31 | 62 |  | 73 |  |  |
| 32 | 79 |  | 109 |  |  |
| 33 | 91 |  | 57 |  |  |
| 34 | 82 |  |  |  |  |
| 35 | 79 |  |  |  |  |

*tj driven Knockdown Data:*
| **Testis #** | ***Control RNAi*** | ***Set1 RNAi*** |
| --- | --- | --- |
| 1 | 88 | 86 |
| 2 | 78 | 72 |
| 3 | 85 | 103 |
| 4 | 68 | 66 |
| 5 | 57 | 84 |
| 6 | 77 | 73 |
| 7 | 91 | 83 |
| 8 | 81 | 95 |
| 9 | 85 | 75 |
| 10 | 90 | 45 |
| 11 | 69 | 107 |
| 12 | 77 | 105 |
| 13 | 61 | 98 |
| 14 | 65 | 114 |
| 15 | 82 | 98 |
| 16 | 78 | 88 |
| 17 | 102 | 115 |
| 18 | 99 | 97 |
| 19 | 84 | 94 |
| 20 | 67 | 101 |
| 21 | 68 | 127 |
| 22 | 96 | 97 |
| 23 | 97 | 96 |
| 24 | 96 | 104 |
| 25 | 83 | 101 |
| 26 | 100 | 133 |
| 27 | 86 | 111 |
| 28 | 93 | 132 |
| 29 | 69 | 123 |
| 30 | 96 | 103 |
| 31 | 79 |  |
| 32 | 92 |  |
| 33 | 127 |  |

*bam driven Knockdown Data:*
| **Testis #** | ***Control RNAi*** | ***Set1 RNAi*** |
| --- | --- | --- |
| 1 | 94 | 122 |
| 2 | 99 | 150 |
| 3 | 110 | 147 |
| 4 | 98 | 99 |
| 5 | 97 | 158 |
| 6 | 96 | 138 |
| 7 | 100 | 144 |
| 8 | 118 | 146 |
| 9 | 110 | 130 |
| 10 | 86 | 161 |
| 11 | 120 | 155 |
| 12 | 102 | 149 |
| 13 | 101 | 134 |
| 14 | 96 | 122 |
| 15 | 89 | 157 |
| 16 | 130 | 107 |
| 17 | 117 | 142 |
| 18 | 88 | 108 |
| 19 | 115 | 130 |
| 20 | 94 | 146 |
| 21 | 100 | 128 |
| 22 | 81 | 133 |
| 23 | 110 | 138 |
| 24 | 105 | 138 |
| 25 | 136 | 116 |
| 26 | 139 | 141 |
| 27 | 124 | 143 |
| 28 | 64 | 100 |
| 29 | 79 | 105 |
| 30 | 112 | 107 |
| 31 | 111 | 143 |
| 32 |  | 129 |
| 33 |  | 116 |
| 34 |  | 139 |
| 35 |  | 145 |
| 36 |  | 143 |
| 37 |  | 128 |
| 38 |  | 142 |
| 39 |  | 114 |
| 40 |  | 130 |
| 41 |  | 114 |
| 42 |  | 132 |
| 43 |  | 122 |
| 44 |  | 174 |
| 45 |  | 105 |
| 46 |  | 142 |
| 47 |  | 141 |
| 48 |  | 131 |
| 49 |  | 117 |
| 50 |  | 112 |

*nos-gal4, bam-gal80 driven Knockdown Data:*
| **Testis #** | ***Control RNAi*** | ***Set1 RNAi*** |
| --- | --- | --- |
| 1 | 106 | 70 |
| 2 | 71 | 75 |
| 3 | 69 | 74 |
| 4 | 106 | 76 |
| 5 | 72 | 76 |
| 6 | 76 | 97 |
| 7 | 99 | 80 |
| 8 | 77 | 54 |
| 9 | 75 | 94 |
| 10 | 83 | 81 |
| 11 | 88 | 71 |
| 12 | 87 | 70 |
| 13 | 94 | 70 |
| 14 | 89 | 79 |
| 15 | 95 | 85 |
| 16 | 76 | 87 |
| 17 | 76 | 83 |
| 18 | 83 | 73 |
| 19 | 83 | 79 |
| 20 | 71 | 76 |
| 21 | 93 | 84 |
| 22 | 88 | 112 |
| 23 | 85 | 84 |
| 24 | 81 | 66 |
| 25 | 94 | 55 |
| 26 | 80 | 76 |
| 27 | 82 | 90 |
| 28 | 71 | 72 |
| 29 | 66 | 96 |
| 30 | 71 | 84 |
| 31 | 64 | 69 |
| 32 | 97 | 78 |
| 33 | 67 | 81 |
| 34 | 59 | 78 |
| 35 | 81 | 153 |
| 36 | 82 | 201 |
| 37 | 81 | 121 |
| 38 | 76 | 89 |
| 39 | 76 | 94 |
| 40 | 63 | 82 |
| 41 | 74 | 114 |
| 42 | 72 | 63 |
| 43 | 90 | 78 |
| 44 | 96 | 82 |
| 45 | 90 | 77 |
| 46 | 70 | 96 |
| 47 | 81 | 64 |
| 48 | 80 | 69 |
| 49 | 122 | 75 |
| 50 | 100 | 84 |
| 51 | 86 | 87 |
| 52 | 109 | 92 |

